# CSF1R-CAR T cells induce CSF1R signalling and promote cancer cell growth

**DOI:** 10.1101/2024.12.17.629028

**Authors:** Aurora Callahan, Xinyan Zhang, Amber Wang, Aisharja Mojumdar, Longhui Zeng, Xiaolei Su, Arthur R. Salomon

## Abstract

Chimeric antigen receptor (CAR) T cells have transformed the landscape of cancer therapy and demonstrate unprecedented success in treating relapsed/refractory blood cancers. The mechanism underlaying the interactions and responses of CAR T cells and their targets remain incompletely understood. Previous studies focus on the activation of CAR T cells and attempt to optimise CAR design to increase efficacy, meanwhile ignoring tumours and their responses to CAR ligation. Here, we evaluate the signalling capacity of a second generation CSF1-tageted CSF1R CAR compared with a scFv-targeted CD19-CAR using a SILAC co-culture approach coupled with phosphotyrosine (pY) enrichment and LC-MS/MS. We show that ligation of CSF1R-expressing THP1 cells with CSF1R-CAR T cells induces CSF1R-like signalling in THP1 cells, whereas no target cell signalling response is observed after CD19- CAR/Raji B cell ligation. Using small molecule inhibitors of Lck, actin polymerisation, and CSF1R, we find that CAR-induced CSF1R signalling in THP1 cells depends exclusively on CSF1R kinase activity with no participation from T cell activation. Consistently, CSF1R- CAR T cells promote THP1 growth at low effector-to-target (E:T) ratios but prevent THP1 growth at high E:T ratios. Our data provide evidence for an unintended consequence of CARs; CAR-induced signalling in cancer cells. These data may have broad implications for the choice of CAR antigen for optimal clinical efficacy.

**One Sentence Summary:** CSF1R-CAR activates intracellular signalling cascades in THP1 cells, which promote THP1 cell growth.

## INTRODUCTION

The development of cancer immunotherapies including cancer-targeted antibodies, bispecific T cell receptors (TCRs) engagers, and chimeric antigen receptors (CARs) have profoundly improved survival rates among patients with aggressive cancers. Currently, six CARs are FDA-approved for use in treating relapsed or refractory multiple myelomas, B-cell lymphomas, acute lymphoblastic leukemias, and mantle cell lymphomas (*1*). The success and development of CARs is greatly aided by their modular design, as each CAR contains one or more single chain variable fragment (scFv) from an antibody, providing target specificity towards one or more proteins overexpressed on the tumour, a transmembrane domain, and multiple intracellular signalling domains from receptors involved in T cell activation. All currently approved CARs include the intracellular domain of the ζ-chain, a component of the TCR complex involved in TCR signal initiation by acting as a primary T cell activation signal (*2*, *3*). Costimulation, a requirement for full activation and effector function of cytotoxic T cells, is achieved in CAR T cells through the inclusion of intracellular signalling domains from canonical T cell costimulatory receptors such as CD28 or 4-1BB (*4–7*). While only six CARs have been approved, many CARs targeting a diverse array of target-associated antigens (TAAs) are being developed and tested in animal models and patients (*8–10*).

Selecting viable target proteins that reduce off-target effects and retain specificity toward the tumours is necessary for the efficacy of CAR T cell therapy. The approved CARs target cluster of differentiation 19 (CD19) or B cell maturation antigen (BCMA), which are expressed widely on cells in the B cell lineage (*11–13*). While these CARs have been effective in treating multiple myelomas, non-Hodgkin lymphomas, and acute lymphoblastic leukemias, CAR treatment indiscriminately lyses B cell lineage cells expressing target antigens, leading to B cell aplasia (*14*, *15*). Many patients experience antigen escape, whereby CARs eradicate tumours expressing the TAA and inadvertently select for a TAA-negative tumour population or are downregulated by the tumour microenvironment (*16*). Immune evasion is a hallmark of cancer development, which is a limitation of immunotherapies and an active area of research (*17*, *18*).

There has been a wave of research evaluating the use of CAR T cells for the clearance of tumour associated macrophages (TAMs), which reside in and are heavily regulated by tumours to promote downregulation of immune cells that would otherwise respond to the tumour (*19–21*). Therapies targeting TAMs can tremendously improve standard cancer therapeutics (*22*), and are thus an attractive secondary target for CAR T cell therapies. Although few studies have evaluated CAR T cell efficacy against TAMs, two notable studies by Rodriguez-Garcia *et al*. 2021 and Achkova *et al.* 2022 provide different targeting strategies to deplete TAMs or lymphoma lines expressing TAM markers with some success. Rodriguez-Garcia *et al.* show that a subpopulation of TAMs express folate receptor β (FRβ) with an M2-like macrophage phenotype, and FRβ^+^ TAMs downregulate mesothelin-targeting CAR T cells during antigen ligation. Further, by targeting FRβ^+^ TAMs with a third generation scFv-based CAR in conjunction with a third generation CD19 CAR, Rodriguez-Garcia *et al*. observe FRβ+ TAM targeting and depletion, which improves CD19-CAR tumour lysis (*20*). Achkova *et al*. opted to target colony stimulating factor 1 receptor (CSF1R), a receptor tyrosine kinase that drives macrophage growth and differentiation that is highly expressed on TAMs and some cancers, using CSF1 and interleukin 34 (IL-34), two native ligands of CSF1R with different binding affinities. They show that using CSF1 as the targeting domain for CD28-TCRζ or CD28-41BB- TCRζ CARs (28ζ and 28bbζ, respectively) leads to cytokine production and CSF1R-expressing lymphoma line depletion in vitro (*23*). While Rodriguez *et al*. show the potential for targeting and depleting TAMs, Achkova *et al*. show that targeting using native ligands may also be effective, and do so while targeting a protein necessary for macrophage development and persistence.

Considering the high expression of signalling active proteins with known ligands in many cancers and on TAMs, we expect many researchers to evaluate ligand-targeted CARs for tumour and TAM clearance in the future. Here, we evaluate the signalling capacity of a second generation CSF1R targeting CAR constructed using CSF1 as the targeting moiety (CSF1-28tm- 28co-ζ, CSF1R-CAR hereafter) in comparison to a well-established CAR design targeting CD19 (CD19scFv-8tm-28co-ζ, CD19-CAR hereafter). We show that CAR expression is stable in a Jurkat T cell model system, and use liquid chromatography tandem mass spectrometry (LC-MS/MS) to evaluate bi-directional phosphotyrosine (pY) signalling with a stable isotopic labelling of amino acids in cell culture (SILAC) co-culture method our laboratory previously established (*24*). We find that CAR signalling differs between CD19-CAR T cells and CSF1R- CAR T cells, with a higher signalling response observed in CD19-CARs. In contrast, Raji cells show no trend of activation during CD19-CAR co-culture, whereas THP1 cells expressing CSF1R show robust signal induction during CSF1R-CAR co-culture. Small-molecule inhibition of CSF1R ablates the signalling phenotype in THP1 cells, whereas inhibition of Lck or actin polymerisation in CSF1R-CARs does nothing. Finally, we found that CSF1R-CARs can induce THP1 growth at low effector to target (E:T) ratios and can clear THP1 cells at high E:T ratios in vitro. Our data show that the method of targeting influences the propensity for clearance, which should be considered during the design of new CARs.

## RESULTS

### Phosphotyrosine signal induction is altered between CD19-CAR and CSF1R-CAR

To determine whether a ligand-targeted CAR is capable of productively engaging its target receptor, we expressed a second generation CSF1R-targeting CAR that uses CSF1, a native ligand of CSF1R, as the targeting moiety in Jurkat T cells. As a control, we expressed a CD19-targeting CAR modelled after the therapeutic Yescarta (Axicabtagene ciloleucel; Figure 1A, Supporting Figure 1A-B) (*25*). Both CSF1R-CAR and CD19-CAR were expressed on the surface (Figure 1B-C) with high GFP expression in CD19-CAR T cells (Supporting Figure 1C). To evaluate signal induction during CAR-target ligation, we used CSF1R-expressing THP1 cells (Supporting Figure 1D) and CD19-expressing Raji cells in co-culture stimulation experiments using a 2:1 T- to target-cell ratio (E:T - effector to target). We found that CD19-CAR/Raji co-cultures show a characteristic, CAR-dependent T cell signalling-like induction pattern by α-pY (clone 4G10) Western blot (*26*), with statistically significant induction of bands between 50-100 kDa and a high molecular weight band corresponding to PLCγ1 after 2- and 5- minutes of co-culture (Figure 1D, Supporting Figure 2A). In contrast, CSF1R-CAR/THP1 co-cultures showed an altered, not statistically significant banding pattern after 2 minutes of co-culture, with fewer 50-100 kDa pY bands inducing and profound induction of a THP1-specific high molecular weight pY protein (Figure 1E, Supporting Figure 2C). In contrast, non-CAR expressing Jurkat T cells showed a significant reduction in global tyrosine phosphorylation after co-culture with either Raji cells or THP1 cells (Supporting Figure 2B, 2D) indicating the CAR is necessary for any pY changes observed by Western blot. Together, these data suggest there are differences in pY signal induction between CD19-CAR/Raji and CSF1R-CAR/THP1 co-cultures.

**Figure 1:**
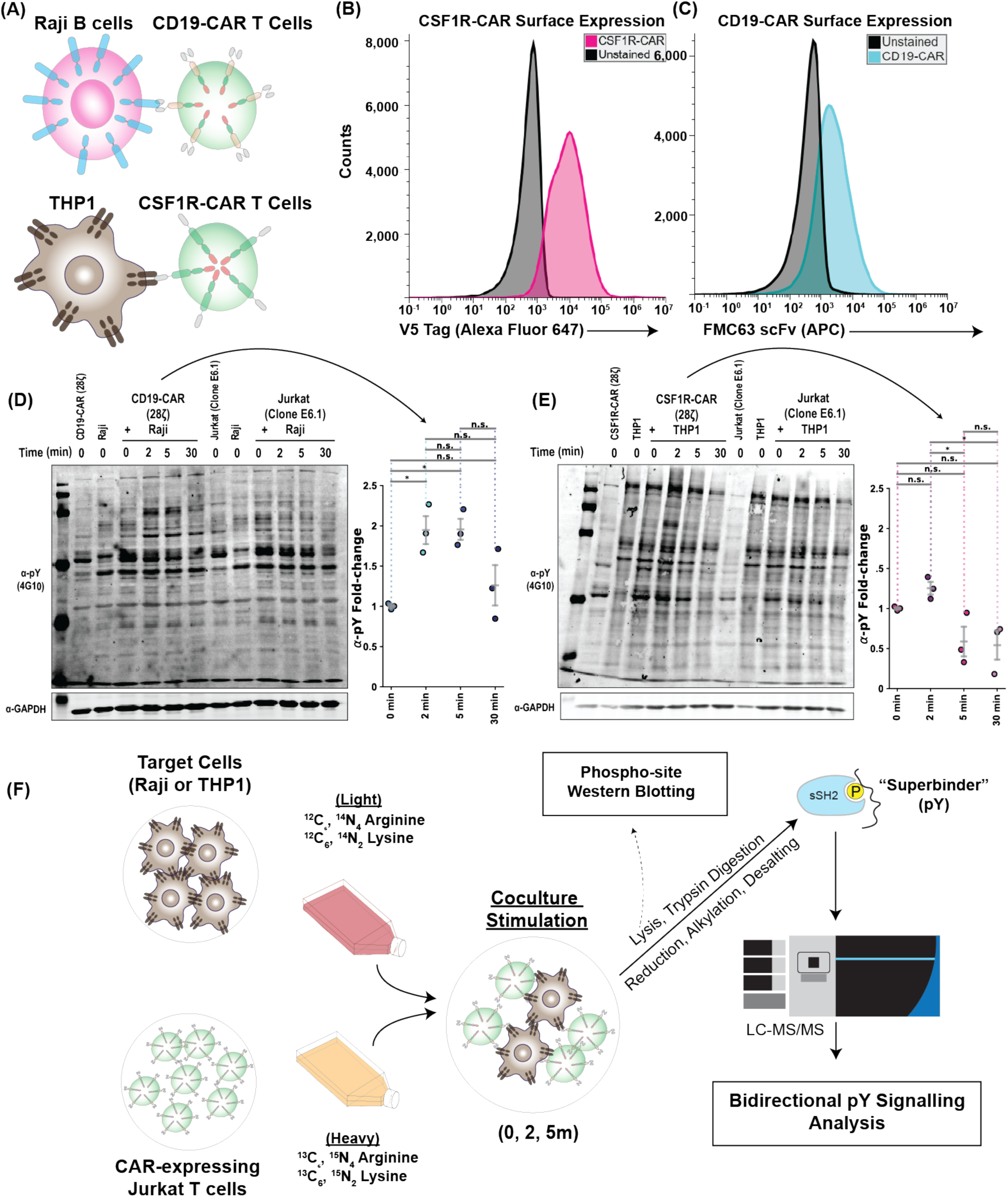
Phosphotyrosine signal induction is altered between CD19-CAR and CSF1R-CAR. **(A)** Schematic of co-culture activation of CD19-CAR and CSF1R-CAR. **(B)** Flow cytometry analysis showing surface expression of CSF1R-CAR (left) andCD19-CAR (right). **(C)** Analysis of tyrosine phosphorylation by Western blot for CD19-CAR/Raji co-culture samples. **(D)** As in (C), except for CSF1R-CAR/THP1 cells. Statistical significance is determined by a two-way ANOVA with Fisher’s least significant difference and the Holm- Sidak correction for multiple comparisons. * : p < 0.05, ** : p<0.01, *** : p<0.001. **(E)** Workflow for SILAC co-culture experiments analysed by mass spectrometry.

**Figure 2:**
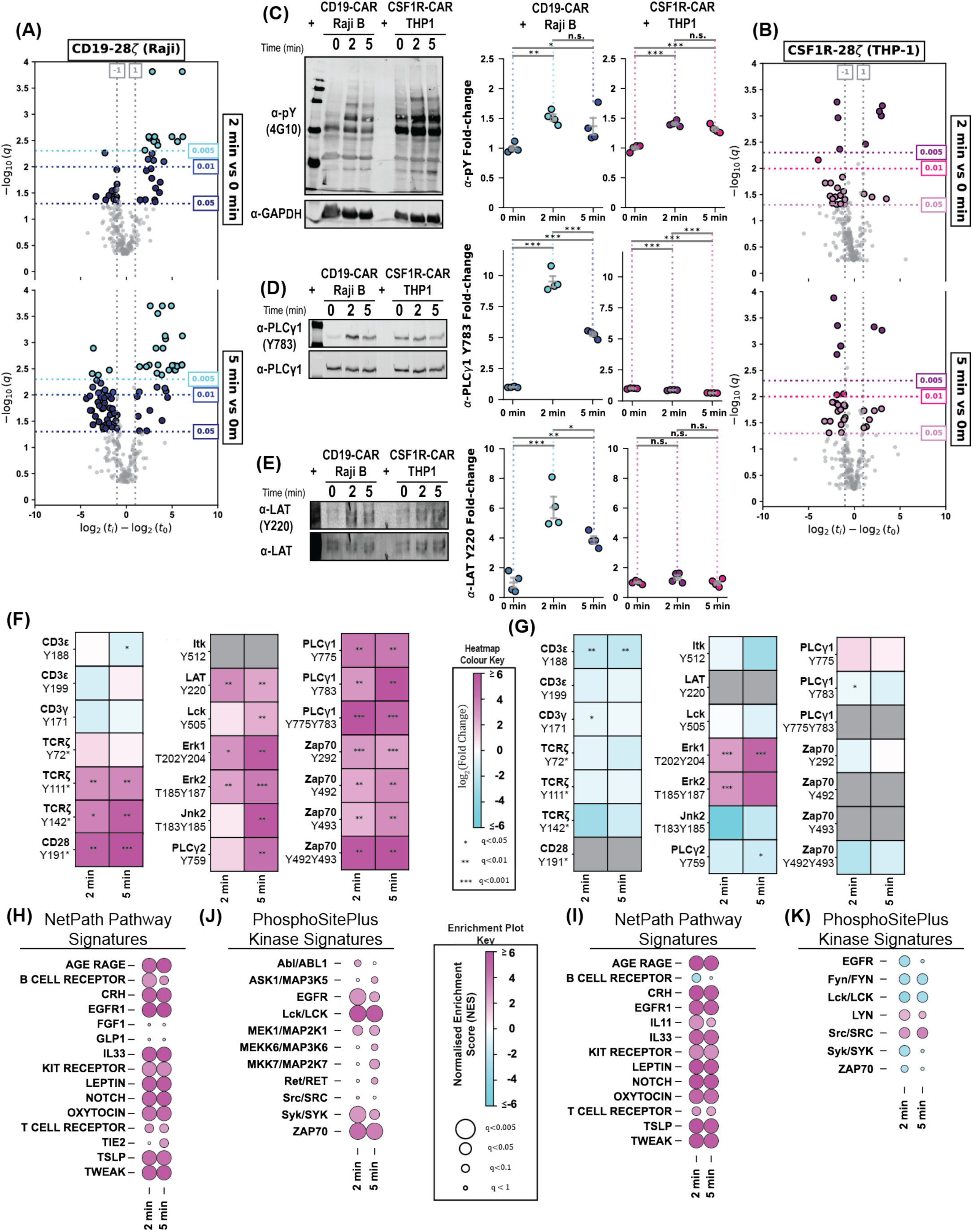
The ligand-targeted CSF1R-CAR minimally engages the T cell receptor signalling pathway. **(A)** Volcano plot analysis of pY sites observed in CD19-CAR T cells during co- culture with Raji cells after 2- and 5-minutes. **(B)** As in (A), except for CSF1R-CAR/THP1 cells. **(C)** Western blot analysis of pY (4G10) abundance in proteomics samples, with GAPDH-normalised quantification to the right. **(D)**, **(E)** as in (C) except for PLCγ1 Tyr^783^ and LAT Tyr^220^ respectively. LAT Tyr^220^ is normalised to PLCγ1 protein due to low LAT protein signal. Statistical significance is determined by a two-way ANOVA with Fisher’s least significant difference and the Holm-Sidak correction for multiple comparisons. * : p < 0.05, ** : p<0.01, *** : p<0.001. **(F)** Heatmaps showing quantification of individual phosphorylation sites observed in CD19-CAR T cells during co-culture with Raji cells. **(G)** As in (F) except for CSF1R-CAR T cells and THP1 cells. **(H)** Post-translational modification signature enrichment analysis (PTM-SEA) showing enrichment of CD19-CAR T cell PTM sites in NetPath Pathways. **(G)** PTM-SEA showing enrichment of CD19-CAR T cell PTM sites in the PhosphoSitePlus Kinase-signatures database. **(I)**, **(J)** As in (H) and (J), respectively, except for CSF1R-CAR T cells.

### CSF1R-CAR minimally engages the T cell receptor signalling pathway

To provide a comprehensive analysis of bi-directional pY signalling during co-culture, we used our previously established SILAC co-culture mass spectrometry-based methodology (Figure 1F) which allows us to use isotopic labelling as a cell identity marker for peptides acquired by mass spectrometry (*24*). In our previous study, we found that an FMC63 scFv-based, third generation CD19-CAR robustly induces phosphorylation of canonical TCR signalling proteins (*24*). Here, we report that our CD19-CAR reproducibly engages the T cell signalling pathway, similarly to that of a parameter matched third generation CAR, with large scale induction of signalling after 2 and 5 minutes of co-culture with Raji cells (Figure 2A, Supporting Figures 3A, 4). In contrast, there was very little pY induction in CSF1R-CAR T cells at either 2 or 5 minutes post co-culture despite reproducible peptide sequencing (Figure 2B). Western blot analysis of pY proteomics lysates revealed increased global tyrosine phosphorylation in both CD19-CAR/Raji and CSF1R-CAR/THP1 co-cultures, although only CD19-CAR/Raji co-cultures showed significant elevation of two key TCR signalling pY sites, PLCγ1 Tyr^783^ and LAT Tyr^220^ (Figures 2C-E, Supporting Figure 5) (*27*, *28*). In CD19- CAR/Raji co-cultures, our mass spectrometry data revealed induction of TCRζ immunoreceptor tyrosine based activation motifs (ITAMs) but not ITAMs on CD3ε, CD3γ, or CD3δ, which are involved in the forward progression of TCR signalling (*29*, *30*). Further, we observed induction of pY sites on proximal T cell signalling proteins, including LAT Tyr^220^, PLCγ1 Tyr^775^, PLCγ1 Tyr^783^, PLCγ1 Tyr^775^Tyr^783^, Zap70 Tyr^492^, Zap70 Tyr^493^, and Zap70 Tyr^429^Tyr^493^, as well as distal TCR proteins like Jnk2 Thr^183^Tyr^185^, Erk1 Thr^202^Tyr^204^, and Erk2 Thr^185^Tyr^187^ (Figure 2F) (*31*). In agreement with our Western blot analysis, CSF1R- CAR T cells displayed little, if any, pY induction on TCR signalling proteins with the exception of the Erk1 Thr^202^Tyr^204^ and Erk2 Thr^185^Tyr^187^, which lay toward the bottom of feedforward CAR signalling (Figure 2G). Using post-translational modification signature enrichment analysis (PTM-SEA), an algorithm which reveals PTM-specific enrichment of signalling pathways, kinases, and drug signatures (*32*), we saw that TCR signalling, and signalling pathways in general, are significantly upregulated in CD19-CAR and CSF1R- CARs after co-culture, despite little significant signal induction in the TCR signalling pathway pY sites in CSF1R-CARs (Figures 2H, 2J). Substrates for the integral TCR kinases Lck and Zap70 were exclusively enriched in the CD19-CAR/Raji co-cultures, whereas CSF1R-CARs showed enrichment of Src substrates (Figures 2I, 2K). Taken together, our results demonstrated that ligand-targeted CSF1R-CARs weakly engage the TCR signalling pathway compared to the therapeutically used CD19-CARs under the experimental conditions tested.

**Figure 3:**
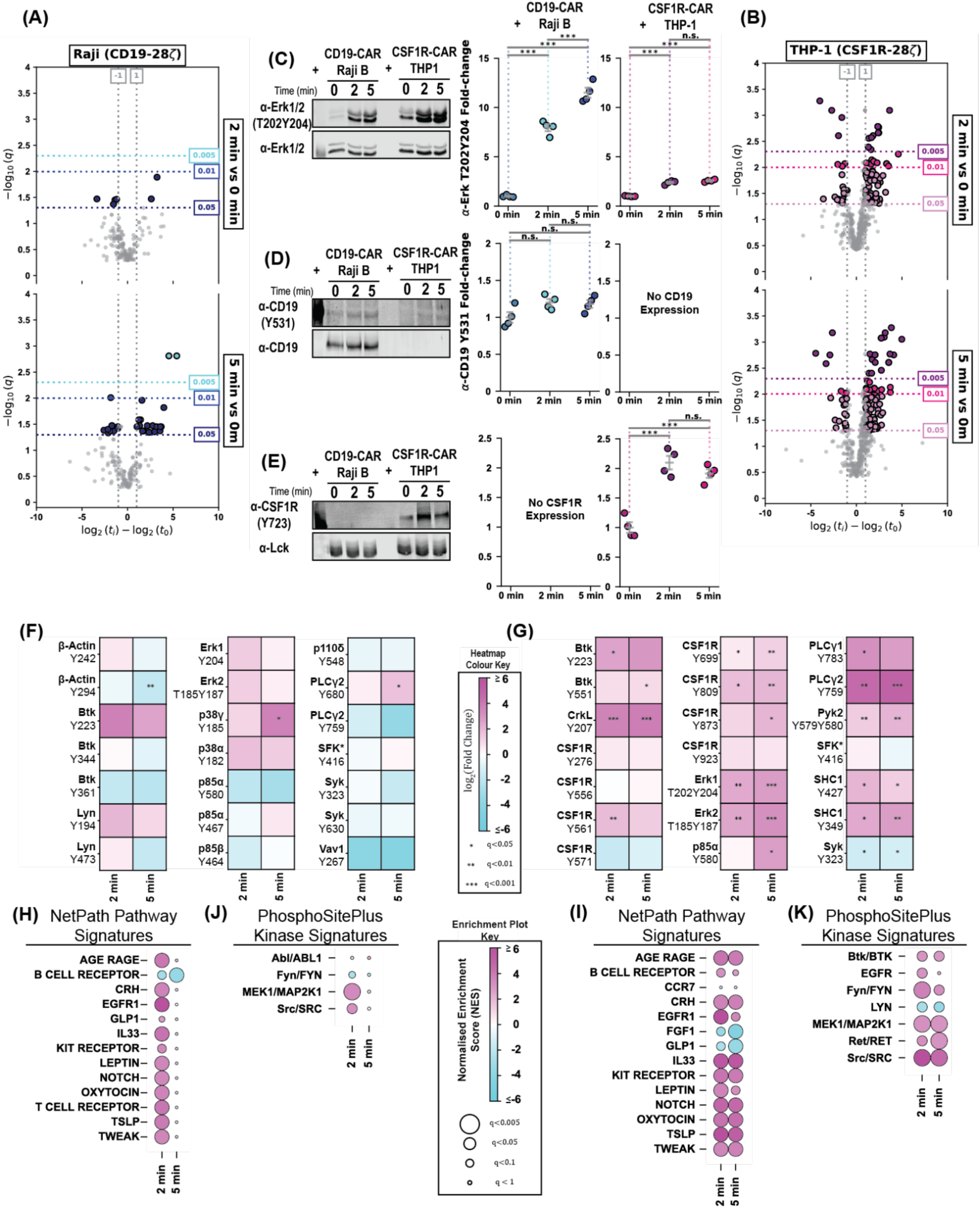
The ligand-targeted CSF1R-CAR activates CSF1R signalling in THP1 cells. **(A)** Volcano plot analysis of pY sites observed in Raji cells during co-culture with CD19-CAR T cells after 2- and 5-minutes. **(B)** As in (A), except for THP1 cells/CSF1R-CAR T cells. **(C)** Western blot analysis of Erk1/2 Thr^202^Tyr^204^ phosphorylation in proteomics samples, with quantification to the right. **(D)**, **(E)** as in (C) except for CD19 Tyr^531^ and CSF1R Tyr^723^ and respectively. CSF1R Tyr^723^ is normalised to total Lck. Statistical significance is determined by a two-way ANOVA with Fisher’s least significant difference and the Holm-Sidak correction for multiple comparisons. * : p < 0.05, ** : p<0.01, *** : p<0.001. **(F)** Heatmaps showing quantification of individual phosphorylation sites observed in Raji cells during co- culture with CD19-CAR T cells. **(G)** As in (F) except for THP1 cells and CSF1R-CAR T cells. **(H)** Post-translational modification signature enrichment analysis (PTM-SEA) showing enrichment of Raji PTM sites in NetPath Pathways. **(G)** PTM-SEA showing enrichment of Raji PTM sites in the PhosphoSitePlus Kinase-signatures database. **(I)**, **(J)** As in (H) and (J), respectively, except for THP1 cells.

### CSF1R-CAR activates CSF1R signalling in THP1 cells

Ligand-based targeting systems benefit from native receptor-ligand interactions that provide specific, efficacious targeting, as well as higher surface expression compared with scFv designs (Figure 1B Supporting Figure 1C) (*33–35*). Using our SILAC co-culture methodology (Figure 1G), we gain precise information about pY induction in the target Raji or THP1 cells during CAR co-cultures. We observed no clear trend in pY induction in Raji cells after second generation CD19-CAR co-culture, in agreement with our results for third generation CD19-CARs (Figure 3A, Supporting Figures 3C, 6) (*24*). In contrast, pY sites in THP1 cells showed patterns consistent with effective CSF1-CSF1R engagement (Figure 3B, Supporting Figure 3D). By Western blot, we showed elevated phosphorylation of Erk1

Thr^202^Tyr^204^ and Erk2 Thr^185^Tyr^187^ in both CD19-CAR/Raji and CSF1R-CAR/THP1 co-cultures, with a much larger magnitude in CD19-CAR/Raji co-cultures that likely originates from CD19-CAR signalling (Figures 2F, 3C, 3F, Supporting Figures 7A-B). We did not observe phosphorylation of CD19 Tyr^531^ in CD19-CAR/Raji co-cultures, however we observed significant phosphorylation of CSF1R Tyr^723^, a phosphorylation site that promotes binding of phosphoinositide-3-kinase components (*36*), in CSF1R-CAR/THP1 co-cultures (Figures 3D-E, Supporting Figures 7C-F). Site-specific analysis of our CD19-CAR/Raji pY proteomics data showed few, if any, significant changes in B cell signalling related phosphorylation sites in Raji cells, particularly on kinases involved in forward-progression of the signalling cascade (Figure 3F). In contrast, CSF1R-CAR/THP1 co-cultures induced phosphorylation of many proteins involved in CSF1R signalling, including CSF1R Tyr^561^, CSF1R Tyr^699^, CSF1R Tyr^809^, and CSF1R Tyr^873^ showing significant elevation at 2- and/or 5- minutes post co-culture (Figure 3G). CSF1R signalling is not well characterised and is heavily cell dependent (*37–40*), however we observed pY induction on various proteins thought to be involved in CSF1R signalling in macrophage lineage cells, including Pyk2 Tyr^573^Tyr^580^ which regulates its binding capacity (*41*), SHC1 Tyr^349^ which regulates GRB2 recruitment (*42*), the activating sites PLCγ2 Tyr^753^ and PLCγ2 Tyr^759^ (*43*), and the activation sites Erk1 Thr^202^Tyr^204^ and Erk2 Thr^185^Tyr^187^ (*44*, *45*). There was no significant induction of the activating pY site SFK Tyr^416^ (conserved sequence LIEDNEYTAR for Src family kinases) (*46*), with a significant reduction in the negative regulatory site Syk Tyr^323^ (*47*, *48*). Similarly, specific pY sites on CSF1R did not show significant changes after co-culture, including CSF1R Tyr^276^, CSF1R Tyr^556^, CSF1R Tyr^571^, and CSF1R Tyr^923^ (Figure 3G). PTM-SEA revealed that, despite few significant changes in pY induction, many signalling pathway signatures but only kinase signatures of MEK1 and Src are upregulated in Raji cells after 2 minutes of co-culture (Figures 3H-I). In THP1 cells, pathway signatures and kinase signatures for Btk, MEK1, and Src, which are integral for the progression of CSF1R proximal and distal signalling, were upregulated after 2 and 5 minutes of co-culture. Taken together, our data demonstrate that a ligand-targeted CSF1R-CAR is capable of activating pY signalling in CSF1R-expressing THP1 cells.

### CSF1R-CAR induced CSF1R signalling requires CSF1R kinase activity, not T cell activation or immune synapse formation

During TCR- and CAR-based T cell activation, actin polymerisation is induced which pushes signalling proteins into a small space in the cell-to-cell junction, termed the immune synapse. While CARs are less effective in forming an immune synapse, previous studies suggest that immune synapse formation is still involved in CAR-based T cell activation (*49*, *50*), which could promote or enhance activation of CSF1R during CSF1R-CAR/THP1 co-culture. To uncouple T cell activation and CSF1R signalling in THP1 cells, we inhibited Lck, the activating kinase for T cell signalling (*51*, *52*), with PP1 and actin polymerisation with Cytochalasin D in CSF1R-CAR T cells (Figure 4A) (*53–56*). PP1 pretreatment of CSF1R- CAR T cells at 0.2 μM or 2 μM for 3 hours prior to co-culture did not affect induction of CSF1R Tyr^723^ by Western blot, however 20 μM of PP1 reduced CSF1R Tyr^723^ levels back to the no treatment, no stimulation control (Figure 4B-C). Similarly, treatment of CSF1R-CAR T cells with Cytochalasin D at 0.2 μM, 2 μM, and 20 μM showed no influence on CSF1R Tyr^723^ induction after co-culture (Figure 4D-E). Notably, PP1 pretreatment significantly reduced Erk1 Thr^202^Tyr^204^/Erk2 Thr^185^Tyr^187^ at 2 μM and 20 μM, whereas Erk1 Thr^202^Tyr^204^/Erk2 Thr^185^Tyr^187^ phosphorylation was low and similarly responsive at all Cytochalasin D concentrations (Supporting Figures 8A-D). Cytochalasin D is well known to disrupt actin polymerisation at concentrations greater than 1 μM with a 30 minute pre-treatment without significantly altering Erk activation (*53*, *55*, *56*). To uncouple CSF1R-CAR/CSF1R ligation from CSF1R kinase activity, we used Vimseltinib (*57*), Pexidartinib (*58*), and PLX5622 (*59*), three small molecule inhibitors of CSF1R (Figure 4F). At all tested concentrations (0.2 μM, 2 μM, and 20 μM), inhibition of CSF1R significantly reduced phosphorylation of CSF1RY723 compared with a no-treatment control (Figure 4 G-L).

**Figure 4:**
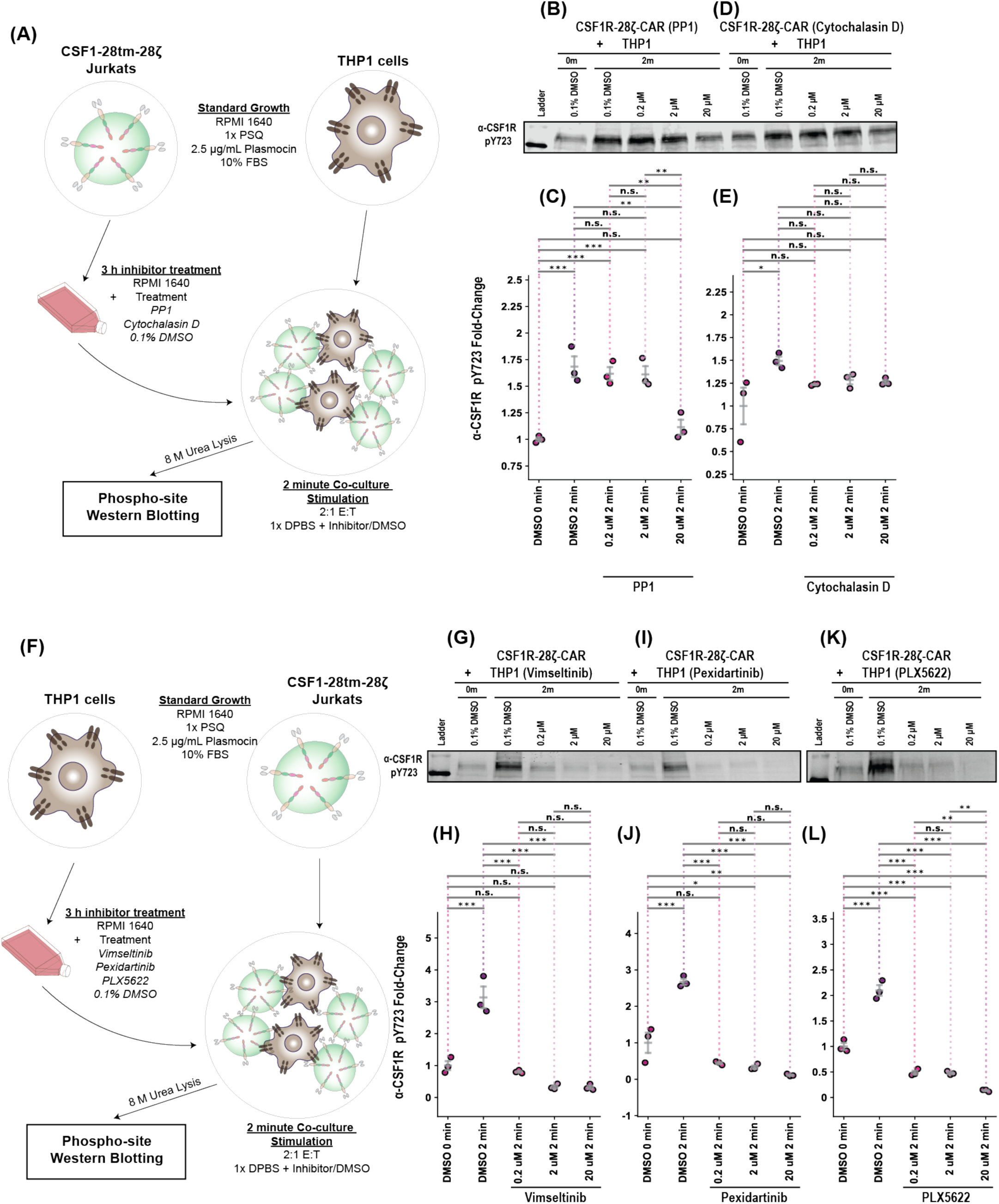
CSF1R signalling requires CSF1R kinase activity, not T cell activation or immune synapse formation. **(A)** Schematic depicting PP1 (Lck inhibitor) and Cytochalasin D (actin polymerisation inhibitor) treatment of CSF1R-CAR T cells before co-culture. **(B)** and **(C)** Western blot analysis of CSF1R Tyr^723^ after PP1 treatment of CSF1R-CAR T cells and quantification (n=3), respectively. **(D)** and **(E)** As in (B) and (C) except for Cytochalasin D treatment. **(F)** Schematic depicting Vimseltinib, Pexidartinib, and PLX 5622 (CSF1R kinase inhibitors) treatment of THP1 cells before co-culture. **(G)** and **(H)** Western blot analysis of CSF1R Tyr^723^ after Vimseltinib treatment of CSF1R-CAR T cells and quantification (n=3), respectively. **(I)** and **(J)** As in (G) and (H) except for Pexidartinib treatment. **(K)** and **(L)** As in (G) and (H) except for PLX 5622 treatment. CSF1R Tyr^723^ is normalised to a ∼100 kDa off target band due to low CSF1R protein signal (Supporting Figure 8). Statistical significance is determined by a two-way ANOVA with Fisher’s least significant difference and the Holm- Sidak correction for multiple comparisons. * : p < 0.05, ** : p<0.01, *** : p<0.001.

**Figure 5:**
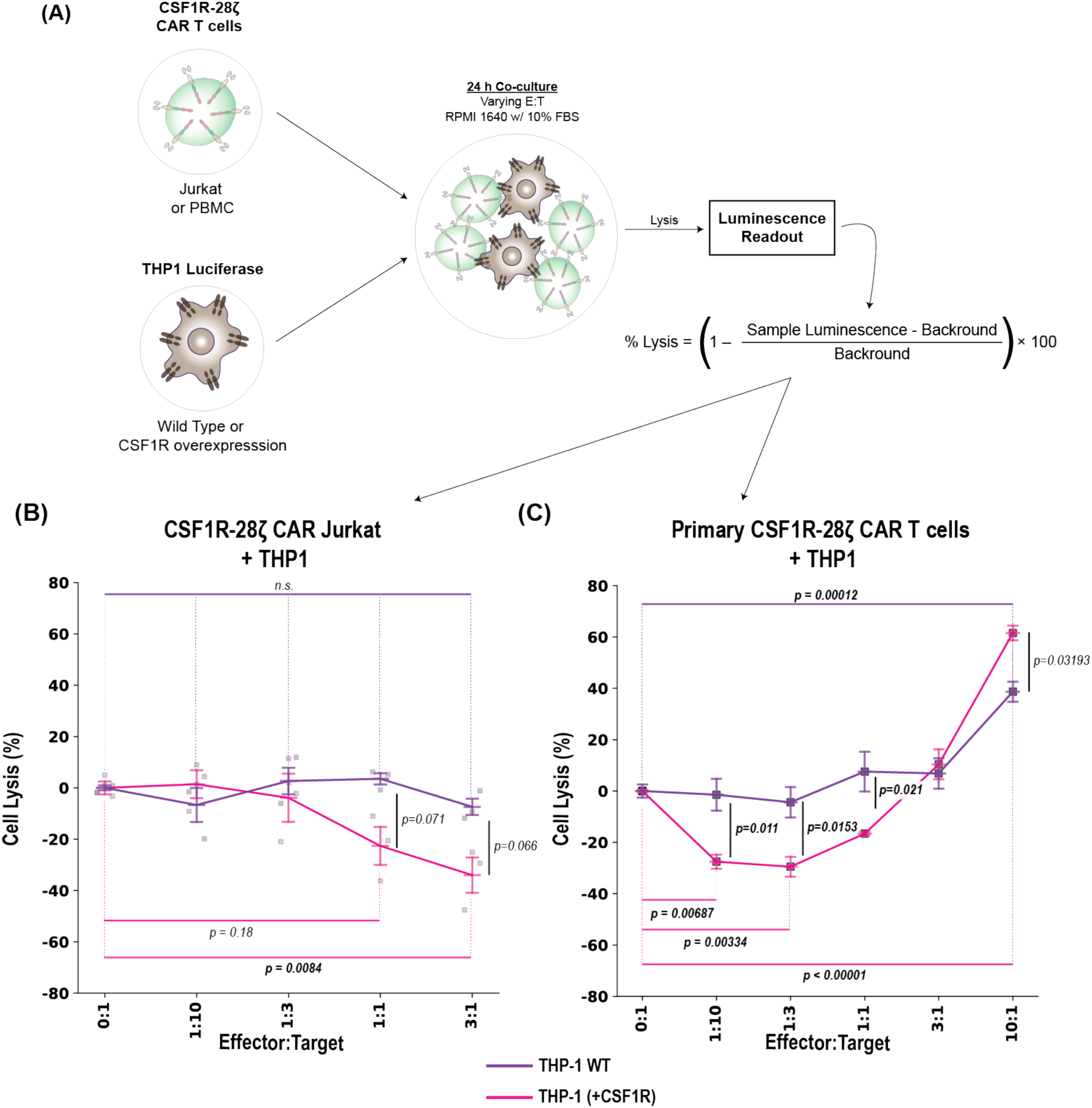
Dual effects of CSF1R CAR T cells on the growth of THP1 cells. (A) Schematic depicting co-culture growth assays. (B) some growth assay stuff (C) Model depicting pY signalling from (left to right) CD19-CAR T cells, CSF1R-CAR T cells, Raji cells, and THP1 cells. (D) Model depicting the effect of CAR T cells on target cells. p-values determined by a one-way ANOVA with Fisher’s LSD and the Holm-Sidak correction for mean percent lysis comparisons.

PLX5622 in particular reduced CSF1R Tyr^723^ abundance below the basal state at all tested concentrations (Figure 4I-J). We saw that inhibiting CSF1R kinase activity also reduced Erk1 Thr^202^Tyr^204^/Erk2 Thr^185^Tyr^187^ abundance significantly, usually reducing induction from around 4- to 6-fold above basal without treatment to 2-3 fold with treatment (Supporting Figure 8E-J), which was in agreement with our proteomics data showing significant induction of Erk1 Thr^202^Tyr^204^/Erk2 Thr^185^Tyr^187^ in THP1 cells and less induction in CSF1R-CAR T cells (Figures 2G, 3C, 3G). Taken together, our results demonstrate that the kinase activity of CSF1R is the primary determinant of signalling in THP1 cells during CSF1R-CAR/THP1 co-culture, with Lck activity in CSF1R-CAR T cells playing a minimal role.

### Dual effects of CSF1R-CAR T cells on the growth of THP1 cells

Considering that CSF1R is known to control macrophage growth (*60*), we sought to determine whether CSF1-targeted CARs are able to induce growth of targeted THP1 cells under our experimental conditions. To determine THP1 monocyte number during co-culture with CSF1R-CAR T cells, we used THP1 cells expressing a luciferase reporter and calculate the percent lysis compared with a control without T cells (Figure 5A) (*61*). Co-culture of CSF1R-CAR expressing Jurkat T cells (Supporting Figure 9, top), which have no lytic activity, with THP1 cells overexpressing CSF1R (Supporting Figure 1D) showed a dose-dependent effect on THP1 cell growth. As a control, native THP1 cells expressing low levels of CSF1R showed no significant change in percent cell lysis compared to a no-CAR control (Figure 5B). Next, we tested the effect of primary human T cells expressing CAR, which contain cytolytic activity, on the growth of THP1 cells. Co-culture of CSF1R-CAR expressing T cells isolated from healthy donors (Supporting Figure 9, middle) with CSF1R overexpressing THP1 cells promoted cell growth at E:T ratios of 1:10 (p=0.00687) and 1:3 (p=0.00334), while significantly inducing cell lysis at an E:T ratio of 10:1 (p<0.00001) when compared to a no-CAR control (Figure 5C). In contrast, co-culture of primary CSF1R-CAR T cells with native THP1 cells only induced lysis at an E:T ratio of 10:1 (p=0.00012) although less than CSF1R overexpressing THP1 cells (p=0.03193). These data suggest that CSF1R- CAR T cells display a dual effect on target cells: they promote THP1 cell growth at low E:T ratios but inhibit THP1 cell growth at high E:T ratios.

## DISCUSSION

The boom of chimeric antigen receptor T cell therapy has revolutionised targeted cancer immunotherapies and is currently under investigation for use in autoimmune disorder and fungal infection treatment and in various non-T immune cells (*62–65*). Despite the clinical success of second generation CARs for treating relapsed or refractory B cell acute lymphoblastic leukaemia, B cell lymphoma, and multiple myelomas, many CARs fail in clinical trials (*66*, *67*) and patients suffer from a number of side effects including off-target activation, cytokine release syndrome, and neurotoxicity, which can vary severely between patients (*68*, *69*). Side effects of CAR T cell therapy can be due to CAR design which have been ameliorated in newer generations and designs or off-target CAR-target ligation (*70*, *71*). The current age of CAR design focuses heavily on rational design of new CARs, leading to the creation of new targeting moieties (*72*) and receptor designs (*73–75*), scFv screening (*76*– *80*), costimulatory domains, primary stimulation domains (*81*, *82*), and cytotoxicity (*83*, *84*), which has improved the discovery of new, effective CARs. In particular, the therapeutically approved designs are well studied, with mechanistic studies characterising CAR signalling and using these insights to improve CAR design (*85–92*). The majority of the literature, however, focuses on modulating the activation timing and strength of CARs, and little is known regarding how and if CAR engagement promotes responses in the targeted cells.

In a recent study, Zhong *et al*. (2023) evaluate the role of tumour-derived small extracellular vesicles (sEV) in CAR T cell resistance, showing that sEV excreted by tumours carrying large amounts of CAR-targeted antigens and PD-L1. These sEV act as both decoys, preferentially binding CAR T cells rather than the targeted tumour, and negative regulators of CAR T cell function, owing to the high abundance of the target antigen Mesothelin and PD- L1 (*93*). In their study, Zhong *et al*. use an scFv-based CAR targeting Mesothelin, a transmembrane protein overexpressed in many cancers that sits atop RAS/MAPK, JAK/STAT, and AKT/mTOR signalling pathways (*94*). sEV production by tumours is found to correlate with tumour/CAR T cell interactions, suggesting that CAR T cells are able to induce sEV excretion from the pancreatic ductal adenocarcinomas (PDAC) used by Zhong *et al*. While Zhong *et al*. do not provide a precise mechanism for sEV production by PDAC in response to CAR T cell treatment, they provide ample evidence of CAR-specific tumour responses. While our CAR construction and targets are different, our study provides a similar story: CARs can induce specific, unexpected responses from tumours. We show that using CSF1, a native ligand of CSF1R, as the targeting moiety for second generation CAR T cells is able to produce a signalling response in THP1 cells expressing CSF1R (Figures 1 & 3, Supporting Figures 2, 7). This is in stark contrast to CD19-targeted CARs, which consistently produce no response in Raji cells expressing CD19 irrespective of CAR generation (Figures 1 & 3, Supporting Figures 2, 5) (*24*). The observed signalling response requires the kinase activity of CSF1R (Figure 4F-L), which is required for the many functions of CSF1R (*95*), with minimal input from the activity of CAR T cells (Figure 4A-E) Further, we find that CARs can induce growth of THP1 cells in vitro when the effector-to-target stoichiometry favours the target (Figure 5). Together, our results with those of Zhong *et al*. provide an interesting consequence of CAR T cell therapy, the activity of target cells, which should be screened during the development of new CAR/target pairs to avoid unintended outcomes.

Due to the current limited clinical usage of CAR T cells (*96*), expanding the specificity and retention of CAR T cells is a current priority in the field (*97*). Many researchers have proposed and researched the use of ligand-targeted CARs, those containing a native ligand for a TAA highly expressed on the targeted tumour, with results that appear favourable overall. A study by Chinsuwan *et al*. (2023) shows that EGF- and TGFα-CARs targeting EGFR can be expressed stably and can clear a number of non-small cell lung cancer (NSCLC) cell lines in vitro, although in vitro killing efficiency is reduced in some cases when target cells are more abundant than effector T cells. Chinsuwan *et al*. (2023) do show, however, that the EGF- CARs are able to induce clearance in lympho-depleted mouse models at clinically relevant doses, suggesting that some ligand-targeted CARs can overcome the low effector to target ratios in vivo (*33*). In another study, Achkova *et al*. (2022) propose two CSF1R directed CARs using CSF1 and IL-34 as targeting moieties and show that CSF1-targeted CARs have cytolytic activity and can clear lymphoma cell lines expressing CSF1R in vitro to a greater degree than IL-34-targeted CARs (*23*). As with many CAR T cell studies, both Chinsuwan *et a*l. (2023) and Achkova *et al*. (2022) assume the target cells are static during treatment, and do not investigate any possible mechanisms by which ligand-targeted CARs differentially activate or target cells participate in phenotypes such as cytokine production. In our study, we find that CAR activation at the level of pY signalling varies greatly with the choice of target and binding mechanism (Figure 2). We also show that CSF1R signalling can be induced by CARs and that THP1 cells can be induced to grow by CARs in an effector-to-target- and CSF1R expression-dependent manner (Figures 3-5). These previously unexplored findings suggest that the monolithic view of CAR optimisation through T cell specific modifications may require reworking to include the activity and role of the target cells.

The use of CAR T cells is expanding rapidly, with researchers evaluating CAR T cells for use in treating B cell-based autoimmune disorders, CAR expression in different immune cell types (CAR-natural killer T cells, CAR macrophages), and fungal infections (*62–64*, *73*). As promising new therapeutic routes arise, evaluating both the treatment efficacy and off target effects of CAR T cell therapies is vital to ensuring the safety of patients. In this study, we use THP1 cells as a model macrophage lineage cell to evaluate the influence of a CSF1-based CSF1R-CAR on their intracellular signalling and growth. We find that CARs using CSF1 as the targeting moiety can effectively engage CSF1R and induce CSF1R signalling, leading to in vitro THP1 growth in an effector to target ratio and CSF1R expression dependent manner. Our data suggest that CAR-dependent effects on targeted cells should be evaluated at an organismal and clinical level to ensure safe and effective treatment.

## MATERIALS AND METHODS

### Cell culture, CAR expression, and inhibitor treatment

Jurkat T cells (clone E6.1, ATCC #TIB-152), Raji cells (ATCC #CCL-86) and THP1 cells (ATCC #TIB-202), were cultured in RPMI 1640 supplemented with 10% FBS (PeakSerum #PS-FB3), 1X penicillin-streptomycin-glutamine (PSQ; HyClone #SV30082.01), 2.5 μg/mL Plasmocin (InvivoGen #ant-mpp) at 37 C, 5% CO2 in a humidified incubator. For co-culture mass spectrometry, Jurkat T cells were grown to confluence, washed twice in 1X DPBS, then expanded in SILAC RPMI (Thermo #) supplemented with 0.38 mM ^13^C_6_ ^15^N_4_ Arginine (Millipore #), 0.22 mM ^13^C_6_ ^15^N_2_ Lysine (Cambridge Isotopes #), 10% dialysed FBS, 1X PSQ, and 2.5 μg/mL Plasmocin for 7 days. Primary human T cells were isolated from PBMCs of healthy donors (Zenbio #SER-PBMC-200) using the EasySep Human T cell Isolation Kit (StemCell #17951), cultured in RPMI 1640 supplemented with 10% FBS, 50 nM β- mercaptoethanol, 300 U/mL IL-2 (PEPROTECH #200-02), and treated with α-CD3/α-CD28 dynabeads (Thermo #11161D) for expansion.

For CAR T cell production, HEK293T cells were cultured in DMEM supplemented with 10% FBS and 1X PSQ. One million HEK 293T cells were plated in one well of a six well plate before transfection with pMD2.G, poPAX, and CAR plasmids (0.5 μg each) using PEI MAX. After 10 hours, the culture medium was replaced with primary T cell or Jurkat T cell culture medium, and allowed to incubate for 24 hours, twice, to collect viral particles. Jurkat T cells were cultured with 1.5 mL of viral media per 0.5 million Jurkats, with half of the media replaced with fresh cell culture media after 24 hours. Jurkats were allowed to expand before use. Primary human T cells were infected with 1.5 mL of fresh lentivirus per 0.5 million cells via spinoculation at 800 xg for 90 minutes at 32 C, and half of the culture media was replaced with fresh culture media after 24 hours. Five days after infection, dynabeads were removed and CARs were grown in fresh media for two more days before use.

For T cell inhibitor treatment experiments, Jurkat CSF1R-CAR T cells were collected by centrifugation, washed 1 time with unsupplemented RPMI 1640, then suspended RPMI 1640 supplemented with PP1 (Cayman Chemical Company #14244), Cytochalasin D (Cayman Chemical Company #11330), or DMSO (Fisher Scientific #BP231-100) at a concentration of *1* × *10 ^7^* for 3 hours at 37 C, 5% CO_2_ in a humidified incubator. For THP1 inhibitor experiments, THP1 cells were processed the same as Jurkat CSF1R-CAR T cells and resuspended in RPMI 1640 supplemented with Pexidartinib (Cayman Chemical Company #18271), Vimseltinib (Cayman Chemical Company #38783), PLX 5622 (Cayman Chemical Company #28927), or DMSO. After the treatment, cells were washed 1 time in warm DPBS before co-culture stimulation as described below.

### Co-culture stimulation and mass spectrometry sample preparation

Co-culture stimulations were performed as previously described(*24*). Briefly, CAR T cells and target cells (Rajis, THP1s) were grown to confluence and collected by centrifugation. For Western blotting analysis, 10 million T cells (200 million cells/mL) and 5 million target cells (100 million cells/mL) per replicate were used, and for mass spectrometry experiments 100 million T cells (200 million cells/mL) and 50 million target cells (100 million cells/mL) were used. To promote cell-to-cell contact, mixed cells were immediately centrifuged at 600 xg for 30 seconds. Stimulation was halted by the addition of 9 M Urea Lysis Buffer (9 M Urea, 1 mM sodium orthovanadate, 1 mM sodium pyrophosphate, 1 mM β-glycerophosphate in 20 mM HEPES) to a final concentration of 4 M Urea at the indicated time points. Lysates were chilled on ice for ∼15 minutes, sonicated at 70% amplitude for 30 seconds once, and centrifuged at 1800 xg for 15 minutes to collect the insoluble fraction. Samples for Western blotting were diluted 1:1 in 2X Lamelli sample buffer (4% SDS, 125 mM TRIS-HCl pH 6.8, 20% glycerol, 5% β-mercaptoethanol, 0.01% bromophenol blue), boiled at 95 C for 5 minutes, then frozen until use. For mass spectrometry, protein concentrations were evaluated by BCA (Thermo #23225) and each sample was normalised to 9 mg of total protein. Samples were then treated with DTT for 30 minutes at 30 C (final concentration 10 mM) and iodoacetamide for 15 minutes at room temperature in the dark (final concentration 10 mM) before overnight trypsin digestion (1:100 trypsin:protein, Promega #V5113). The following day, samples were acidified to 1% trifluoroacetic acid (TFA, v/v) and centrifuged at 1800 xg for 15 minutes to remove the insoluble fraction. Peptides were desalted using Sep-Pak C18 plus cartridges (Waters #WAT020515) and lyophilised as previously described(*24*, *98*).

### Phosphotyrosine enrichment with the Src SH2 superbinder

Enrichment of pY peptides was performed using the Src SH2 superbinder (sSH2) domain as previously described(*24*, *99–101*). Briefly, in-house purified recombinant sSH2 protein was conjugated to CNBr-activated sepharose beads at a concentration of 3.2 mg sSH2/mL. Nine milligrams of total desalted peptide was resuspended in 1.4 mL of IAP buffer (10 mM Sodium phosphate monobasic monohydrate, 50 mM sodium chloride, 500 mM MOPS, pH 7.2) and incubated in 100 μg of sSH2 beads for 2 hours at 4 C on a 4 rpm end over end rotator. pY peptide-bound beads were washed 3 times in ice cold HPLC water, 2 times in ice cold IAP buffer, and eluted from beads in 0.15% TFA on a 1150 rpm mixer at room temperature, twice. The eluted pY peptides were desalting using Pierce C18 tips (Thermo #87784) as previously described and dried by speed-vac before LC-MS/MS analysis.

### LC-MS/MS, database searching, and analysis

After enrichment, pY peptides were separated on a lab-packed C18 column (250 mm by 75 μm) with XSelect CSH C18 2 μm resin using an Agilent 1200 HPLC. Peptides were eluted using a 120 minute method starting with a Buffer A (0.1 M acetic acid in HPLC water) to Buffer B (0.1 M acetic acid in acetonitrile) ratio of 95:5 and progressing to a final ratio of 70:30 in 95 min, with a 100% buffer B spike at the end of the gradient. Peptides were analysed on a QExactive mass spectrometer in positive ion mode with a spray voltage of 2.0kV, funnel RF level of 40, heated capillary temperature of 250 C. Precursor mass range was set to 400-1800 m/z with a resolution of 70,000, AGC target of 3e6, IT of 200 ms in centroid mode. Fragmentation spectra were collected using a Top9 DDA method with a dynamic exclusion time of 30 seconds, and an intensity threshold of 1e3. Monoisotopic precursor ions with charge states between +2 and +5 were fragmented using higher-energy collision induced dissociation (HCD) using a normalised collision energy of 28%.

Fragmentation spectra were acquired using a 2.5 m/z isolation window at a resolution of 17,500 and an AGC target of 2e4 in centroid mode with a maximum injection time of 200 ms.

Thermo .RAW files were analysed using the high throughput autonomous proteomics pipeline (HTAPP) and PeptideDepot, custom in-house software for the automation of PTM proteomics analysis, as previously described(*24*, *102*, *103*). Briefly, MS2 spectra were searched using Mascot against the UniProt complete proteome dataset (98300 nonredundant forward sequences, downloaded August 2019) to 0.1% FDR using a reverse decoy database. The following parameters were used for the search: trypsin enzyme cleavage (up to 2 missed cleavages), 7 ppm precursor mass tolerance, 20 mmu fragment ion mass tolerance, variable modifications for phosphorylation (S/T/Y; +79.9963 Da), oxidation (M; +15.9949 Da), and static modification of either carbamamidomethylation (C; +57.0215 Da) for the analysis of THP1 cells or carbamamidomethylation (C; +57.0215 Da) and SILAC labelling (K: +8.0142 Da, R: +10.0083 Da) for the analysis of CAR-T cells. Retention time alignment and integration of selected ion chromatograms for each peptide spectrum match was used to determine relative peptide abundance as previously described(*104*). Initial mass spectrometry data analysis was performed using R in PeptideDepot, with follow up analysis using Python 3.12.5. Briefly, comparisons were built using PeptideDepot in which all peptides were filtered to include only unique pY peptides, student’s T tests were performed comparing the mean abundance of log_2_ transformed 0-, 2-, and 5-minute stimulation conditions, and the global rate of false discovery was adjusted using the method of Storey(*105–107*). Replicate reproducibility was evaluated using principal component analysis (PCA) and multiple regression. Post-translational modification signature enrichment analysis (PTM-SEA) was performed as previously described(*32*) using the R-script ssGSEA2.0.R and the PTM-signature database (version 2.0.0).

### Western blotting

Western blotting samples were separated on 8% or 10% polyacrylamide gels poured in house at 100 V for 130 minutes before transferring to a PVDF membrane (EMD Millipore #IPFL00010) at 100 V for 100 minutes. Membranes were blocked using Intercept TBS blocking buffer (LI-COR #927-60001) diluted 1:1 in 1X TBS (137 mM NaCl, 20 mM TRIS) for 1 hour at room temperature, before overnight incubation in primary antibodies in 5% BSA in 1X TBS with Tween 20 (TBST). After primary antibody incubation, membranes were washed 3 times with 1X TBST for 5 minutes, incubated in secondary antibodies in 5% BSA in 1X TBST, washed 3 times in 1X TBST for 5 minutes, and 2 times in 1X TBS. Membranes were imaged on the LI-COR Odyssey CLx or LI-COR Odyssey M imaging systems.

Quantification of Western blots was performed as previously described(*24*, *108*). Statistical significance between loading-control corrected, 0-minute normalised band intensities was determined using FWER corrected p-values generated using Fisher’s least significant difference test(*109*, *110*) followed by the Holm-Sidak FWER step-up correction(*111*, *112*).

Primary antibodies: pY clone 4G10 (1:2000, Cell Signalling Technologies #96215), GAPDH (1:10000, Millipore Sigma #G9545), PLCγ1 (1:10000, Millipore Sigma #05-163), PLCγ1 Tyr^783^ (1:2000, Cell Signalling Technologies #2821), LAT (only used for identifying LAT bands. 1:2000, Thermo #MA1-19307. PLCγ1 protein was used as the loading control), LAT Tyr^220^ (1:2000, Cell Signalling Technologies #3584), Erk1/2 (Cell Signalling Technologies #9107), Erk Thr^202^Tyr^204^ (1:2000, Cell Signalling Technologies #9101), CD19 (1:2000, Thermo #14-0190-82), CD19 Tyr^531^ (1:2000, Cell Signalling Technologies #3571), CSF1R (1:2000, Thermo #MA5-37676. Off-target band or Lck protein used for normalisation in quantification), CSF1R Tyr^723^ (1:2000, Cell Signalling Technologies #3155).

Secondary antibodies: IRDye 680RD Goat anti-Mouse IgG (LI-COR #926-68070) IRDye 800CW Goat anti-Mouse IgG (LI-COR #926-33210), IRDye 680RD Donkey anti Rabbit IgG (LI-COR #926-68073), IRDye 800CW Donkey anti Rabbit IgG (LI-COR #926-32213)

### Flow cytometry

For flow cytometry analysis, 1 million cells were collected by centrifugation at 400 xg, 5 minutes and resuspended in FACS buffer (2% FBS in 1X DPBS). Cells were incubated in fluorescent antibodies for 30 minutes at 4 C in the dark, then washed 3 times in 1 mL of FACS Buffer before fixing with paraformaldehyde (2% final concentration) for 15 minutes at room temperature in the dark. After fixing, cells were washed 3 times in 1 mL FACS buffer before analysis on a Cytek Aurora flow cytometer. Flow cytometry files were analysed using <u>Floreada</u>, an online flow cytometry analysis tool.

Antibodies: FMC63 scFv APC (AcroBiosystems #FM3AY54P1), V5-Tag AlexaFluor 647 (Santa Cruz Biotechnology #sc-271944)

### Luciferase co-culture lysis assay

To quantify live THP1 cells during CAR T cell co-cultures, THP1 cells expressing a luciferase reporter were washed and resuspended in RPMI 1640 supplemented with 10% FBS, then mixed with CAR Jurkat or CAR Primary T cells at E:T ratios ranging from 1:10 to 10:1, with a THP1 only control (0:1). Cells were co-cultured for 24 hours at 37 C, 5% CO2 in a humidified incubator, then collected, washed in 1X PBS, lysed, and measured for luminescence using a Luciferase Assay Kit (Promega #E1500) on a spectrophotometer. Cell lysis (%) is defined as Cell lysis (%) = [*1* − (𝑆𝑎𝑚𝑝𝑙𝑒 𝐿𝑢𝑚. − 𝐵𝑎𝑐𝑘𝑔𝑟𝑜𝑢𝑛𝑑)/𝐵𝑎𝑐𝑘𝑔𝑟𝑜𝑢𝑛𝑑)] × *100* .

Comparisons between mean percent lysis was determined using FWER corrected p-values generated using Fisher’s least significant difference test followed by the Holm-Sidak FWER step-up correction.

### Plotting and Coding

All plotting was performed using Python 3.12.5 with dependencies matplotlib (version 3.9.1), scipy (version 1.14.0), numpy (version 2.0.1), scikit-learn (version 1.5.1), pandas (version 2.2.2), and a custom ‘helpers’ module used for beautifying matplotlib graphs. Coding and plotting were performed in Jupyter interactive python notebooks. Western blot statistical analysis and plotting was performed in Python using a self-coded version of the <u>Holm-Sidak FWER correction algorithm</u>.

## Supplementary Materials

All ImageStudio files and statistical analysis is available in the CAR-Target-Signalling-code- master.zip file. PeptideDepot output, including peak area quantification, peptide sequences, and Ascore is available in the Supplementary Tables.xls file. All code and original plotting outputs are available in the CAR-Target-Signalling-code-master.zip and on GitHub (https://github.com/Aurdeegz/CAR-Target-Signalling-code). Figs. S1 to S9 is in supporting_information.docx.

## Supporting information

Code and Western RAW data

Proteomic Data

## Acknowledgements

The authors would like to thank the Brown University Proteomics Core Facility for managing and maintaining the mass spectrometer used for this work, and for allowing the Salomon laboratory to operate the instrument. We would like to thank the Brown University Genomics Facility for use of the LI-COR Odyssey CLx and Odyssey M scanners, as well as the Flow Cytometry and Cell Sorter Facility for acquiring our flow cytometry samples. This work was supported by National Institutes of Health grant P01AI091580 (A.R.S.), American Cancer Society Research Scholar Grant 135926 (XS), NIGMS MIRA program R35 GM138299 (X.S.), Gabrielle’s Angel Foundation Medical Research Award (X.S.), NCI Exploratory/Developmental Research Grant R21 CA286364 (X.S.), NIH Director’s Transformative Research Award EB037112 (X.S.). The Author contributions include Conceptualisation: A.C., Methodology: A.C., X.Z., Investigation: A.C., A.W., X.Z., A.M., L.Z., Visualisation: A.C., Funding acquisition: X.S., A.R.S., Project administration: A.C., X.S., A.R.S., Supervision: A.C., X.S., A.R.S., Writing – original draft: A.C., A.R.S., Writing – review & editing: A.C., X.S., A.R.S. The authors declare no competing interests. RAW files of mass spectrometry data are available from the ProteomeXchange Consortium via the PRIDE partner repository (Dataset ID: PXD058576, Reviewer Username: Reviewer Password: jvHCJEoOlSme).

