## Supplementary material for "CSF1R-CAR T cells induce CSF1R signalling and promote cancer cell growth": Code and Western RAW data: luciferase_quant.ipynb

```
# %% [code]
from helpers import mpl_plotting_helpers as mph
from helpers import stats_helpers as sh
from helpers import general_helpers as gh
from helpers.mph_modules.dotplots import get_data_info, add_errorbar
import matplotlib.pyplot as plt
import matplotlib.font_manager as mpl_fm
from math import floor, ceil, log2

# %% [markdown]
# Luciferase Assay Data

The cell below contains the raw data for the Luciferase assay, along with labels
used for grouping/statistics/graphing

# %% [code]
car_p = [ [       ["CAR-P CSF1R 0 : 1", [-1.111991, 0.784206, 0.327785]],  # 0
                  ["CAR-P CSF1R 1:10", [-31.876177, -22.488733, -28.262311]], # 1:10
                  ["CAR-P CSF1R 1 : 3", [-36.943769, -27.651735, -23.932052]], # 1:3
                  ["CAR-P CSF1R 1 : 1", [-17.346278, -17.358159, -15.152269]], # 1:1
                  ["CAR-P CSF1R 3 : 1", [18.9761215, 12.9495665, -0.7912461]], # 3:1
                  ["CAR-P CSF1R 10:1", [56.1437999, 65.6489633, 62.9174534]] ],  # 10:1 | THP1-CSF1R
               [  ["CAR-P WT 0 : 1", [-0.285413, -4.266056, 4.55147]],  # 0
                  ["CAR-P WT 1:10", [-12.261224, -1.3177274, 9.22388869]], # 1:10
                  ["CAR-P WT 1 : 3", [-7.8496116, -12.474932, 7.12350855]], # 1:3
                  ["CAR-P WT 1 : 1", [17.3267401, 13.0628848, -7.6915821]], # 1:1
                  ["CAR-P WT 3 : 1", [16.6538661, 7.53634909, -3.7061872]], # 3:1
                  ["CAR-P WT 10:1", [43.6574817, 30.9364689, 41.4509199]] ]] # THP1-WT

car_j = [ [       ["CAR-J CSF1R 0 : 1",[-1.69746, 4.928515, -3.231054]],  # 0
                  ["CAR-J CSF1R 1:10",[-9.1390943, 8.96966103, 4.42772432]], # 1:10
                  ["CAR-J CSF1R 1 : 3",[-20.937491, 11.4893335, -2.1847127]], # 1:3
                  ["CAR-J CSF1R 1 : 1",[-11.015422, -36.288622, -20.576581]], # 1:1
                  ["CAR-J CSF1R 3 : 1",[-47.561617, -25.109084, -29.338483 ]] ],  # 3:1 | THP1-CSF1R
               [  ["CAR-J WT 0 : 1", [-1.9101139, 0.99490184, 0.91521198]],  # 0
                  ["CAR-J WT 1:10", [-1.5151717, 1.38049909, -19.861034]], # 1:10
                  ["CAR-J WT 1 : 3", [-6.0054965, 2.10943127, 11.8901229]], # 1:3
                  ["CAR-J WT 1 : 1", [6.17780336, -0.790547, 5.22372175]], # 1:1
                  ["CAR-J WT 3 : 1", [-11.803599, -9.2052019, -1.1808156]] ]] # 3:1 |  THP1-WT

xticks = ["0:1", "1:10", "1:3", "1:1", "3:1", "10:1"]

# %% [markdown]
# Line Plot graphs

Below is some code to (a) determine which combinations to ignore, and (b)
create line plots. My HolmSidak class automatically does an ANOVA, determines
whether or not multiple compaisons are warranted, performs Fisher's LSD, then
uses the Holm-Sidak step up correction to correct for FWER. The comparisons to be
ignored are ones between different groups and different E:T ratios 
(for example, CAR-P WT 1:1 CAR-P CSF1R 1:10).

# %% [code]
def _logical_ignore_comps(labelled_line_groups,
                          group_strs,
                          xgroup_strs):
    """
    Only want to compare along a line group (e.g. timecourse) or
    down an x-column (e.g. JE6 DMSO 0m vs JE6 U0126 0m), but not
    all the random other comparisons because statistically they're
    kind of useless
    
    So this function will find all of the pairs that are useless
    """
    groups_unpacked = []
    for group in labelled_line_groups:
        groups_unpacked += group
    # This will hold the ignored pairs
    ignore_me_senpai = []
    # First, get all pairs
    paired = gh.make_pairs(groups_unpacked,
                           dupes = False,
                           reverse = False)
    # Then iterate over and check the labels
    for p in paired:
        gs_check = 0
        xs_check = 0
        # Check all the group strings
        for gs in group_strs:
            if gs_check == 1:
                pass
            elif gs in p[0][0] and gs in p[1][0]:
                gs_check = 1
        # Check all the xgroup strings
        for xs in xgroup_strs:
            if xs_check == 1:
                pass
            elif xs in p[0][0] and xs in p[1][0]:
                xs_check = 1
        # If there isn't a match, in either, ignore
        if gs_check == 0 and xs_check == 0:
            ignore_me_senpai.append(p)
    # Return the ignored pairs at the end
    return ignore_me_senpai

def perform_line_statistics(labelled_line_groups,
                            ignore_comps,
                            comp_type,
                            statsfile):
    """
    labelled_line_groups -> data with labels
                            list of lists of [label, [d1,d2,...,dn]]
    ignore_comps -> list of pairs ("group 1", "group 2") to not be
                    compared
    comp_type -> statistics to use, currently only
                 ["HolmSidak", "TukeyHSD"] are supported
                 (both do an ANOVA first by default)
    statsfile -> a string to the output path and filename
                 for the statistics file output
    #####
    Returns None, just dumps the statsfile
    """
    assert comp_type in ["HolmSidak", "TukeyHSD"], f"Invalid comparison type: {comp_type}"
    groups_unpacked = []
    for group in labelled_line_groups:
        groups_unpacked += group
    if comp_type == "HolmSidak":
        comparison = sh.HolmSidak(*groups_unpacked,
                                  labels = True,
                                  override = True,
                                  alpha = 0.05,
                                  no_comp = ignore_comps)
    elif comp_type == "TukeyHSD":
        comparison = sh.TukeyHSD(*groups_unpacked,
                                  labels = True,
                                  override = True,
                                  alpha = 0.05,
                                  no_comp = ignore_comps)
    comparison.write_output(filename = statsfile,
                            file_type = "csv")
    return None

def find_centres(plotting_info):
    """
    plotting_info -> output from get_data_info, a list of
                     data info and the raw data
                     
    goal: grab the centres for xticks
    """
    centres = []
    for group in plotting_info:
        if len(centres) <= len(group[0]["centers"]):
            centres = group[0]["centers"]
    return centres

def line_plot(labelled_line_groups,
              show_points = False,
              show_legend = False,
              colours = ["grey" for _ in range(20)],
              group_labs = [f"Thing {i}" for i in range(20)],
              markers = ["s" for _ in range(20)],
              linestyles = ["solid" for _ in range(20)],
              xlabels = [f"Time {i}" for i in range(20)],
              ylabel = ["Fold change"],
              ylims = None,
              ignore_comps = [],
              statsfile = None,
              comp_type = "HolmSidak",
              figfile = None):
    """
    labelled_line_groups -> list of lists, where each sublist contains labelled groups
    """
    # First, get some basic plotting information
    plotting_info = [get_data_info(line) for line in labelled_line_groups]
    # Then manage the statistics
    if statsfile != None:
        perform_line_statistics(labelled_line_groups, 
                                ignore_comps, 
                                comp_type, 
                                statsfile)
    # Begin plotting c::
    if ylims == None:
        ylims = floor(min([item for item in gh.unpack_list(labelled_line_groups) if type(item) in [int, float]])), ceil(max([item for item in gh.unpack_list(labelled_line_groups) if type(item) in [int, float]]))
    # 
    fig, ax = plt.subplots(figsize = (6,6))
    # 
    for i in range(len(labelled_line_groups)):
        #
        ax.plot(plotting_info[i][0]["centers"],
                plotting_info[i][0]["means"],
                color = colours[i],
                label = group_labs[i],
                linestyle = linestyles[i])
        #
        for j in range(len(labelled_line_groups[i])):
            add_errorbar(ax, 
                         plotting_info[i][0]["centers"][j],
                         plotting_info[i][0]["means"][j],
                         plotting_info[i][0]["sems"][j],
                         color = colours[i])
            if show_points:
            #
                ax.scatter(plotting_info[i][0]["xs"][j],
                           plotting_info[i][1][j][1],
                           color = colours[i],
                           edgecolor = "black", alpha = 0.3,
                           marker = markers[i],
                           s = 10)
            else:
            #
                ax.scatter(plotting_info[i][0]["centers"],
                           plotting_info[i][0]["means"],
                           color = colours[i],
                           edgecolor = "black", alpha = 0.3,
                           marker = markers[i],
                           s = 30)
    ax.spines["top"].set_visible(False)
    ax.spines["right"].set_visible(False)
    xticks = find_centres(plotting_info)
    ax.set_xticks(xticks)
    ax.set_xticklabels(xlabels[:len(xticks)],
                       fontfamily = "sans-serif", 
                       font = "Arial", 
                       fontweight = "bold", 
                       fontsize = 12,
                       rotation = 90,
                       ha = "center")
    ax.set_ylim(*ylims)
    mph.update_ticks(ax, which = "y")
    if show_legend:
        ax.legend(loc = "best",
                  prop = mpl_fm.FontProperties(family = "sans-serif",
                                               weight = "bold"))
    if figfile == None:
        plt.show()
    else:
        plt.savefig(figfile)
    plt.close()
    return None

# %% [code]
j_ignore = _logical_ignore_comps(car_j, 
                                 group_strs = ["CAR-J CSF1R",
                                               "CAR-J WT"],
                                 xgroup_strs = ["0 : 1",
                                                "1:10",
                                                "1 : 3",
                                                "1 : 1",
                                                "3 : 1",
                                                "10:1"])


line_plot(car_j,
          ylims = [-80,80],
          colours = ["deeppink", "darkviolet", "hotpink", "cyan"],
          markers = ["s", "s", "o", "o"],
          linestyles = ["solid", "solid", "dashdot", "dashdot"],
          xlabels = xticks[:-1],
          show_points = True,
          show_legend = False,
          group_labs = ["THP-1 CSF1R", "THP-1 WT"],
          ignore_comps = j_ignore,
          statsfile = "stats_out/carj_holmsidak_stats",
          figfile = "figs/carj_thp1_growth.pdf"          )


# %% [code]
p_ignore = _logical_ignore_comps(car_p, 
                                 group_strs = ["CAR-P CSF1R",
                                               "CAR-P WT"],
                                 xgroup_strs = ["0 : 1",
                                                "1:10",
                                                "1 : 3",
                                                "1 : 1",
                                                "3 : 1",
                                                "10:1"])

line_plot(car_p,
          ylims = [-80,80],
          colours = ["deeppink", "darkviolet", "hotpink", "cyan"],
          markers = ["s", "s", "o", "o"],
          linestyles = ["solid", "solid", "dashdot", "dashdot"],
          xlabels = xticks,
          #show_points = True,
          show_legend = False,
          group_labs = ["THP-1 CSF1R", "THP-1 WT"],
          ignore_comps = p_ignore,
          statsfile = "stats_out/carp_holmsidak_stats",
          figfile = "figs/carp_thp1_growth.pdf",
          )
```
