## Supplementary material for "CSF1R-CAR T cells induce CSF1R signalling and promote cancer cell growth": Code and Western RAW data: run_ptmsea.R

```
source("/mnt/c/Users/redas/Desktop/jupyter_directory/salomon_lab_folder/car_target_project/r_scripts/ssGSEA2.0.R")
gct_file <- "/mnt/c/Users/redas/Desktop/jupyter_directory/salomon_lab_folder/car_target_project/28bbz_raji/pY_data/qvalue.gct"
output_loc <- "/mnt/c/Users/redas/Desktop/jupyter_directory/salomon_lab_folder/car_target_project/28bbz_raji/pY_data/output"
sig_db <- "/mnt/c/Users/redas/Desktop/jupyter_directory/salomon_lab_folder/car_target_project/database/ptm.sig.db.all.flanking.human.v2.0.0.gmt"
signat.all <- unlist(lapply(sig_db, readLines))
signat.all <- strsplit(signat.all, '	')
names(signat.all) <- sapply(signat.all, function(x)x[1])
signat.all <- lapply(signat.all, function(x) x[-c(1,2)])
names(gct_file) <- paste(  sub('\\.gct$', '', sub('.*/','', gct_file)), 'ssGSEA', sep='_' )
input.ds <- gct_file
i <- 1
gsea.res <-
  ssGSEA2(
    "/mnt/c/Users/redas/Desktop/jupyter_directory/salomon_lab_folder/car_target_project/28bbz_raji/pY_data/qvalue.gct",
    gene.set.databases=sig_db,
    sample.norm.type="none",
    weight=1,
    statistic="area.under.RES",
    output.score.type="NES",
    nperm=10000,
    min.overlap=3,
    correl.type="z.score",
    output.prefix=output_loc,
    par=T,
    spare.cores=1,
    export.signat.gct=F,
    extended.output=T,
    param.file=T,
    global.fdr=TRUE,
  )
```
