## Supplementary figures and images for "CSF1R-CAR T cells induce CSF1R signalling and promote cancer cell growth"

### cfl1_$^{y85}$_foldchange_m.pdf

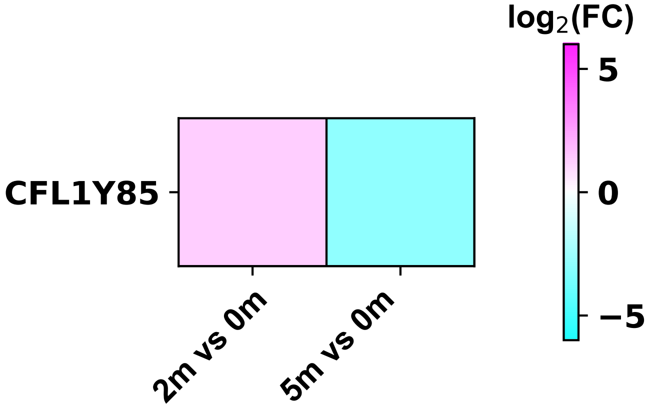

### cgn_$^{y105}$_foldchange_m.pdf

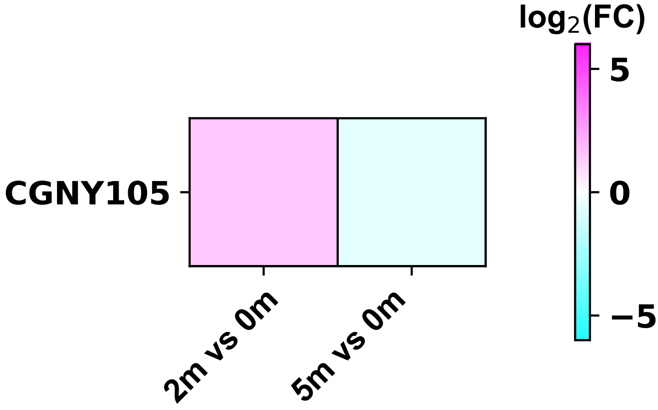

### cherp_$^{y894}$_foldchange_m.pdf

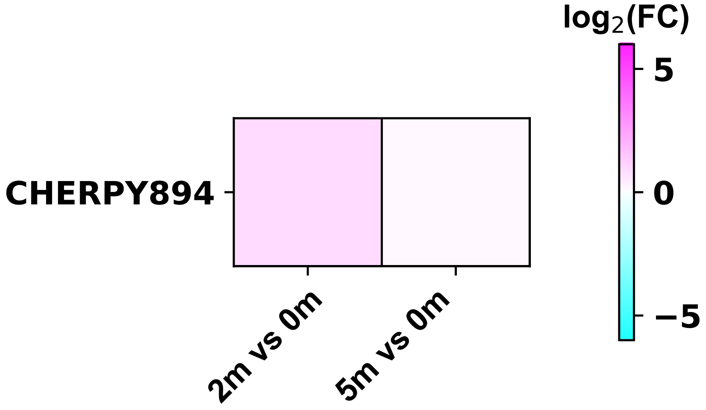

### chn1_$^{y143}$_foldchange_m.pdf

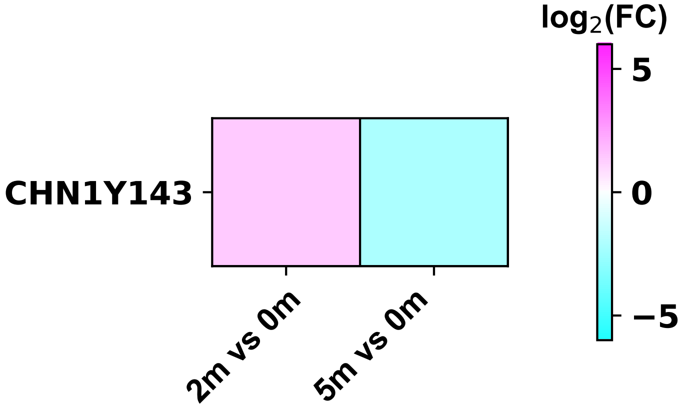

### cirbp_$^{y164}$_foldchange_m.pdf

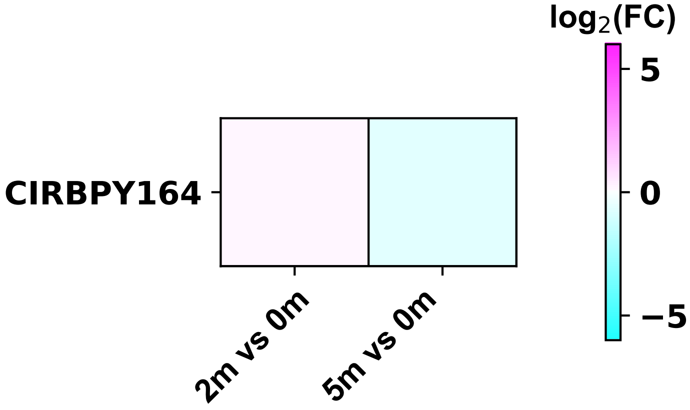

### cirbp_$^{y167}$_foldchange_m.pdf

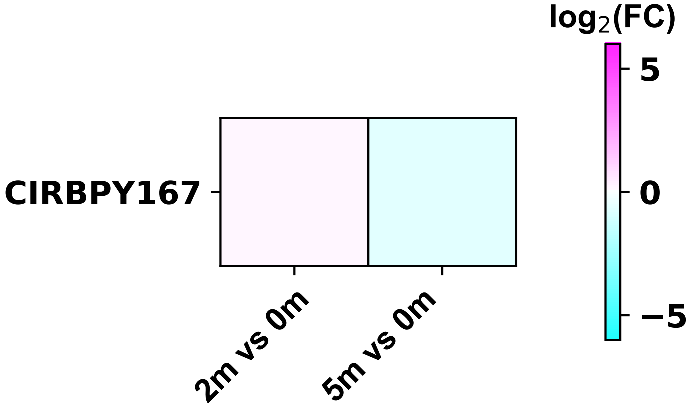

### clasp2_$^{y1234}$_foldchange_m.pdf

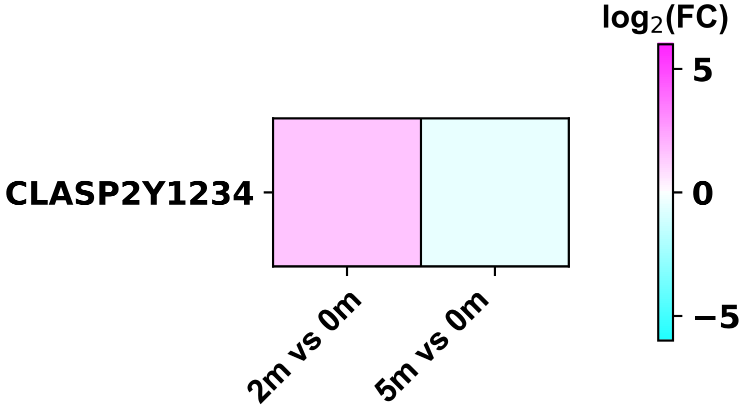

### cltc_$^{y430}$_foldchange_m.pdf

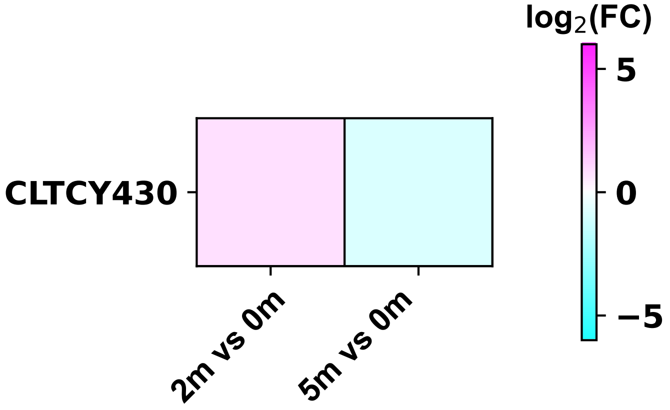

### cltc_$^{y634}$_foldchange_m.pdf

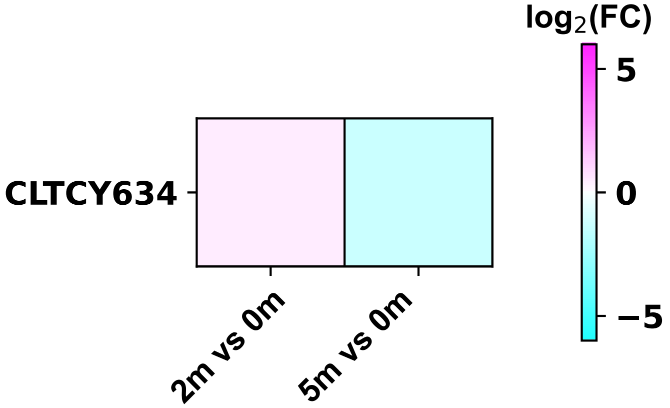

### cltc_$^{y1477}$_foldchange_m.pdf

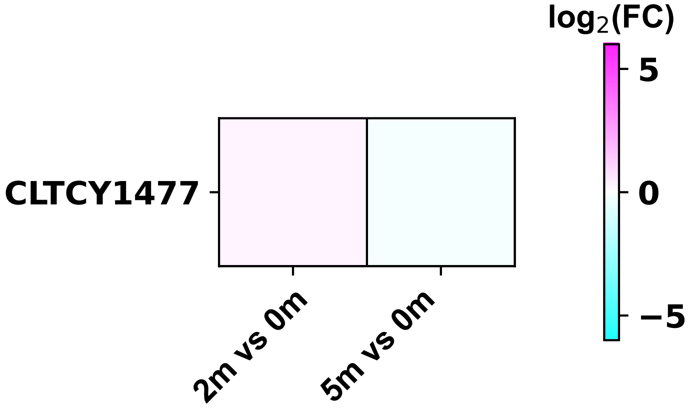

### cltc_$^{y1487}$_foldchange_m.pdf

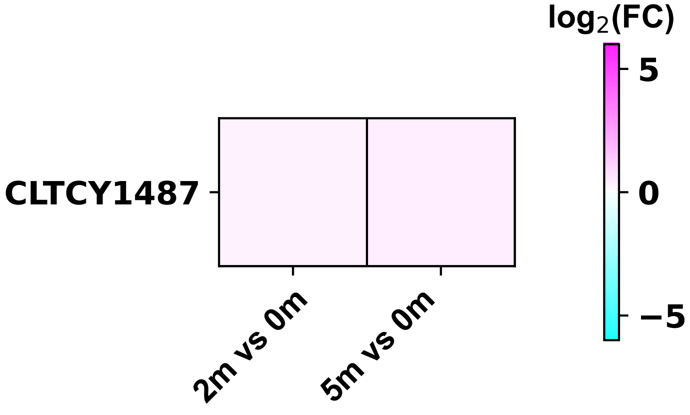

### cnot9_$^{y23}$_foldchange_m.pdf

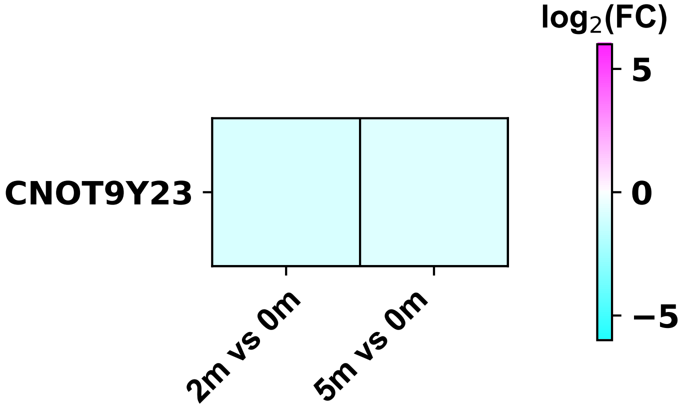

### coro1c_$^{y301}$_foldchange_m.pdf

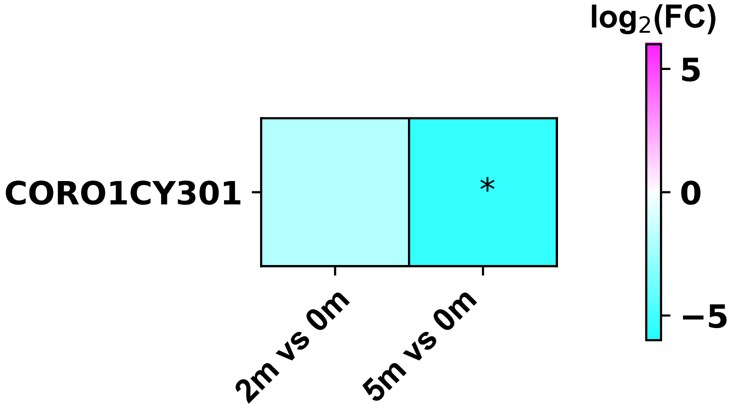

### crk_$^{y190}$_foldchange_m.pdf

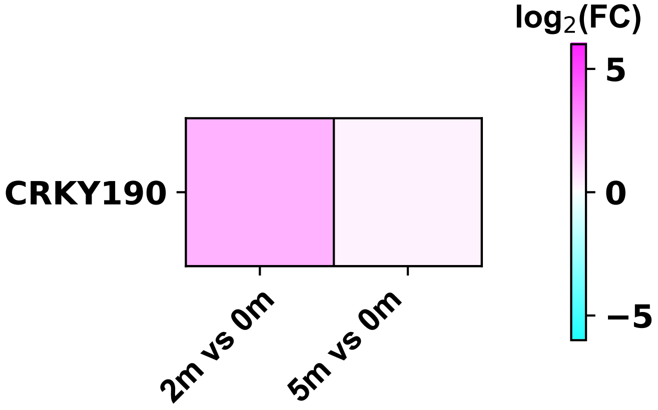

### crkl_$^{y198}$_foldchange_m.pdf

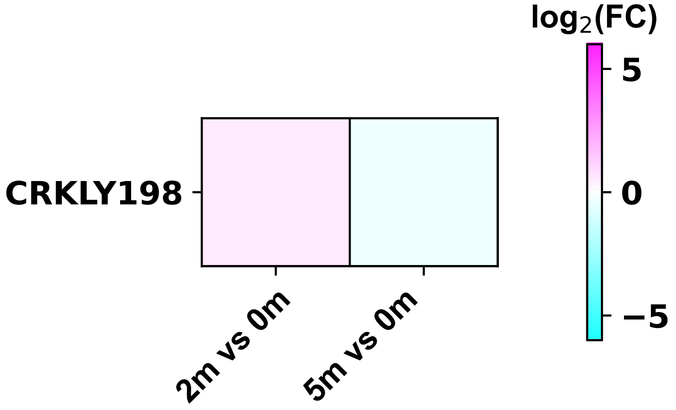

### crkl_$^{y207}$_foldchange_m.pdf

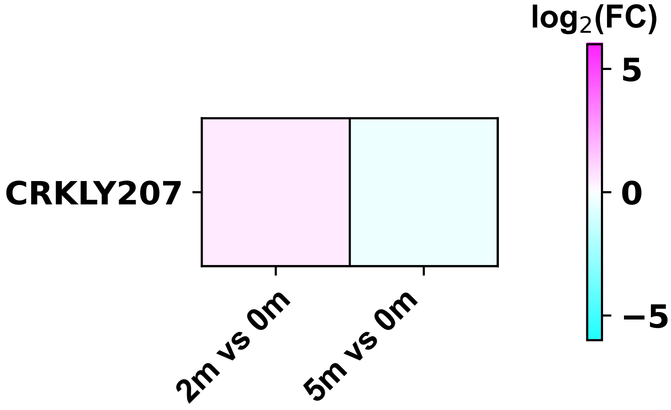

### cttn_$^{y421}$_foldchange_m.pdf

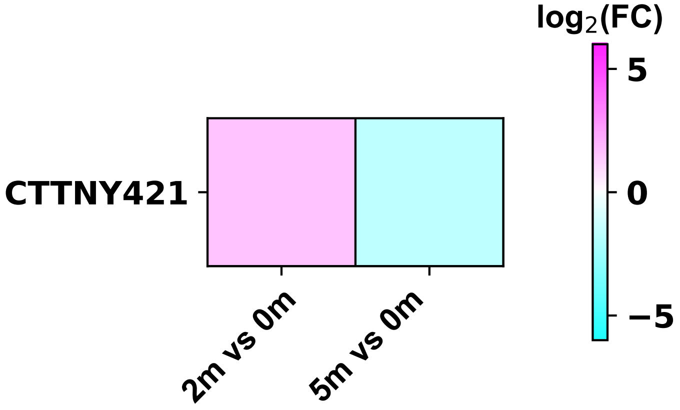

### dbi_$^{y46}$_foldchange_m.pdf

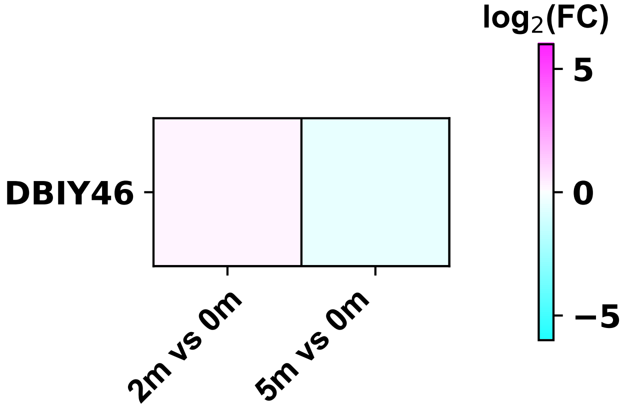

### dbnl_$^{y162}$_foldchange_m.pdf

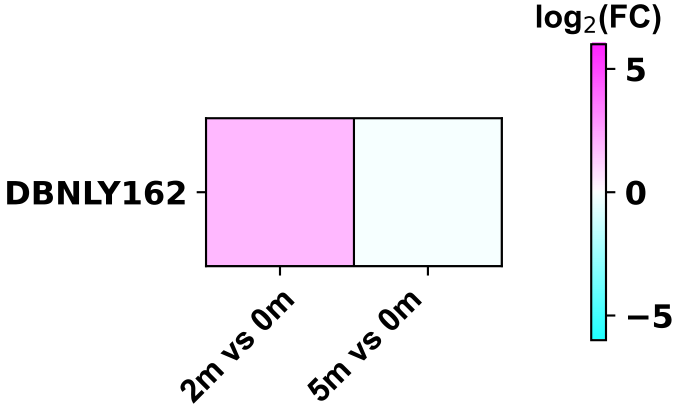

### dcp1b_$^{y133}$_foldchange_m.pdf

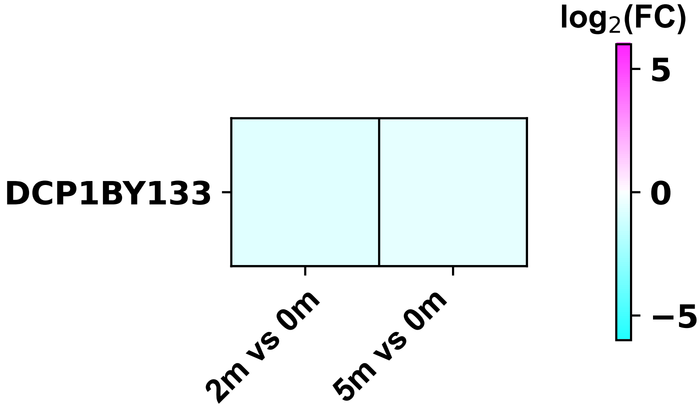

### dcp1b_$^{y191}$_foldchange_m.pdf

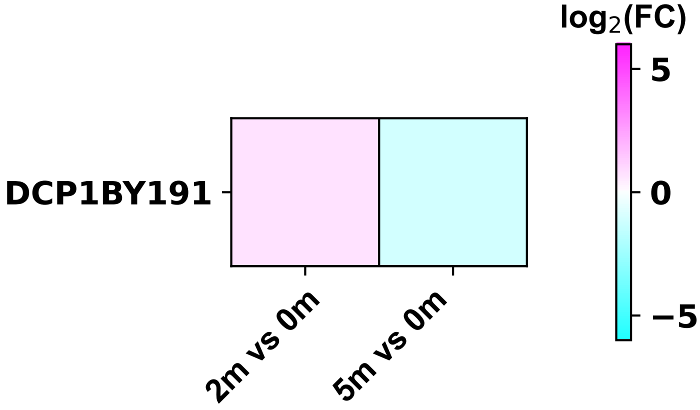

### dctn2_$^{y6}$_foldchange_m.pdf

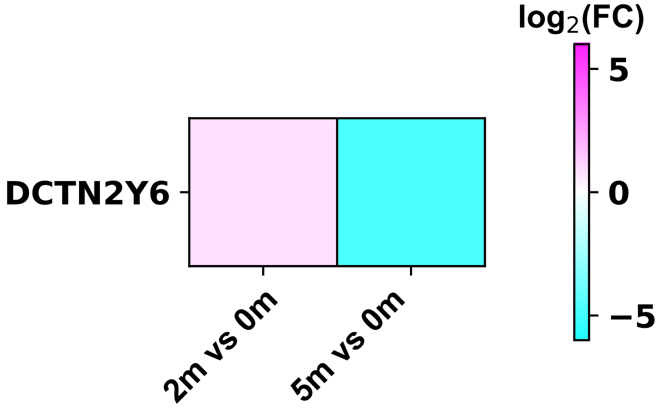

### dctn2_$^{y91}$_foldchange_m.pdf

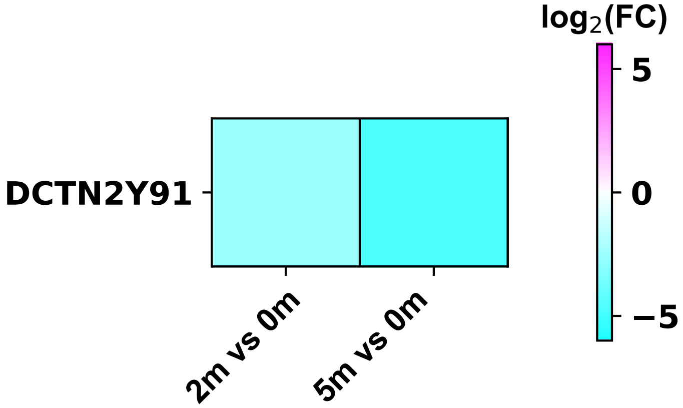

### ddx3x_$^{y53}$_foldchange_m.pdf

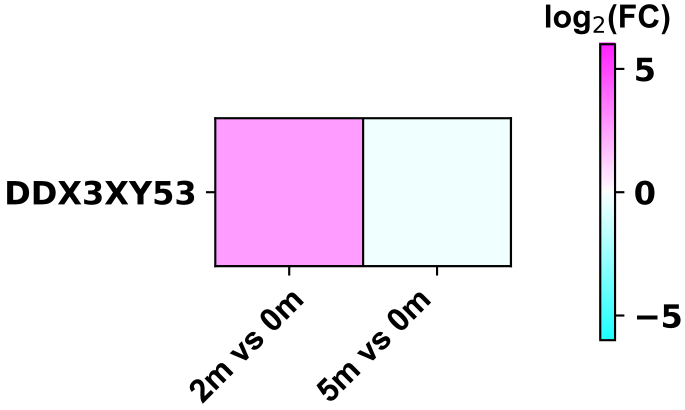

### ddx5_$^{y416}$_foldchange_m.pdf

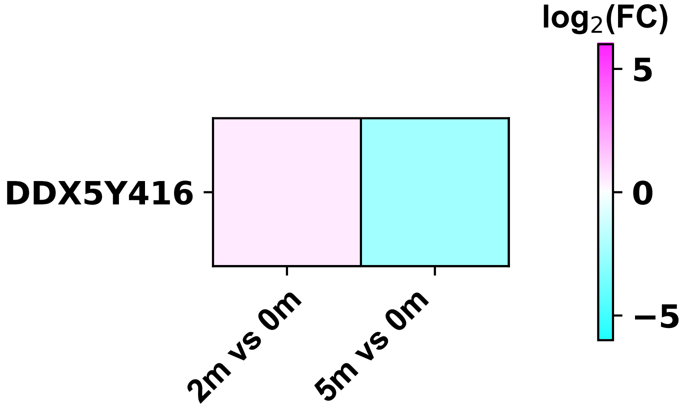

### ddx20_$^{y756}$_foldchange_m.pdf

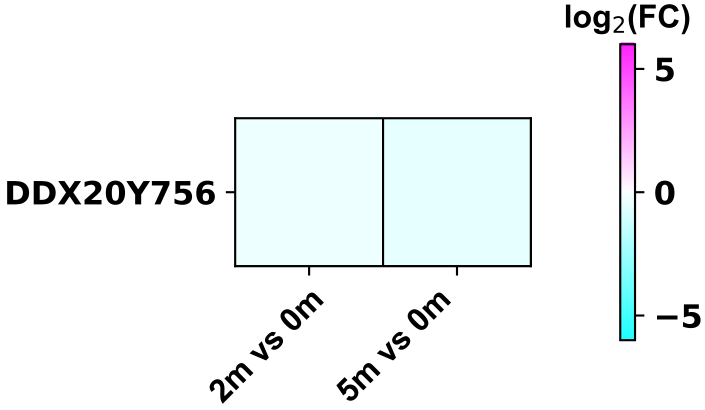

### diaph1_$^{y374}$_foldchange_m.pdf

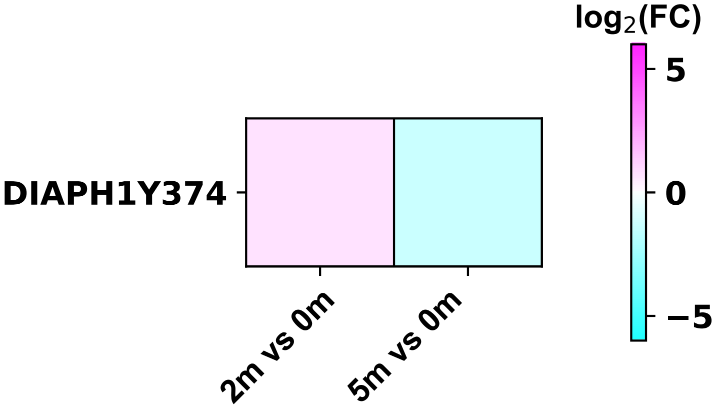

### diaph1_$^{y424}$_foldchange_m.pdf

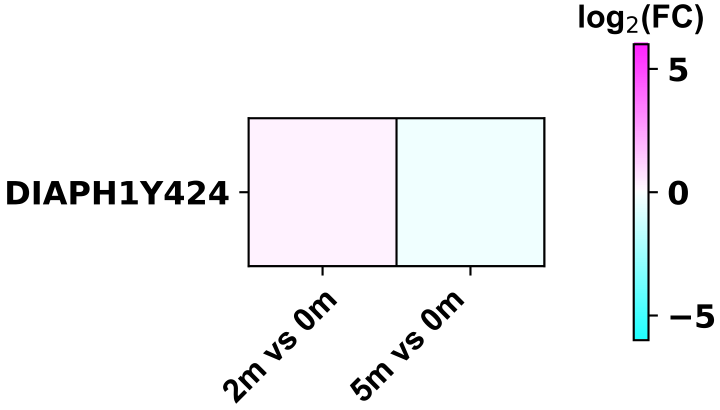

### dido1_$^{y1700}$_foldchange_m.pdf

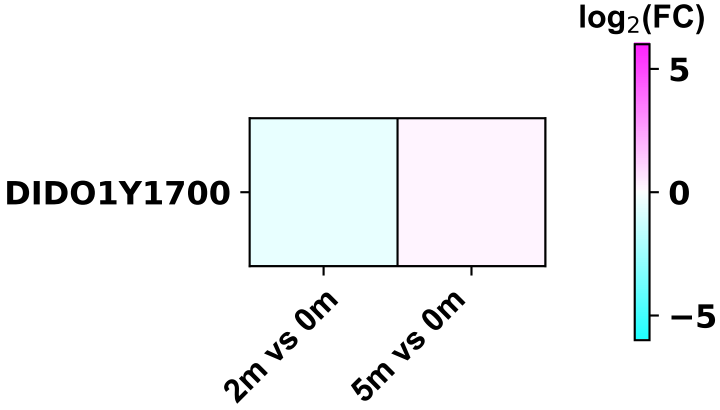

### dnajb1_$^{y176}$_foldchange_m.pdf

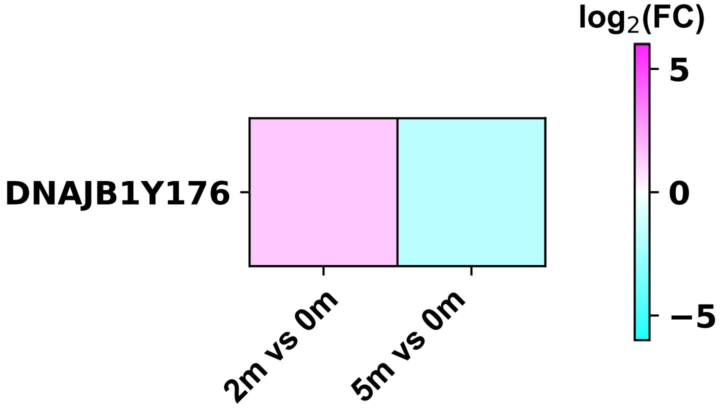

### dnajc2_$^{y90}$_foldchange_m.pdf

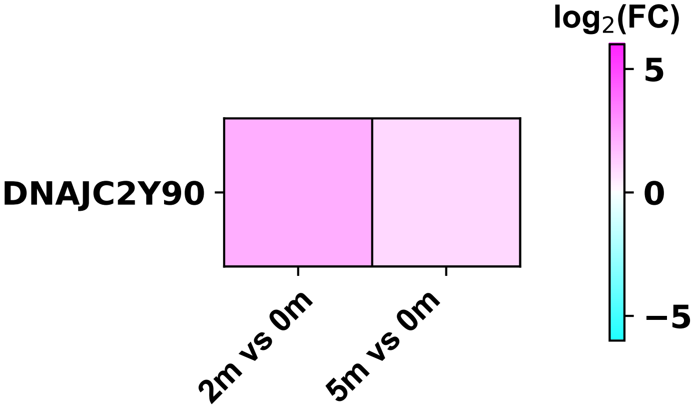

### dock2_$^{y212}$_foldchange_m.pdf

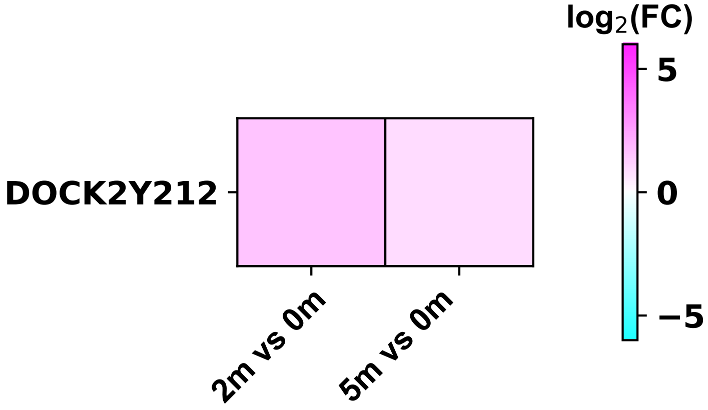

### dock11_$^{y1188}$_foldchange_m.pdf

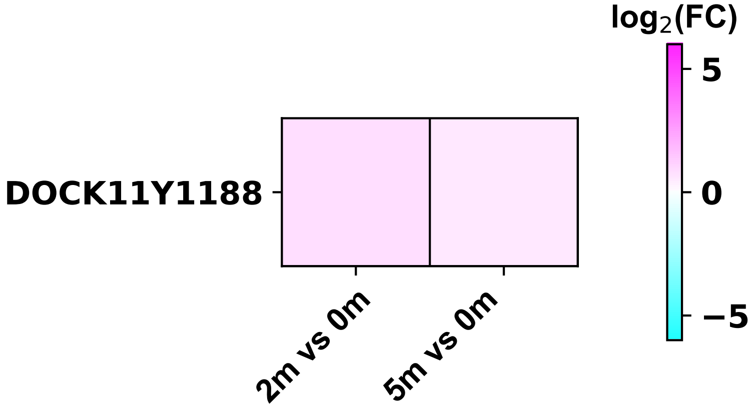

### dok1_$^{y409}$_foldchange_m.pdf

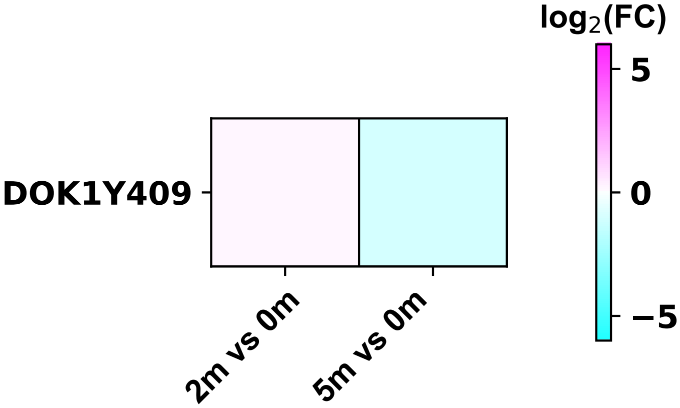

### dok2_$^{y299}$_foldchange_m.pdf

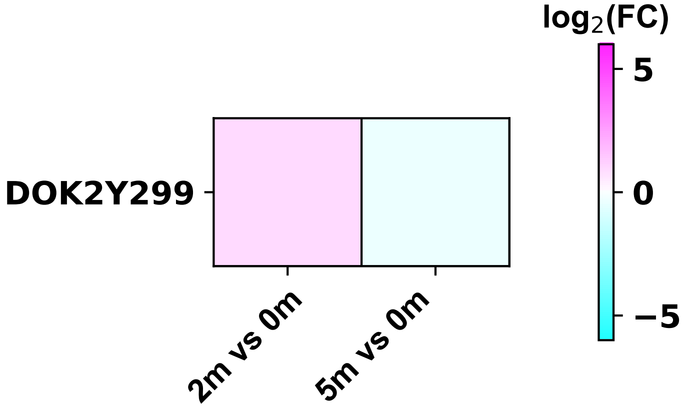

### dpysl2_$^{y499}$_foldchange_m.pdf

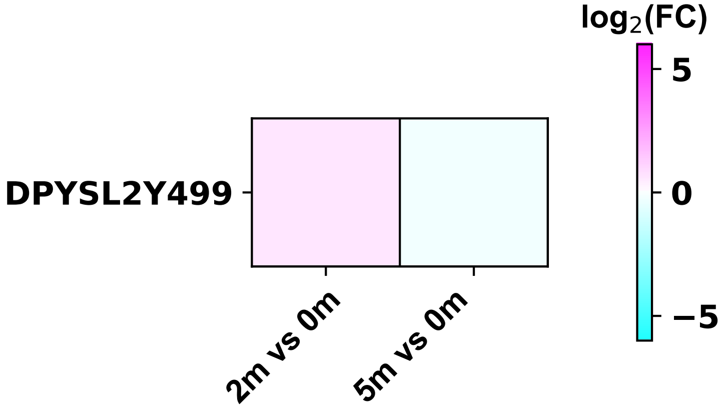

### dync1h1_$^{y1010}$_foldchange_m.pdf

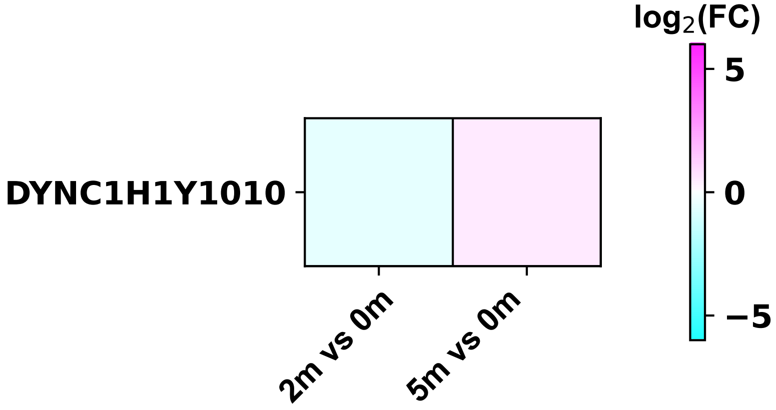

### dyrk1a_$^{y145}$_foldchange_m.pdf

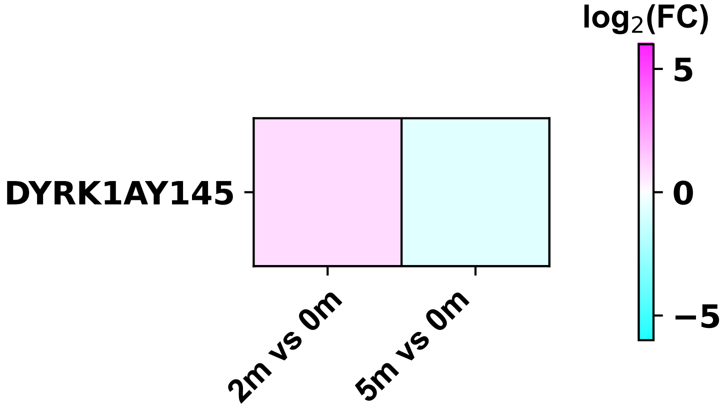

### dyrk1a_$^{y273}$_foldchange_m.pdf

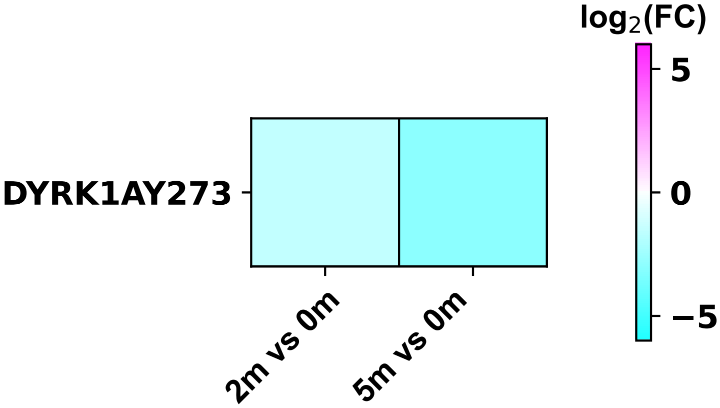

### dyrk3_$^{y369}$_foldchange_m.pdf

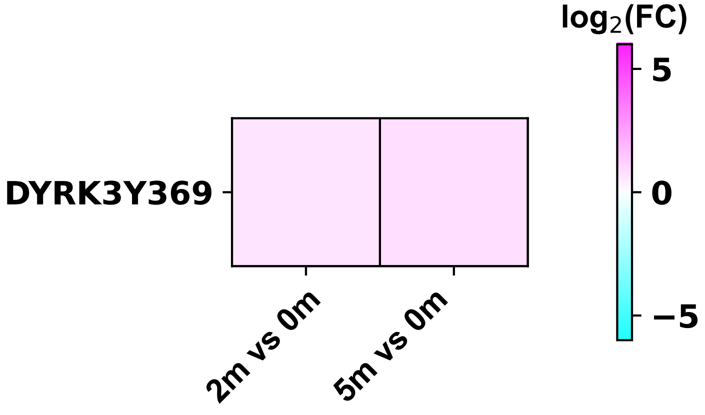

### eef1a1_$^{y85}$_foldchange_m.pdf

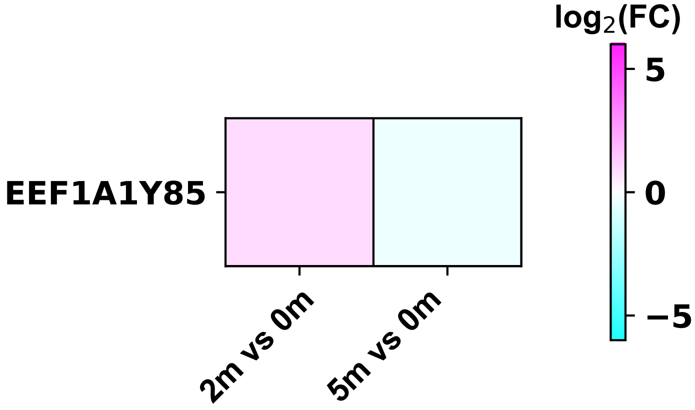

### eef1a1_$^{y86}$_foldchange_m.pdf

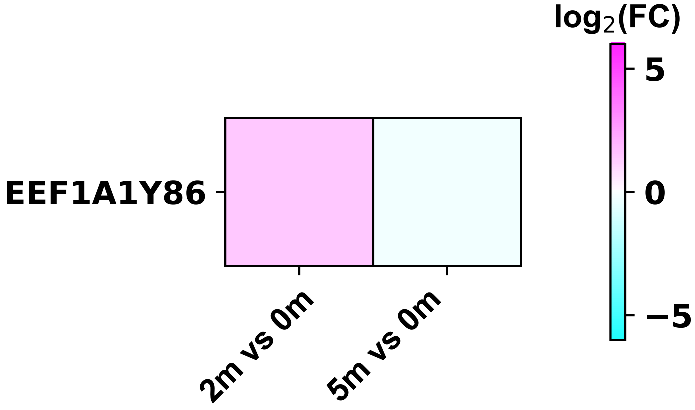

### eef1a1_$^{y141}$_foldchange_m.pdf

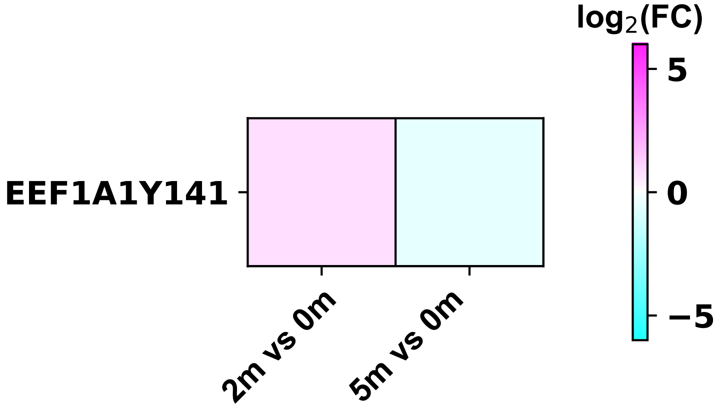

### eef1b2_$^{y79}$_foldchange_m.pdf

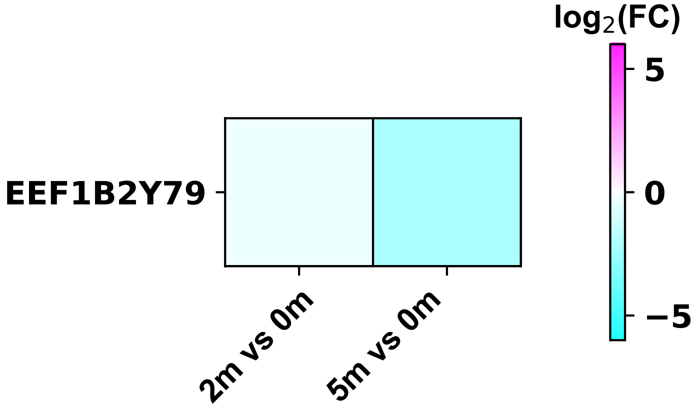

### eef1d_$^{y392}$_foldchange_m.pdf

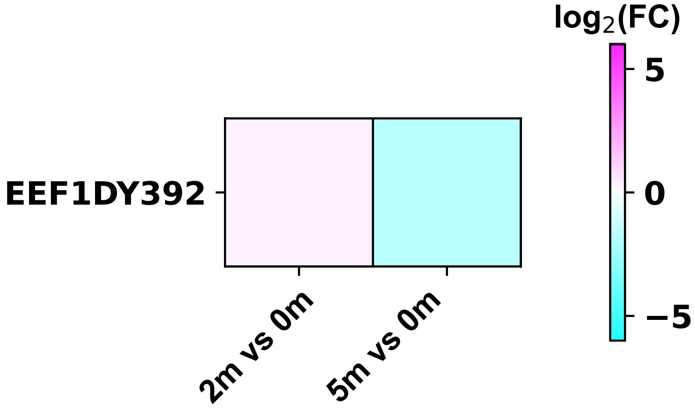

### eif3b_$^{y449}$_foldchange_m.pdf

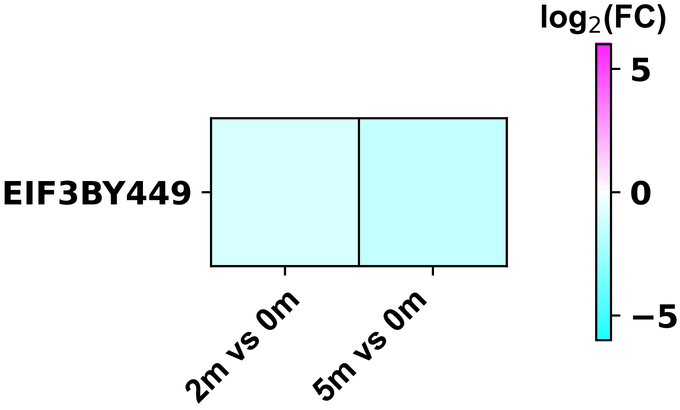

### eif3d_$^{y318}$_foldchange_m.pdf

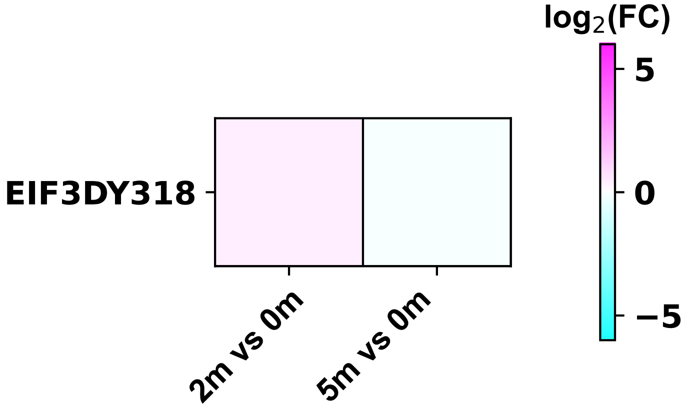

### eif4a1_$^{y71}$_foldchange_m.pdf

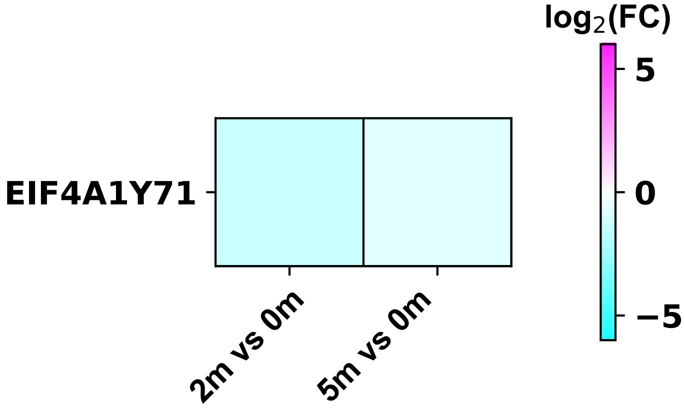

### eif4g3_$^{y123}$_foldchange_m.pdf

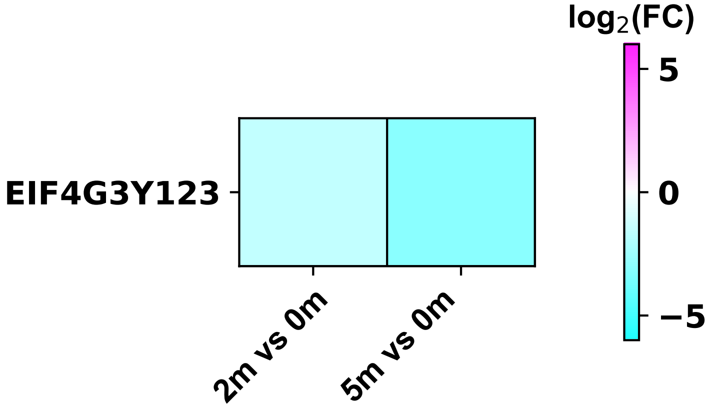

### eif5a2_$^{y69}$_foldchange_m.pdf

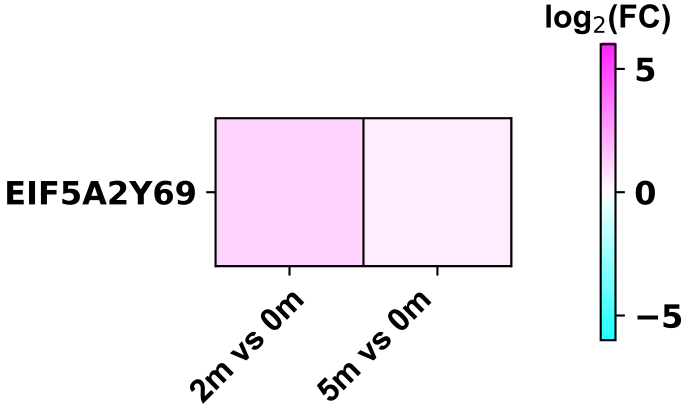

### elmo1_$^{y724}$_foldchange_m.pdf

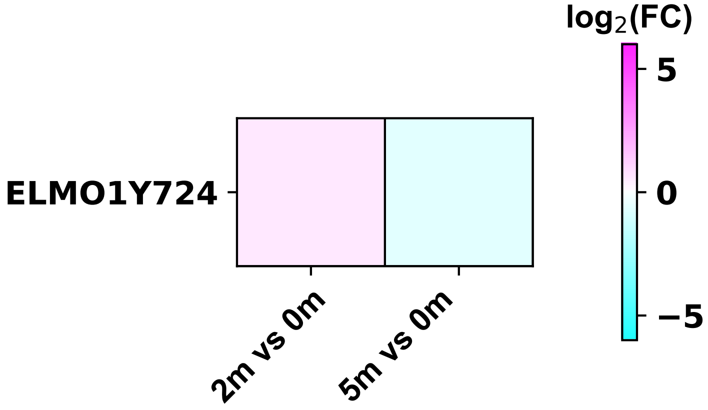

### elp3_$^{y202}$_foldchange_m.pdf

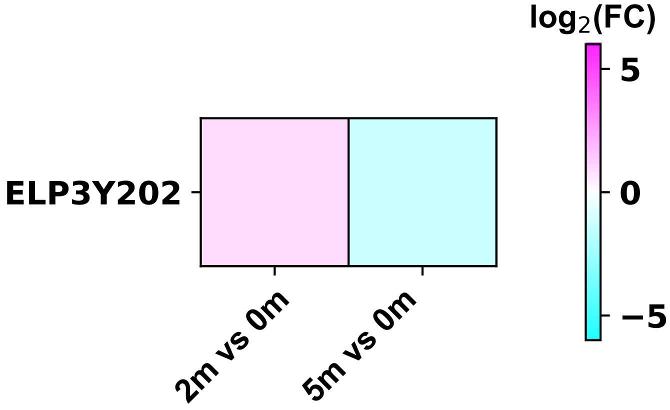

### eno1_$^{y44}$_foldchange_m.pdf

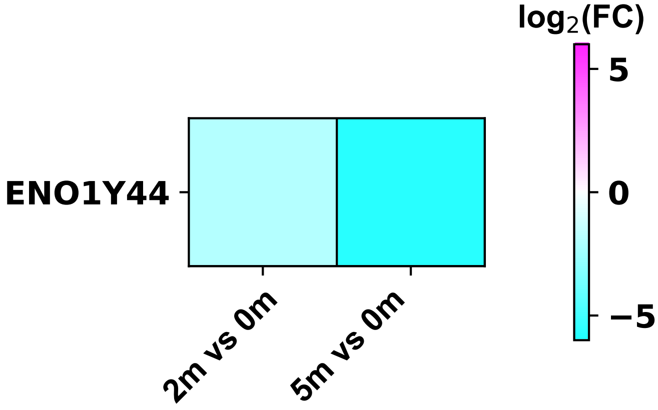

### eno3_$^{y44}$_foldchange_m.pdf

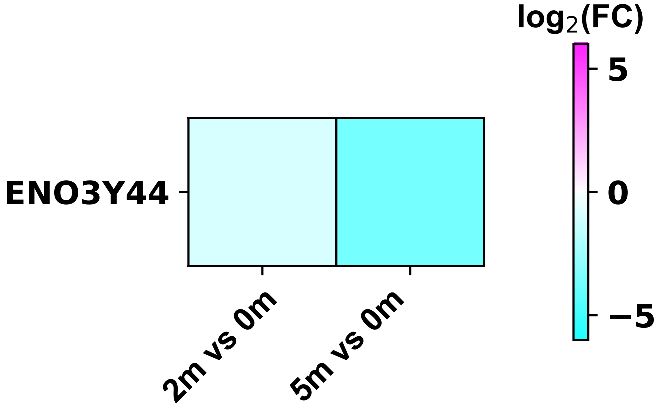

### ensa_$^{y65}$_foldchange_m.pdf

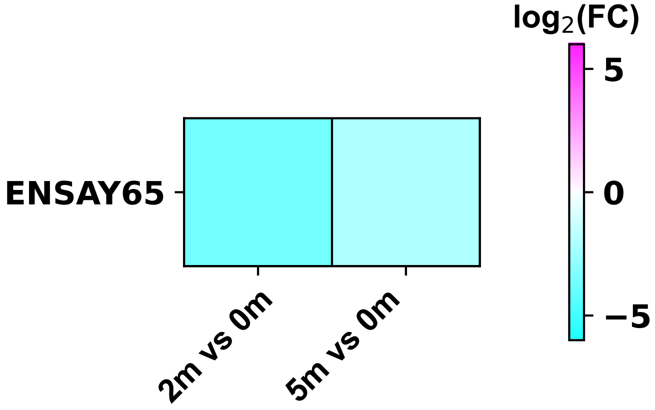

### epb41_$^{y222}$_foldchange_m.pdf

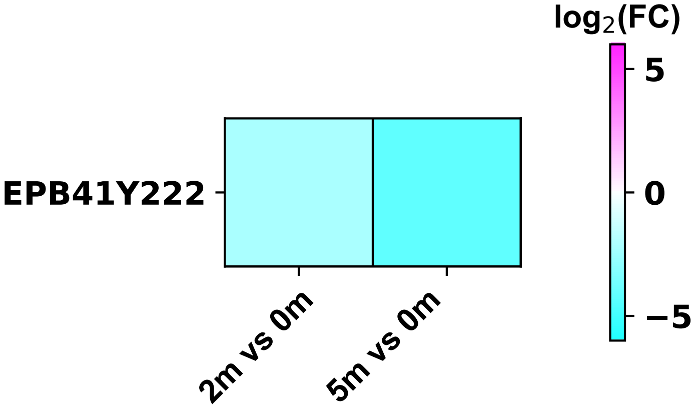

### epb41l2_$^{y773}$_foldchange_m.pdf

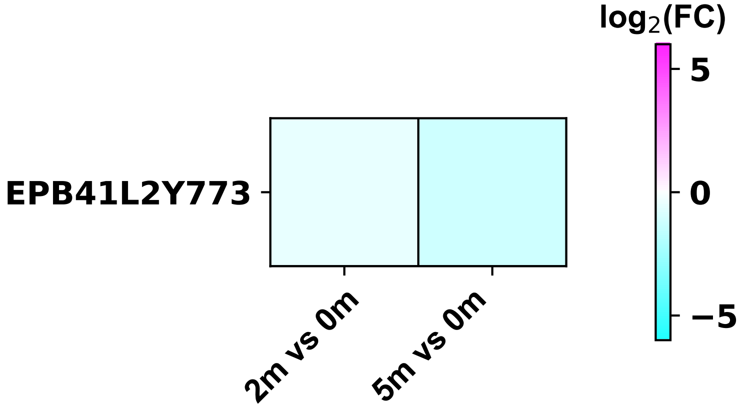

### epha5_$^{y779}$_foldchange_m.pdf

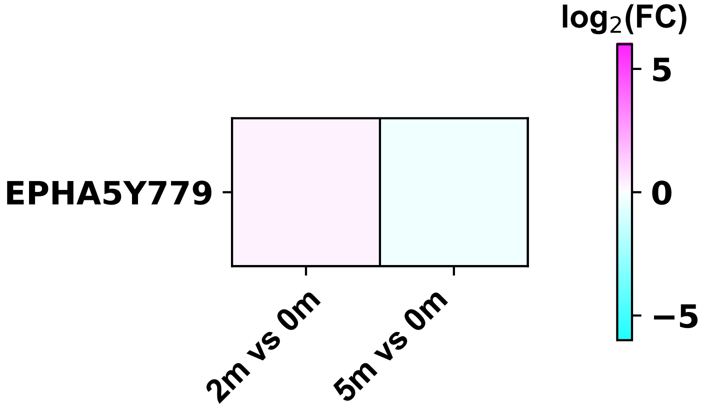

### eprs_$^{y827}$_foldchange_m.pdf

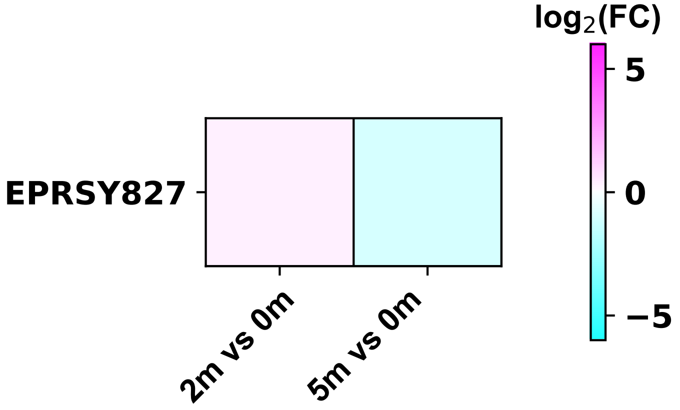

### erbin_$^{y884}$_foldchange_m.pdf

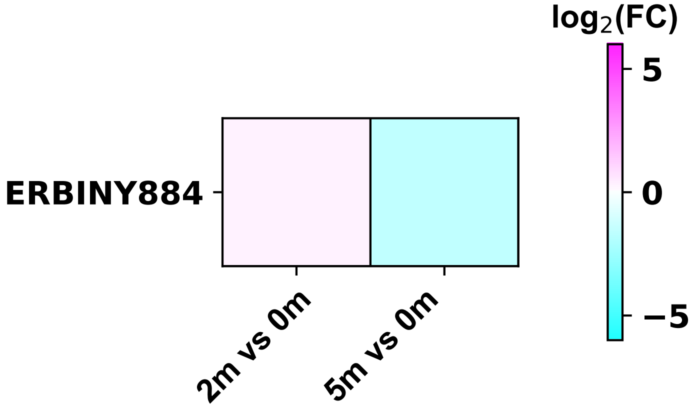

### ercc3_$^{y758}$_foldchange_m.pdf

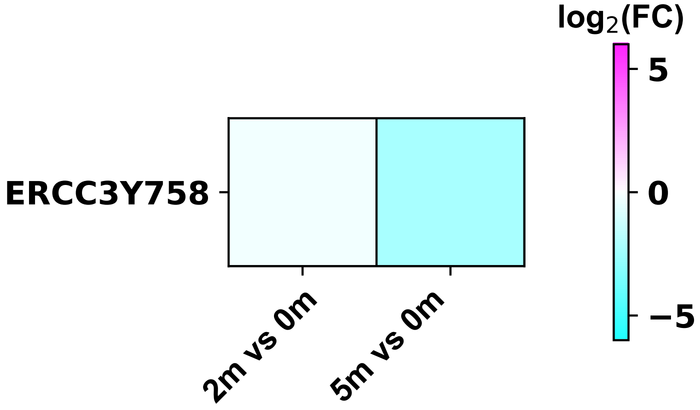

### erh_$^{y92}$_foldchange_m.pdf

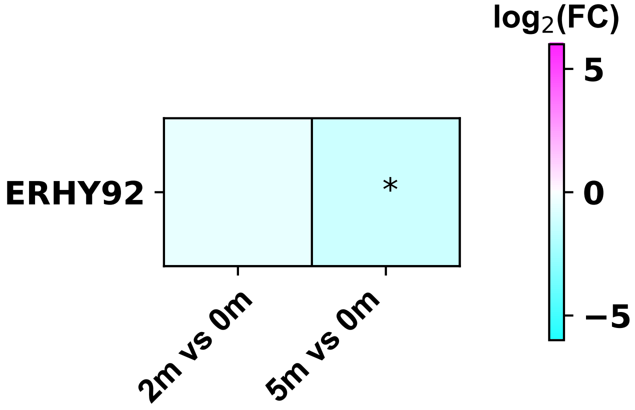

### eri1_$^{y66}$_foldchange_m.pdf

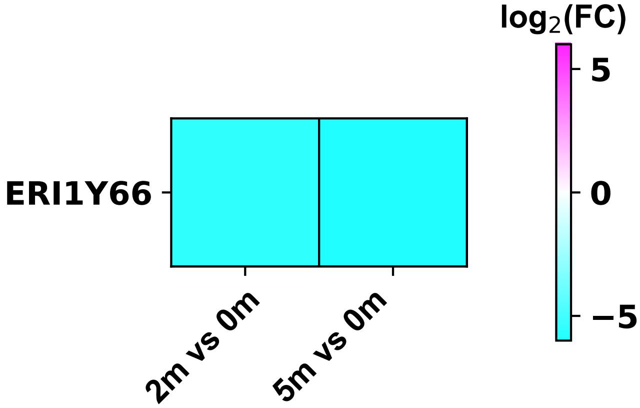

### esyt1_$^{y822}$_foldchange_m.pdf

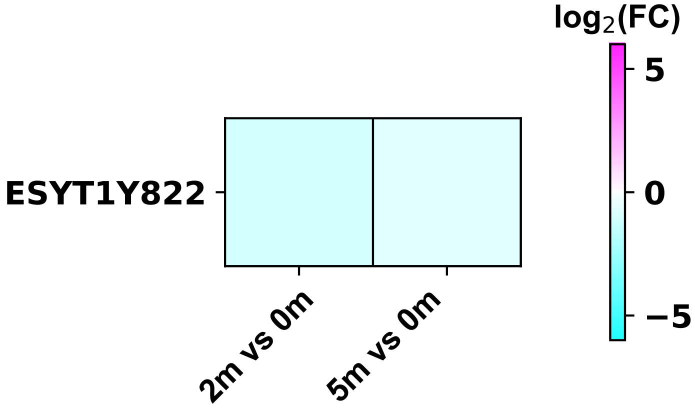

### fam20b_$^{y138}$_foldchange_m.pdf

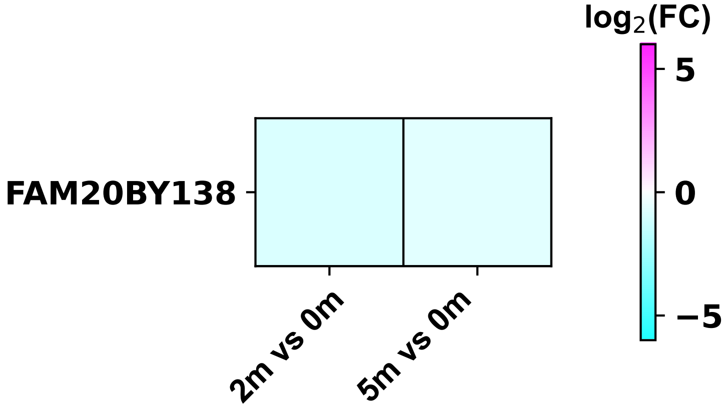

### fam208a_$^{y409}$_foldchange_m.pdf

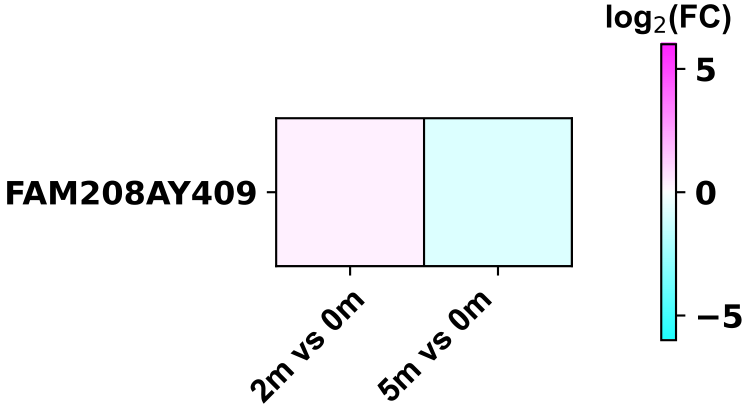

### fermt3_$^{y11}$_foldchange_m.pdf

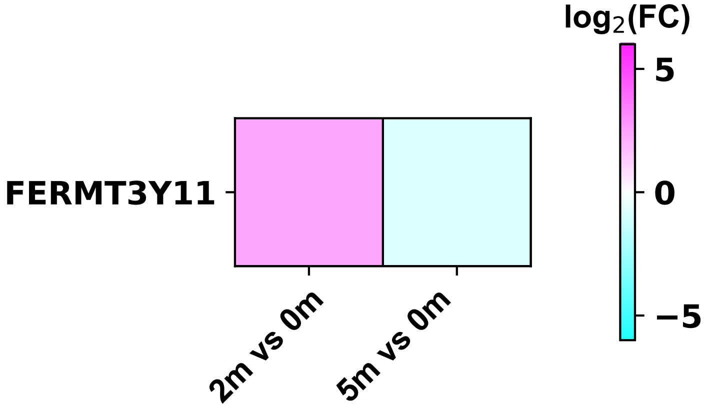

### fermt3_$^{y504}$_foldchange_m.pdf

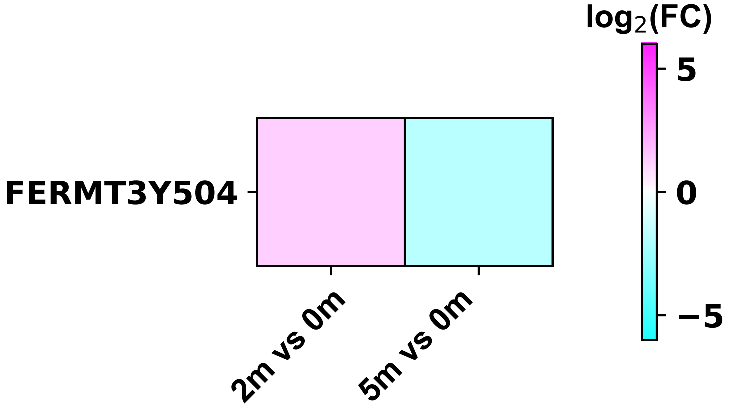

### fgfr1op_$^{y337}$_foldchange_m.pdf

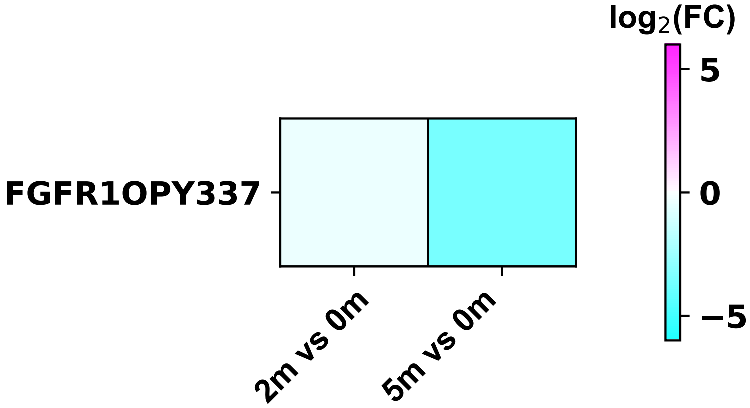

### flj00186_$^{y260}$_foldchange_m.pdf

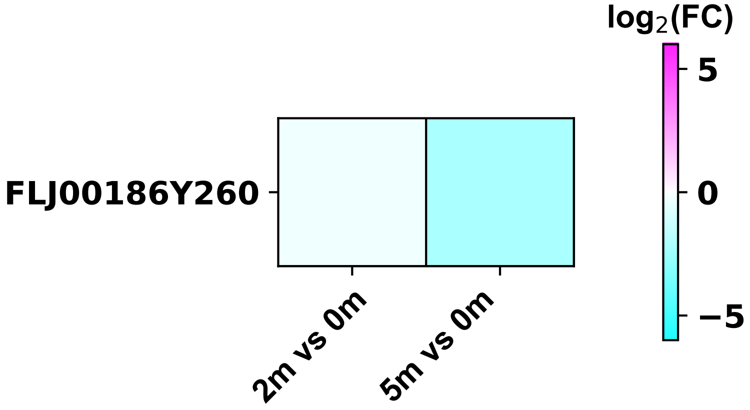

### flnb_$^{y904}$_foldchange_m.pdf

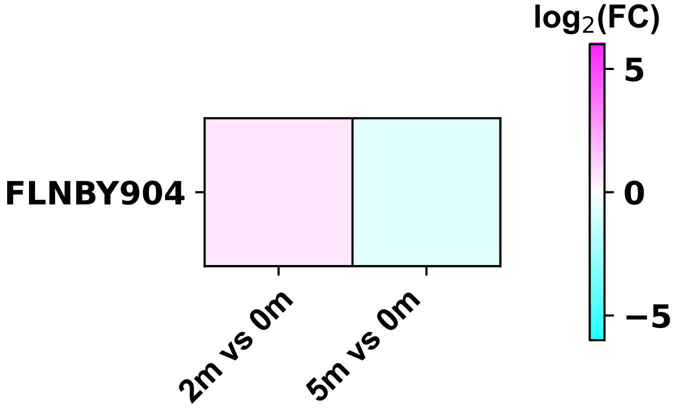

### flnb_$^{y2502}$_foldchange_m.pdf

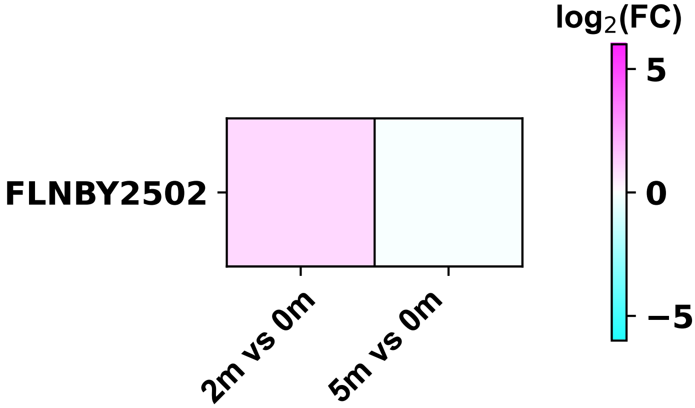

### fyb1_$^{y571}$_foldchange_m.pdf

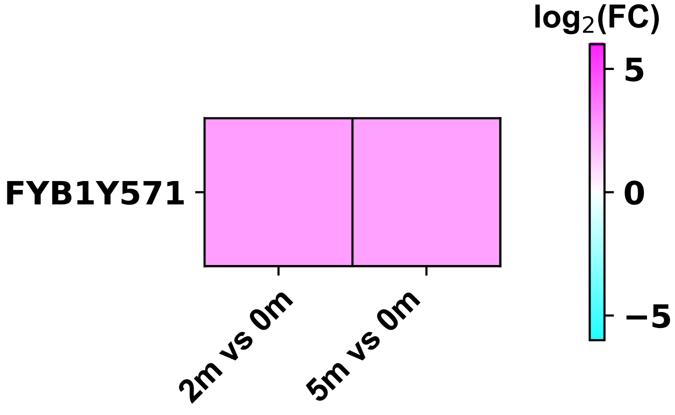

### g6pd_$^{y142}$_foldchange_m.pdf

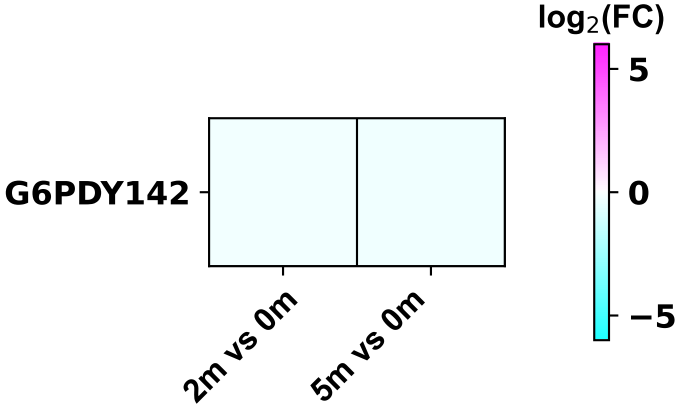

### g6pd_$^{y533}$_foldchange_m.pdf

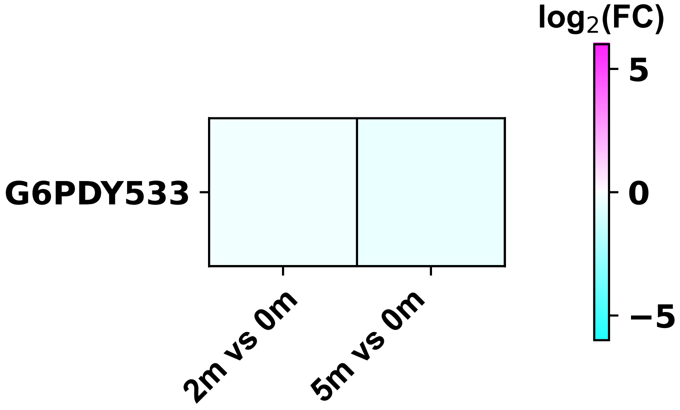

### gatad2b_$^{y317}$_foldchange_m.pdf

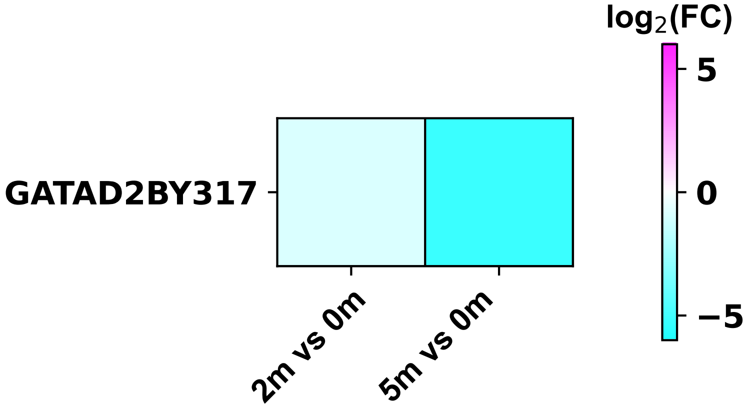

### gdi2_$^{y203}$_foldchange_m.pdf

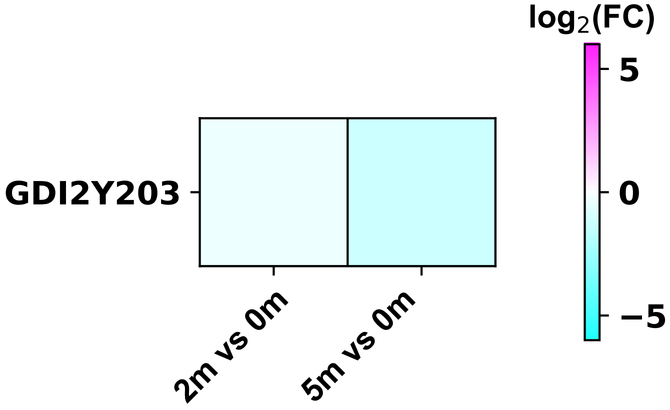

### git1_$^{y545}$_foldchange_m.pdf

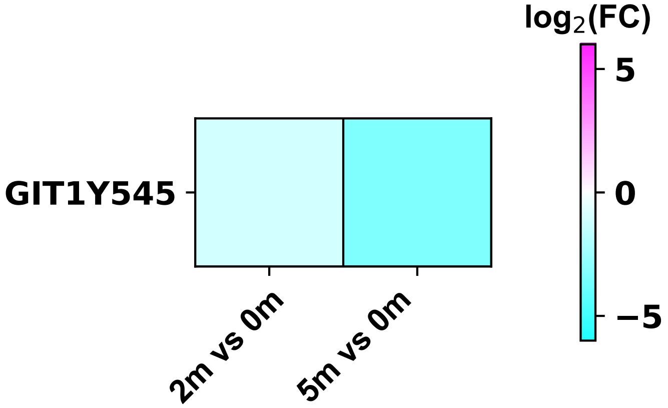

### git1_$^{y598}$_foldchange_m.pdf

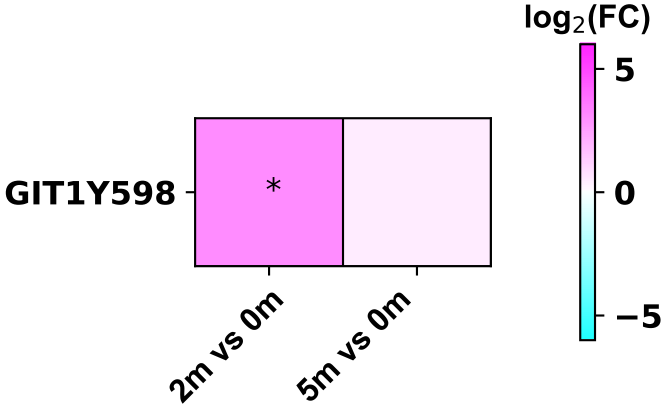

### gmds_$^{y323}$_foldchange_m.pdf

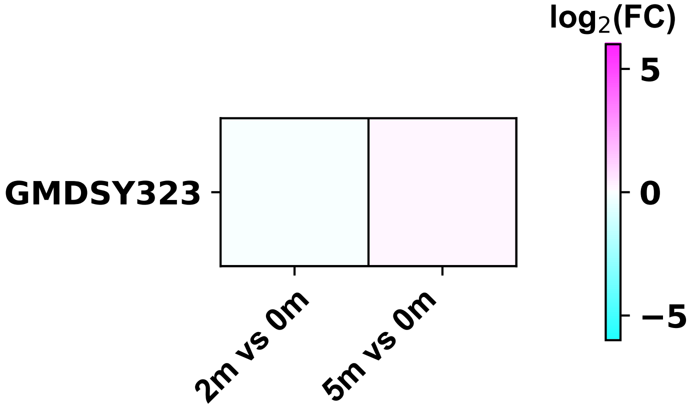

### golga5_$^{y54}$_foldchange_m.pdf

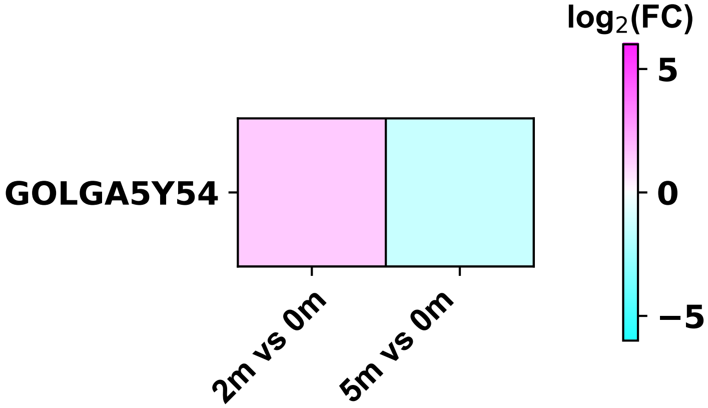

### gpsm3_$^{y108}$_foldchange_m.pdf

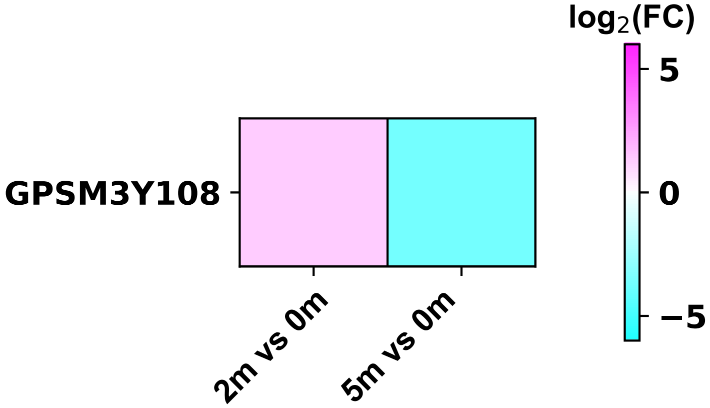

### grap2_$^{y45}$_foldchange_m.pdf

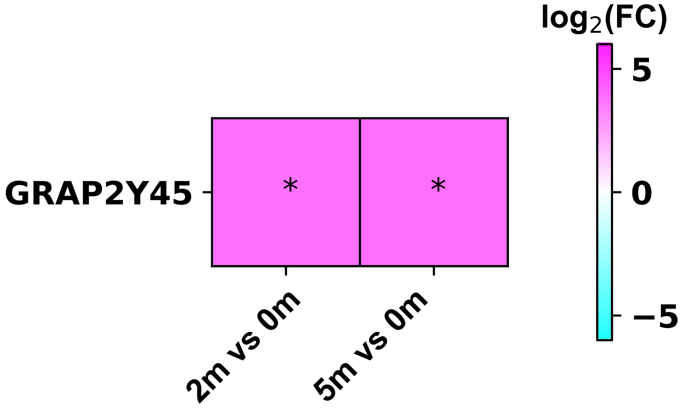

### grasp_$^{y237}$_foldchange_m.pdf

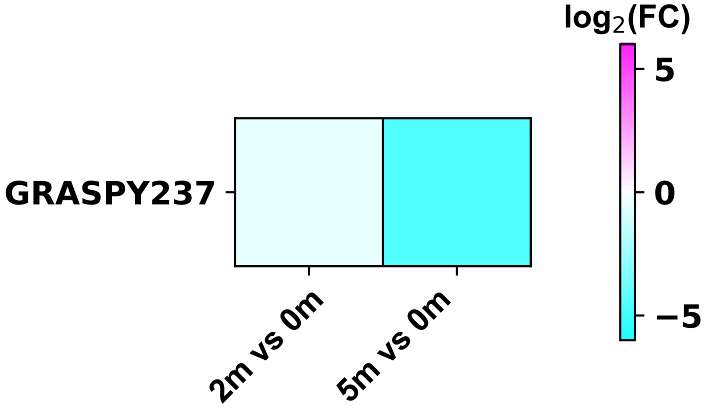

### gsk3b_$^{s215y216}$_foldchange_m.pdf

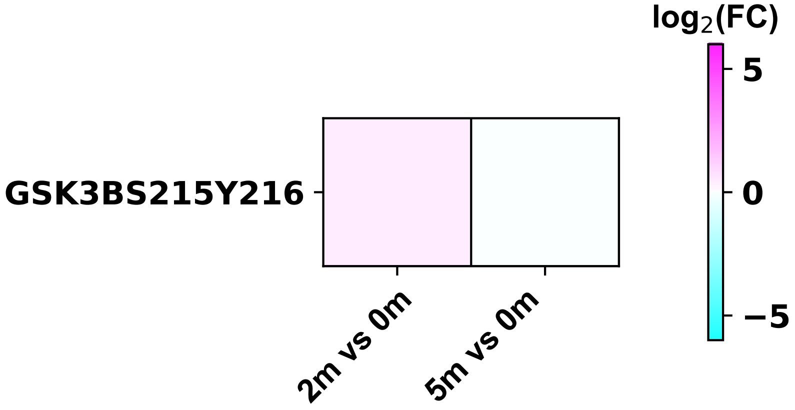
