## Supplementary figures and images for "CSF1R-CAR T cells induce CSF1R signalling and promote cancer cell growth"

### 28BBZ_CD19CarT_Raji_l_nmf.pdf

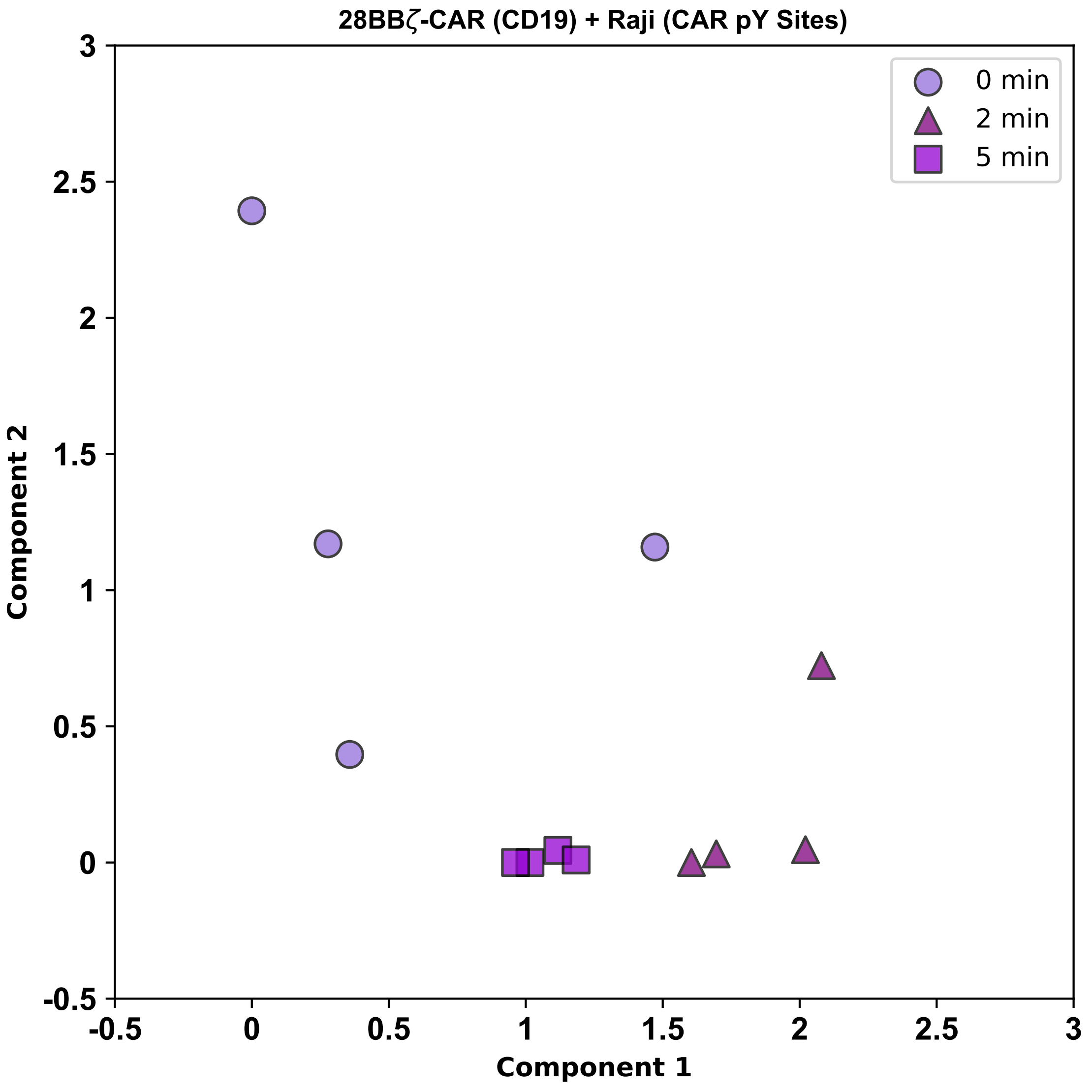

### 28BBZ_CD19CarT_Raji_l_pca.pdf

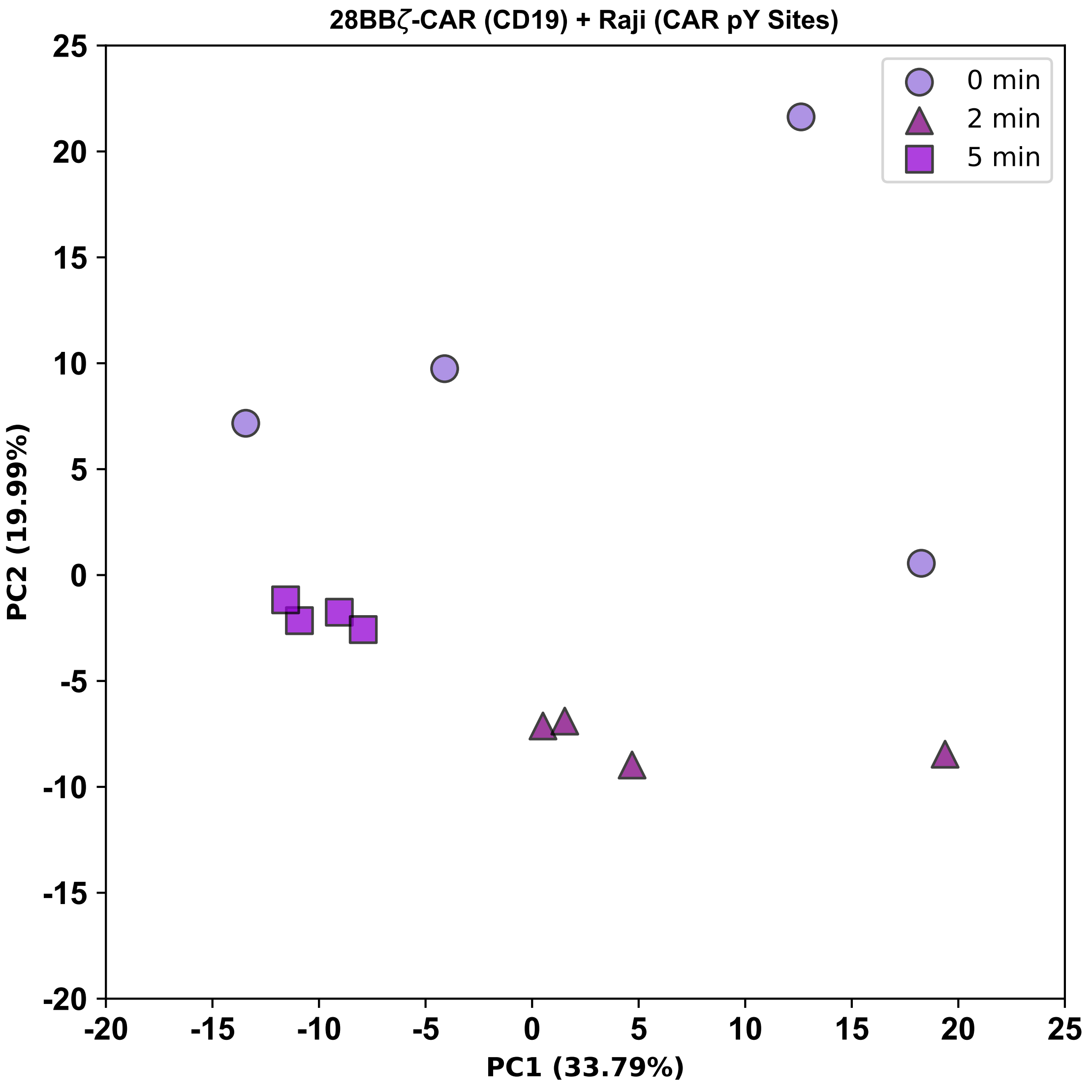

### abcf1_$^{y378}$_foldchange_m.pdf

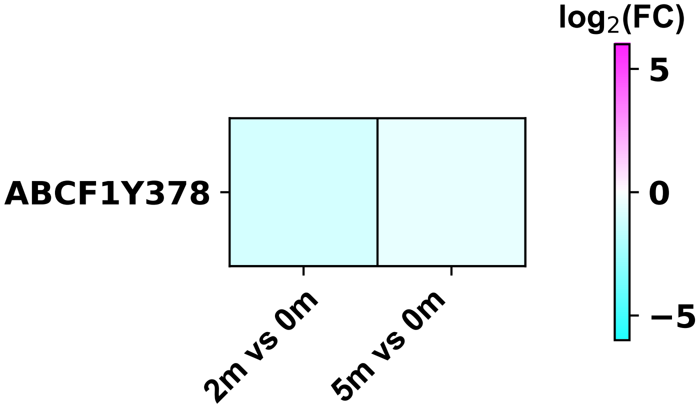

### abi1_$^{y213}$_foldchange_m.pdf

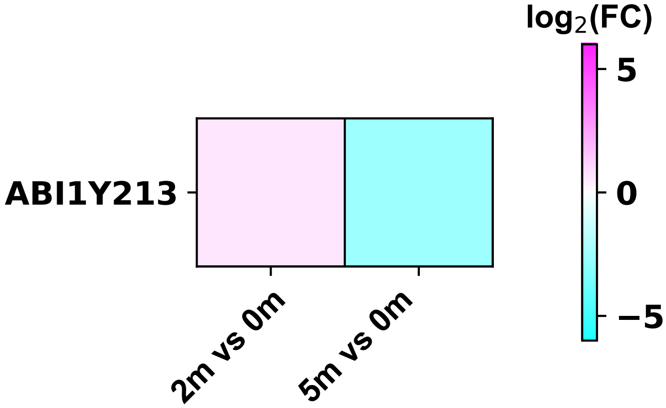

### abl1_$^{y439}$_foldchange_m.pdf

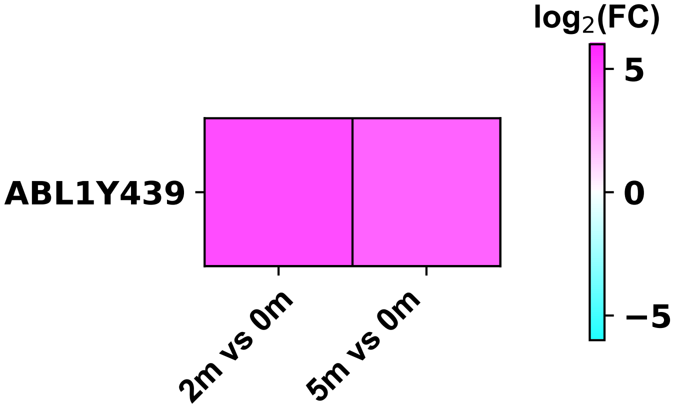

### acly_$^{y682}$_foldchange_m.pdf

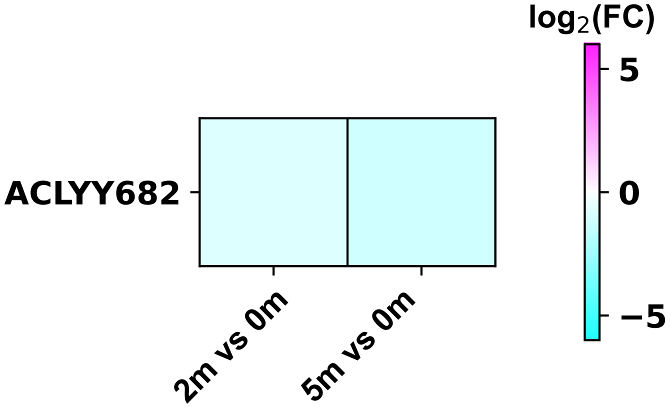

### acp1_$^{y132}$_foldchange_m.pdf

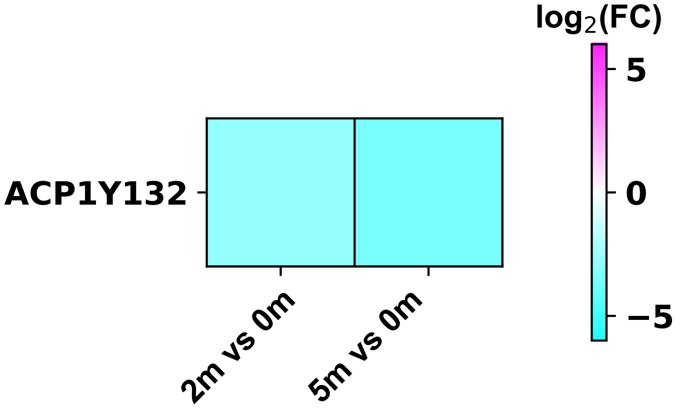

### actb_$^{y93}$_foldchange_m.pdf

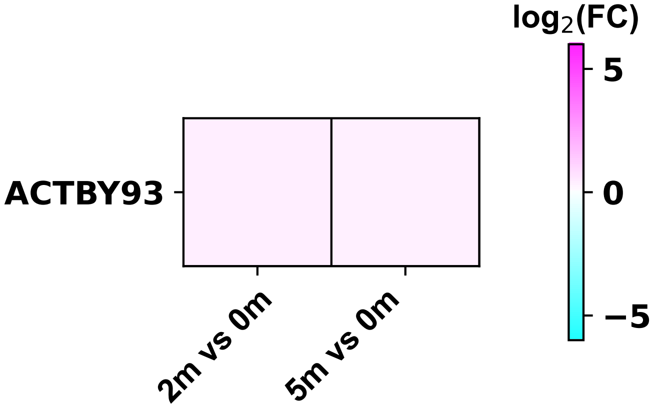

### actb_$^{y166}$_foldchange_m.pdf

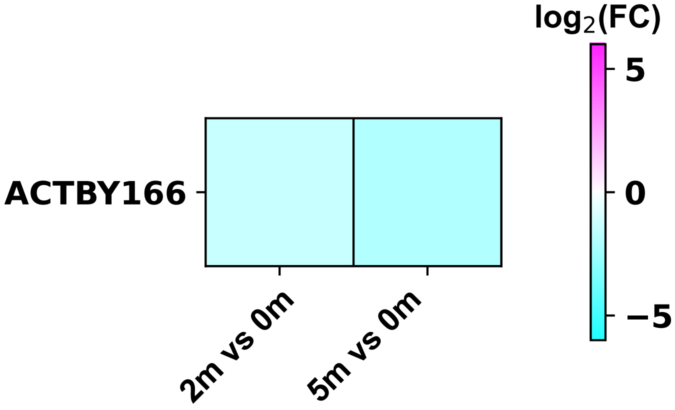

### actb_$^{y242}$_foldchange_m.pdf

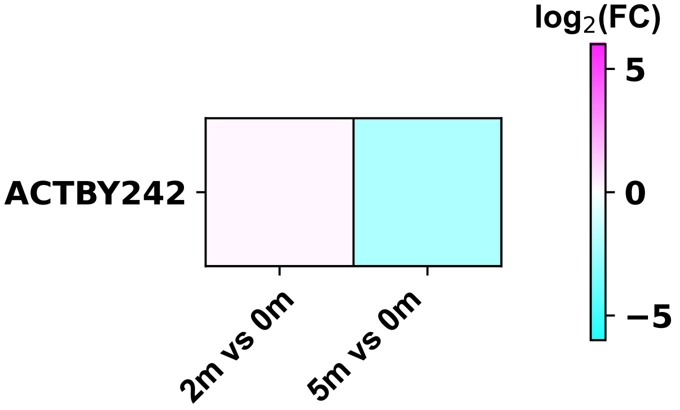

### actb_$^{y294}$_foldchange_m.pdf

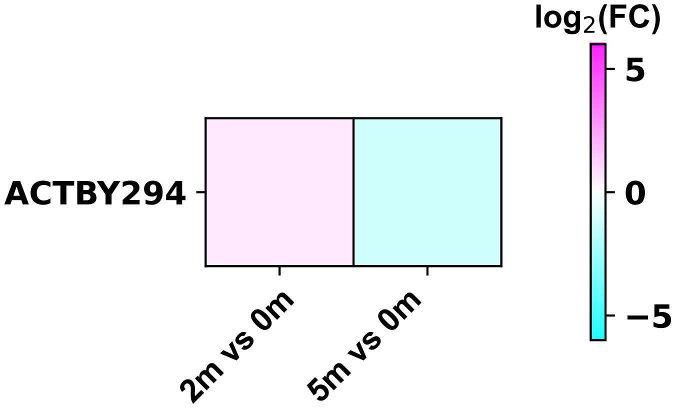

### actr3_$^{y16}$_foldchange_m.pdf

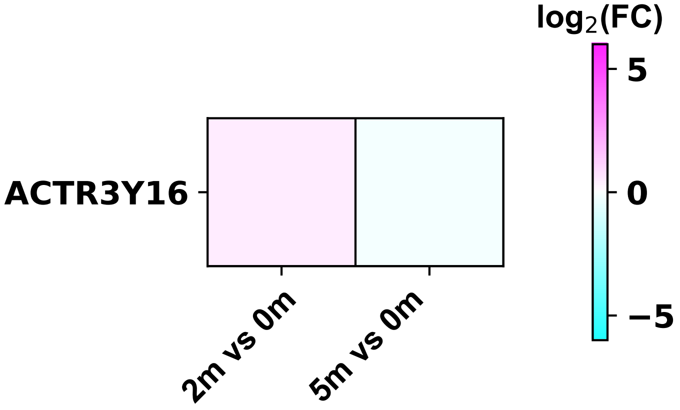

### actr10_$^{y4}$_foldchange_m.pdf

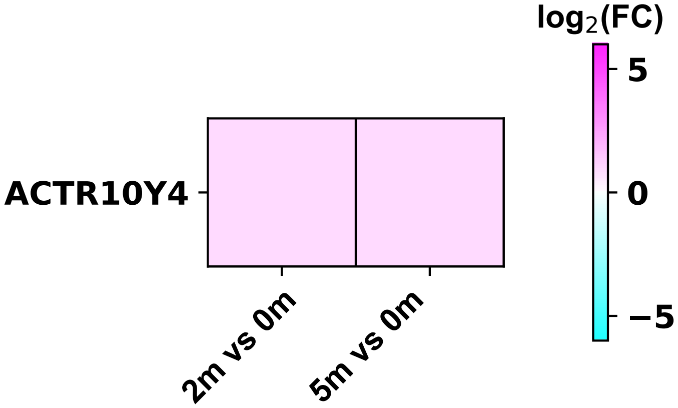

### ahcy_$^{y193}$_foldchange_m.pdf

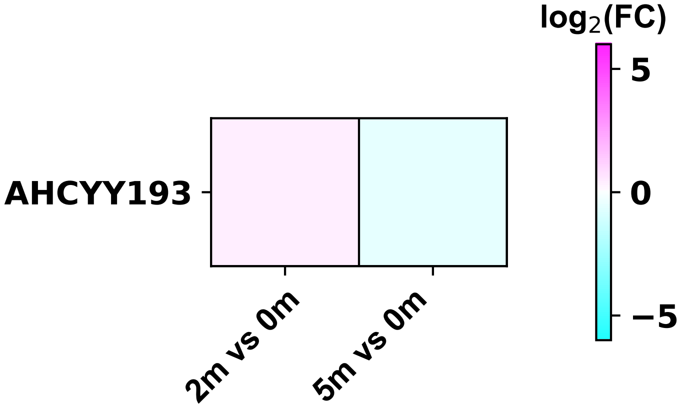

### aip_$^{y203}$_foldchange_m.pdf

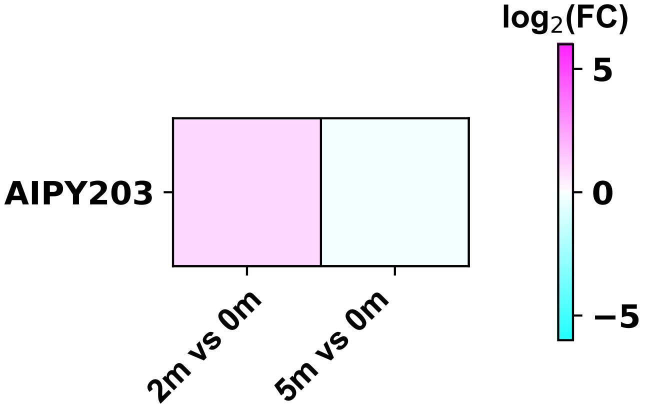

### aip_$^{y248}$_foldchange_m.pdf

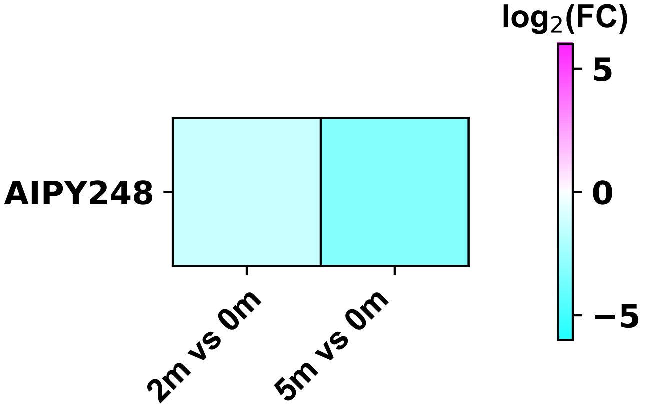

### ak6_$^{y100}$_foldchange_m.pdf

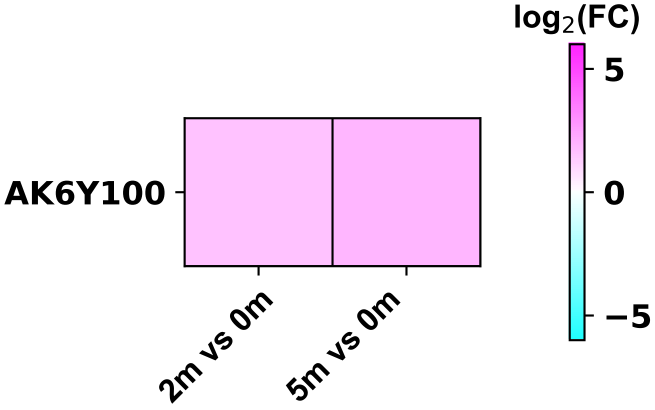

### akap11_$^{y424}$_foldchange_m.pdf

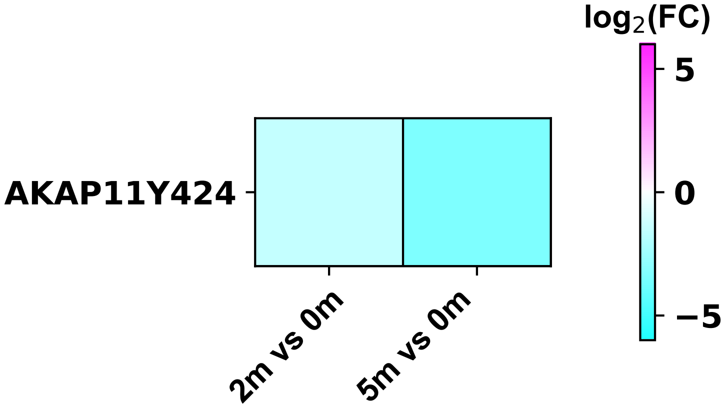

### aldoa_$^{y3}$_foldchange_m.pdf

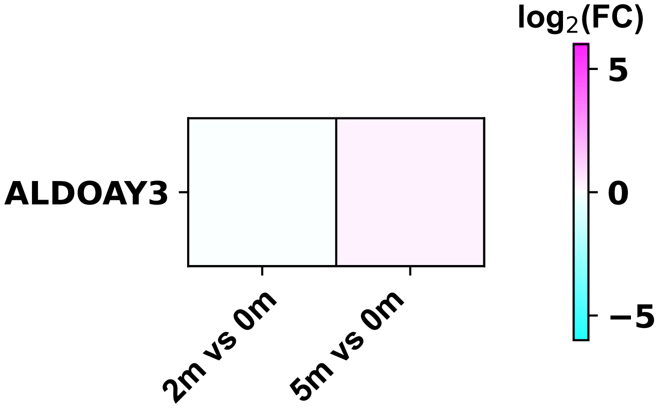

### alkbh8_$^{y379}$_foldchange_m.pdf

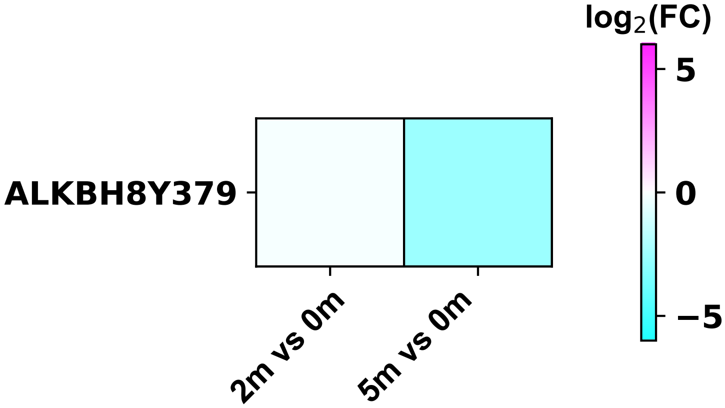

### ankrd12_$^{y37t40}$_foldchange_m.pdf

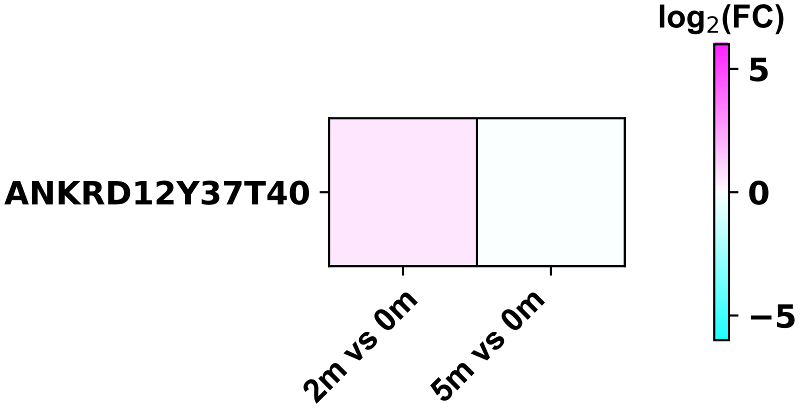

### ankrd39_$^{y65}$_foldchange_m.pdf

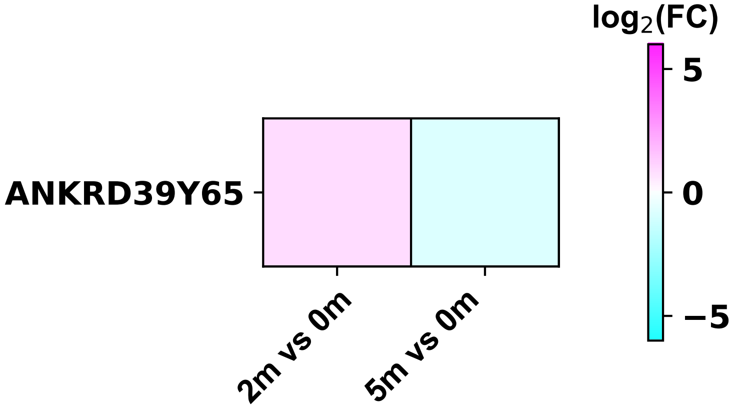

### anks1a_$^{y455}$_foldchange_m.pdf

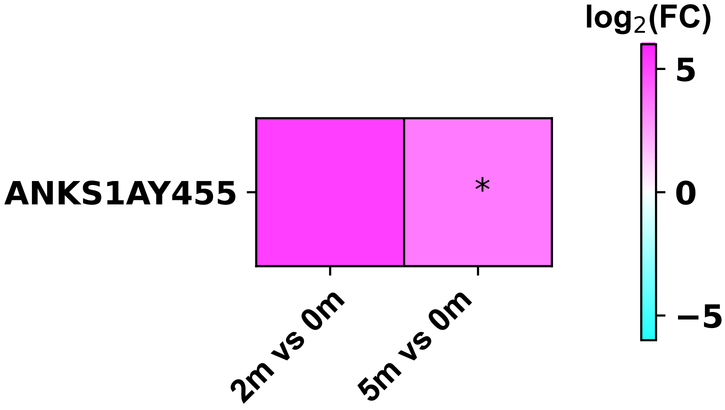

### anks1a_$^{y834}$_foldchange_m.pdf

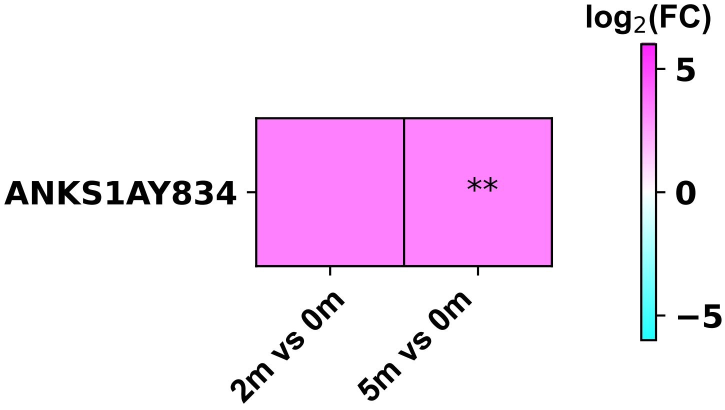

### anxa2_$^{y24}$_foldchange_m.pdf

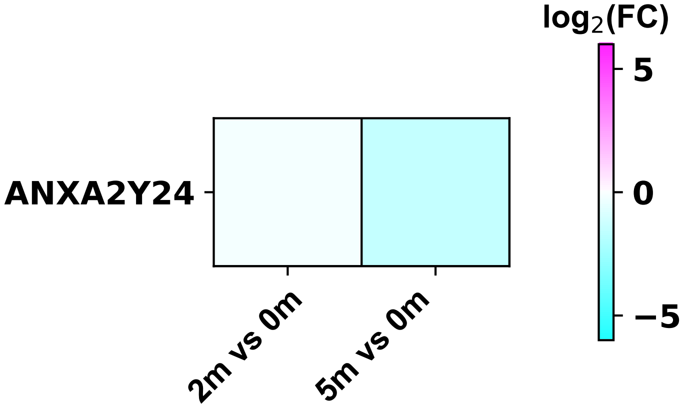

### anxa6_$^{y30}$_foldchange_m.pdf

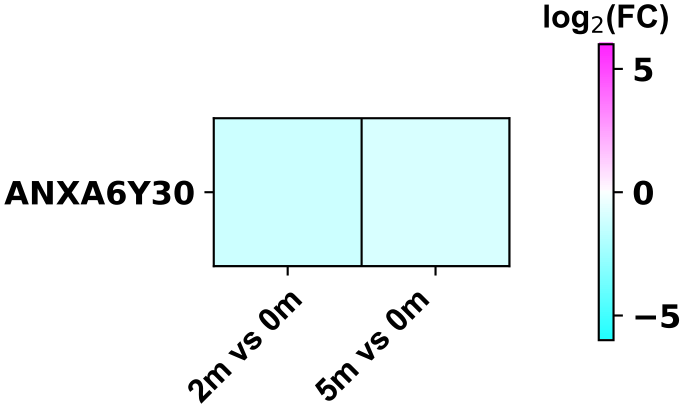

### ap2b1_$^{y897}$_foldchange_m.pdf

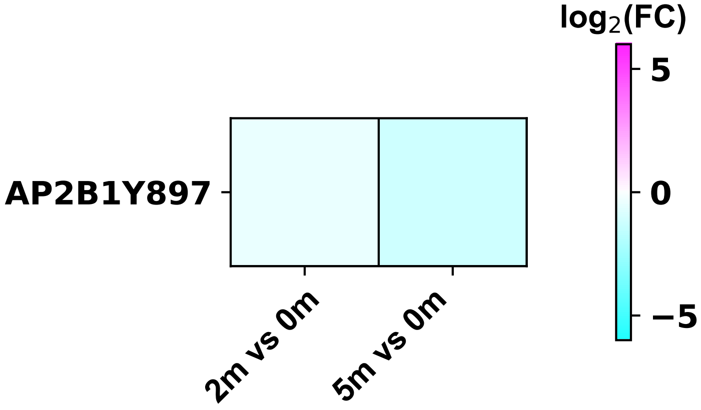

### arap1_$^{y497}$_foldchange_m.pdf

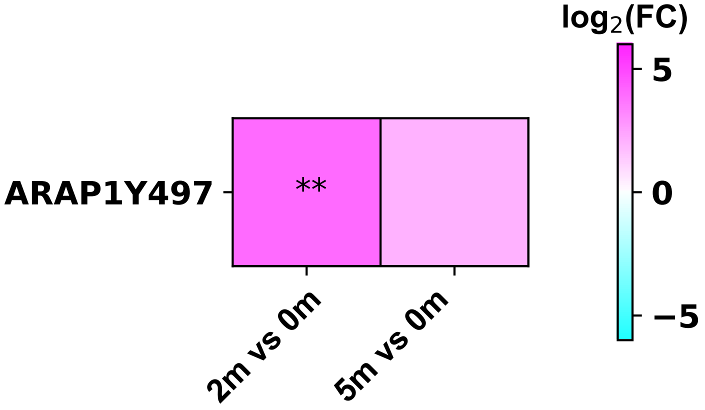

### arhgap27_$^{y28}$_foldchange_m.pdf

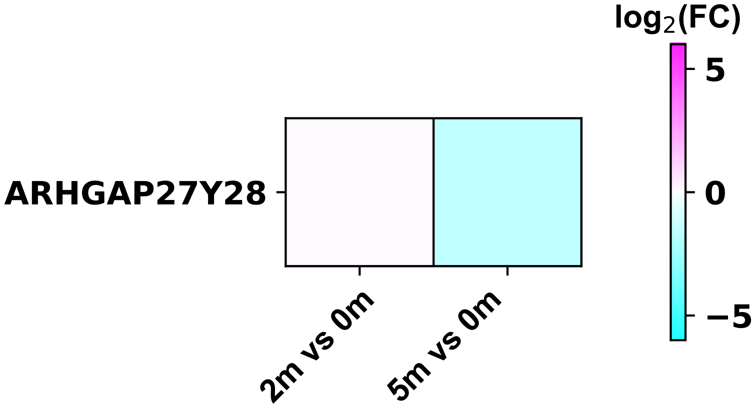

### arhgap27_$^{y228}$_foldchange_m.pdf

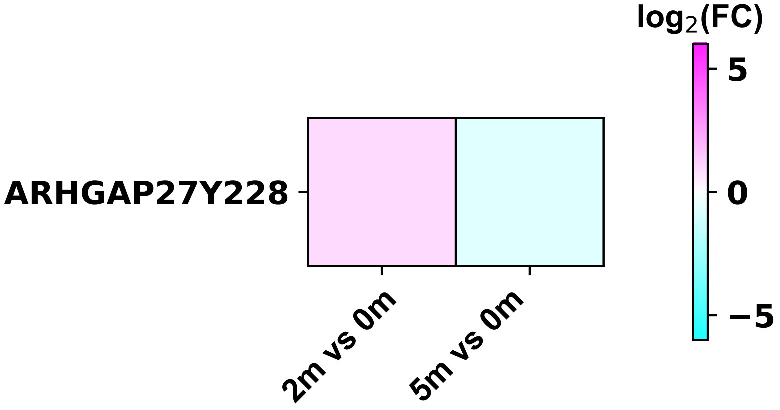

### arhgap30_$^{y816}$_foldchange_m.pdf

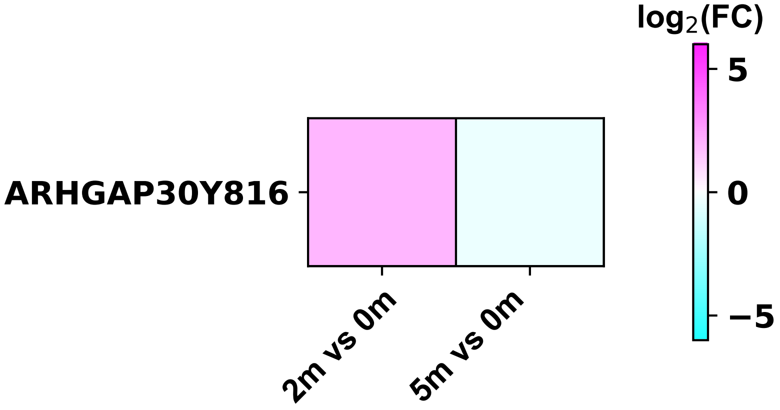

### arhgap35_$^{y1087}$_foldchange_m.pdf

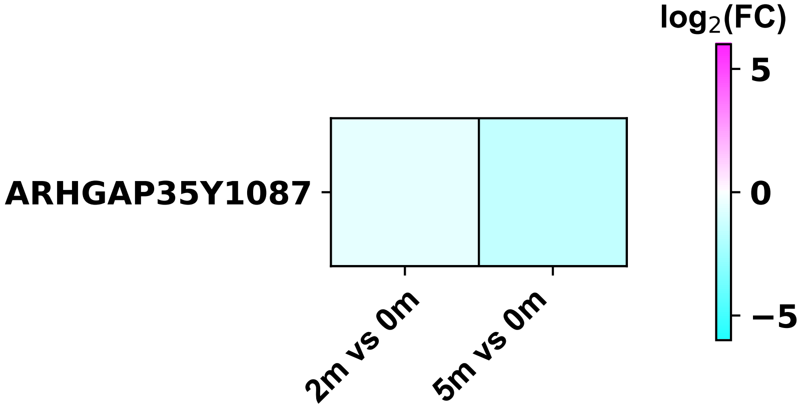

### arhgap35_$^{y1105}$_foldchange_m.pdf

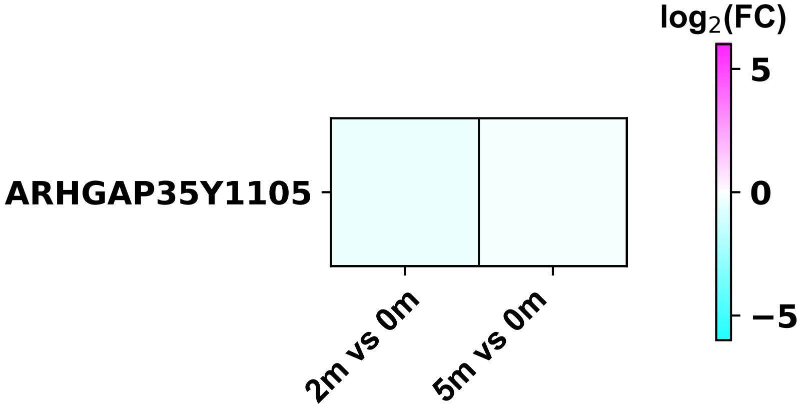

### arrb2_$^{y404}$_foldchange_m.pdf

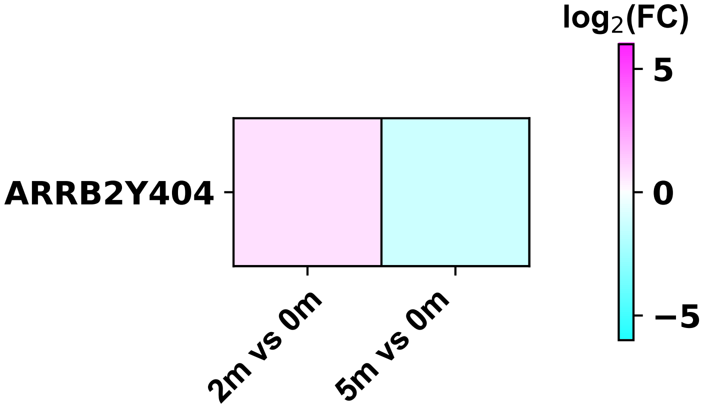

### atg2b_$^{y2012}$_foldchange_m.pdf

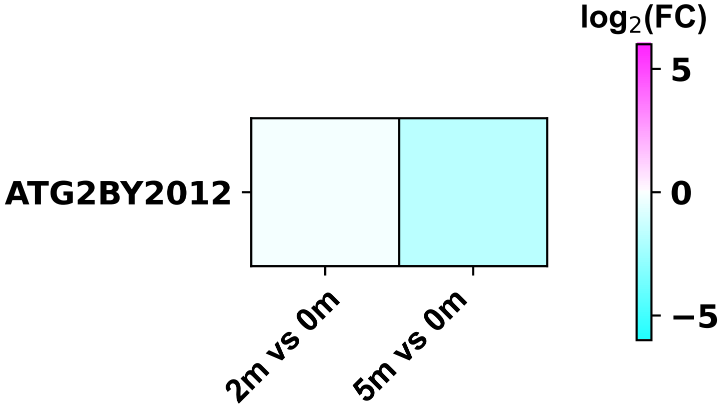

### atg4b_$^{y8}$_foldchange_m.pdf

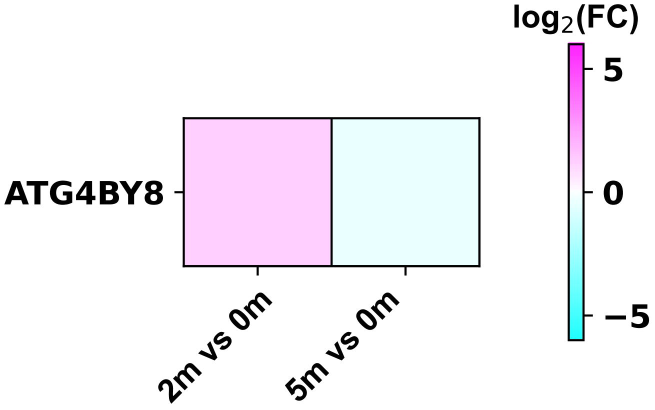

### atp1a1_$^{y260}$_foldchange_m.pdf

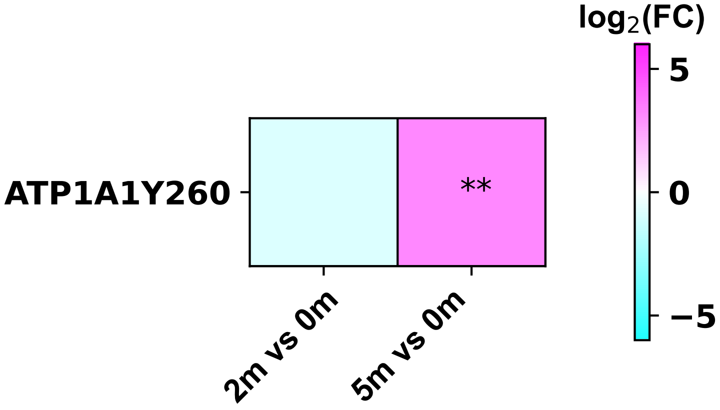

### atp11a_$^{y233}$_foldchange_m.pdf

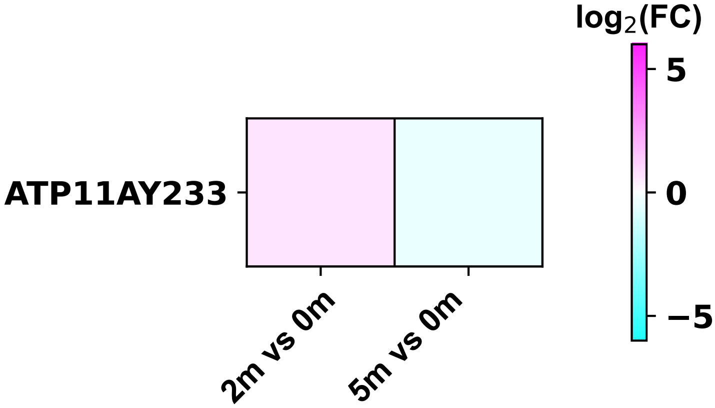

### atxn2_$^{y471}$_foldchange_m.pdf

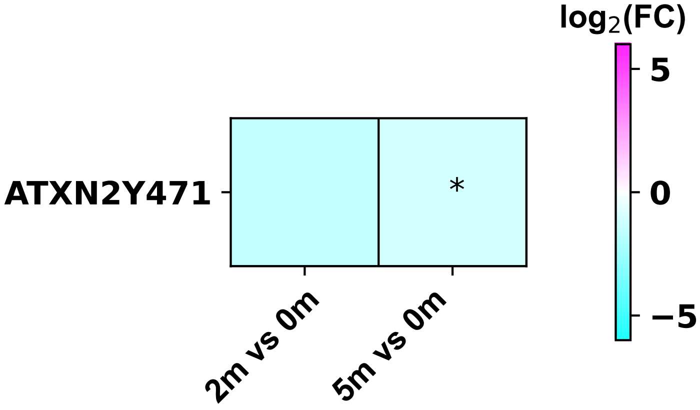

### atxn2l_$^{y349}$_foldchange_m.pdf

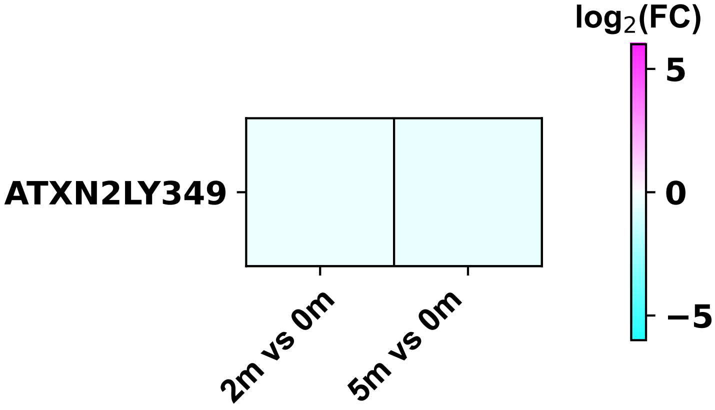

### bcr_$^{y177}$_foldchange_m.pdf

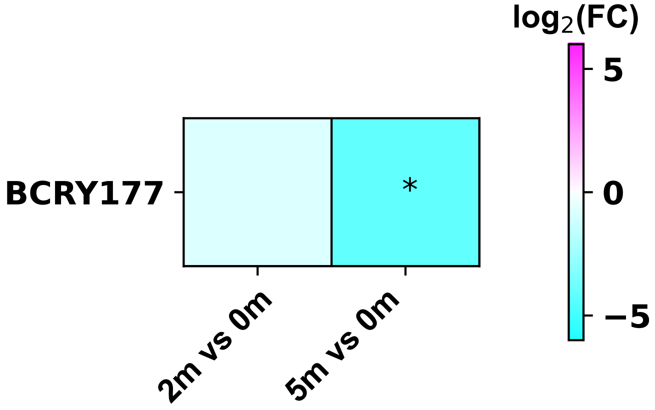

### bcr_$^{y246}$_foldchange_m.pdf

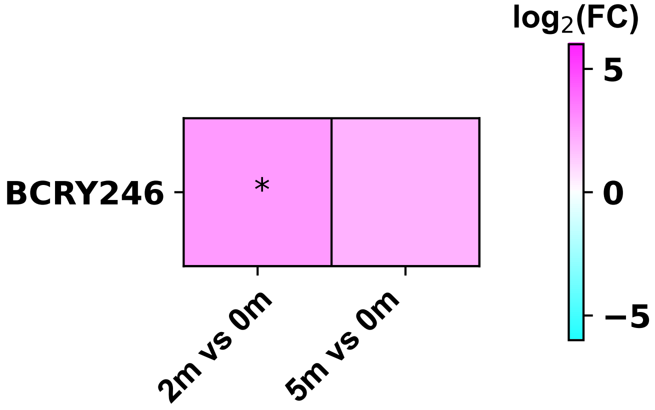

### blk_$^{y389}$_foldchange_m.pdf

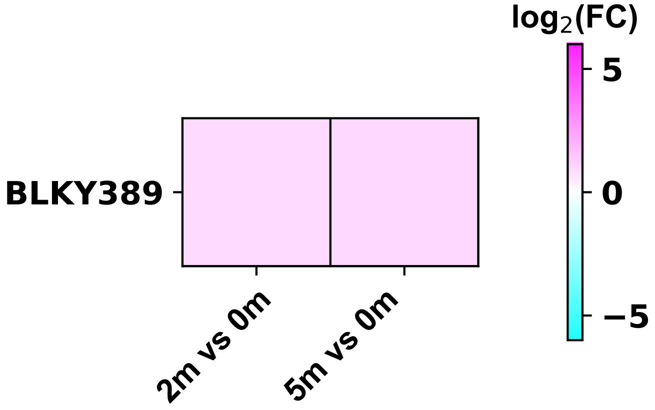

### bysl_$^{y49}$_foldchange_m.pdf

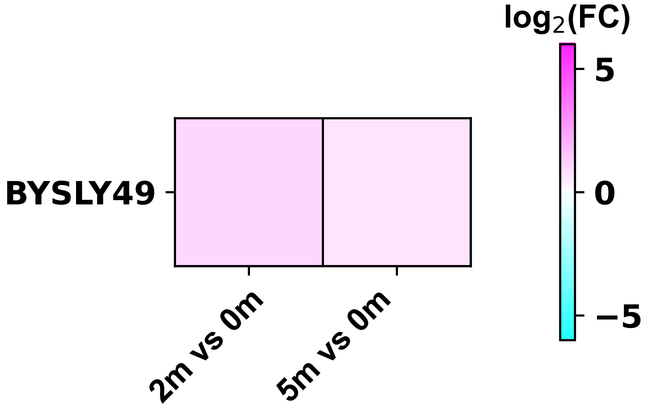

### c9orf78_$^{y147}$_foldchange_m.pdf

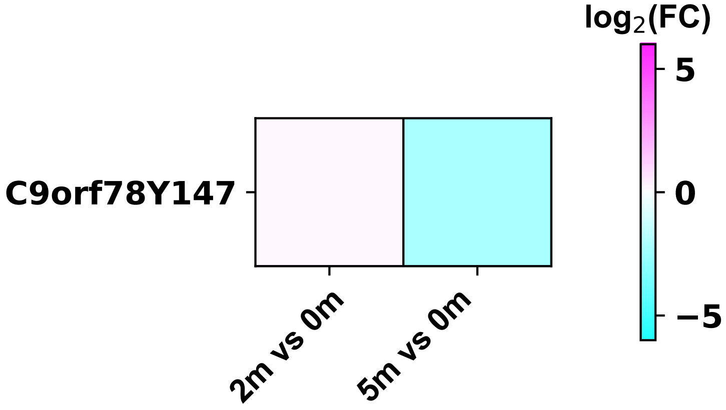

### cacybp_$^{y71}$_foldchange_m.pdf

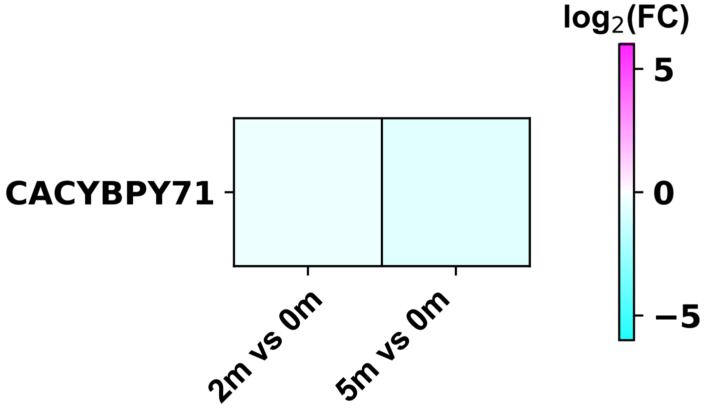

### calm2_$^{y100}$_foldchange_m.pdf

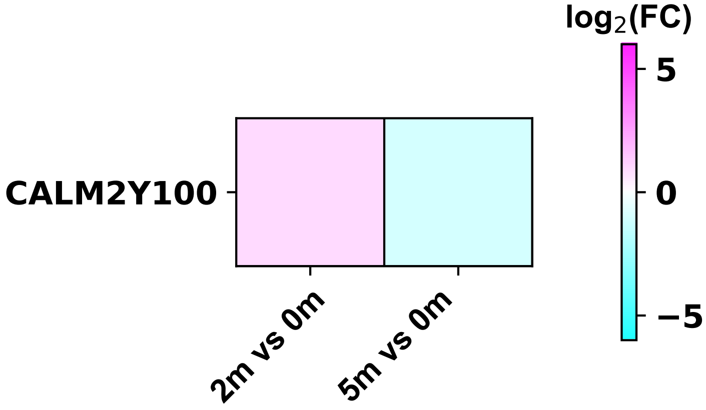

### calu_$^{y47}$_foldchange_m.pdf

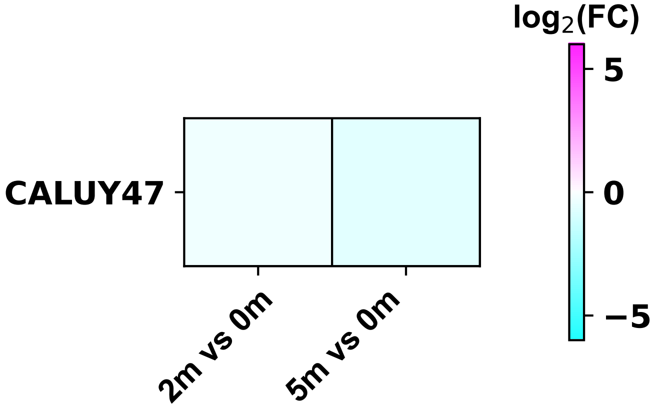

### capzb_$^{y232}$_foldchange_m.pdf

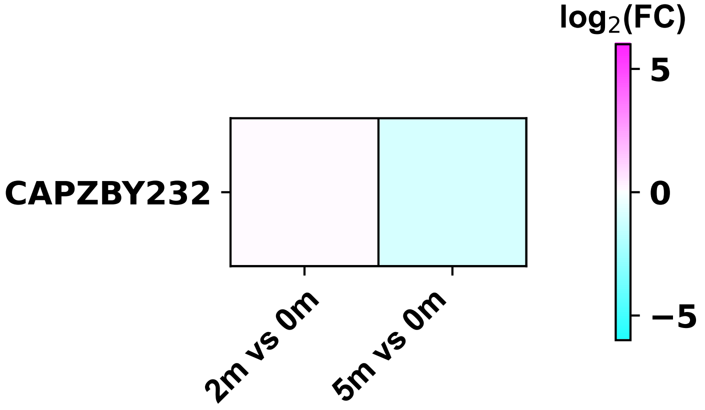

### carj_thp1_growth.pdf

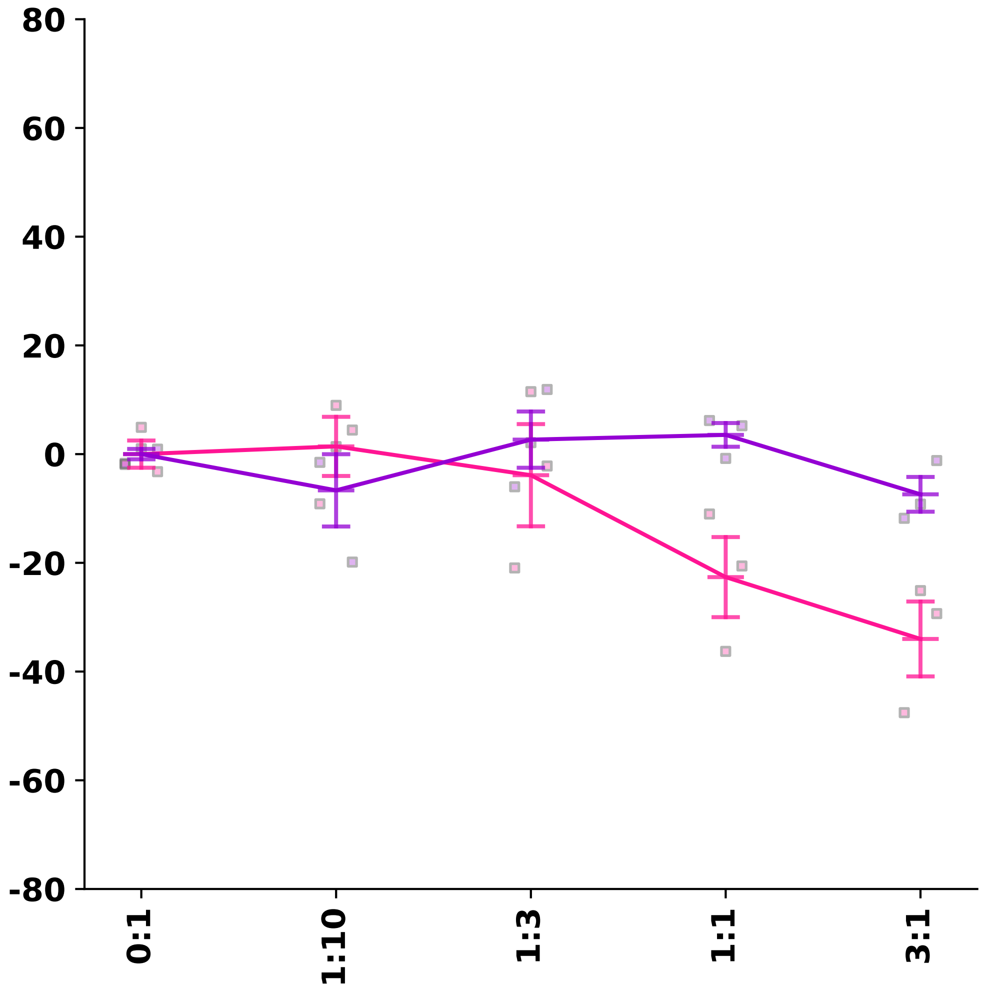

### carp_thp1_growth.pdf

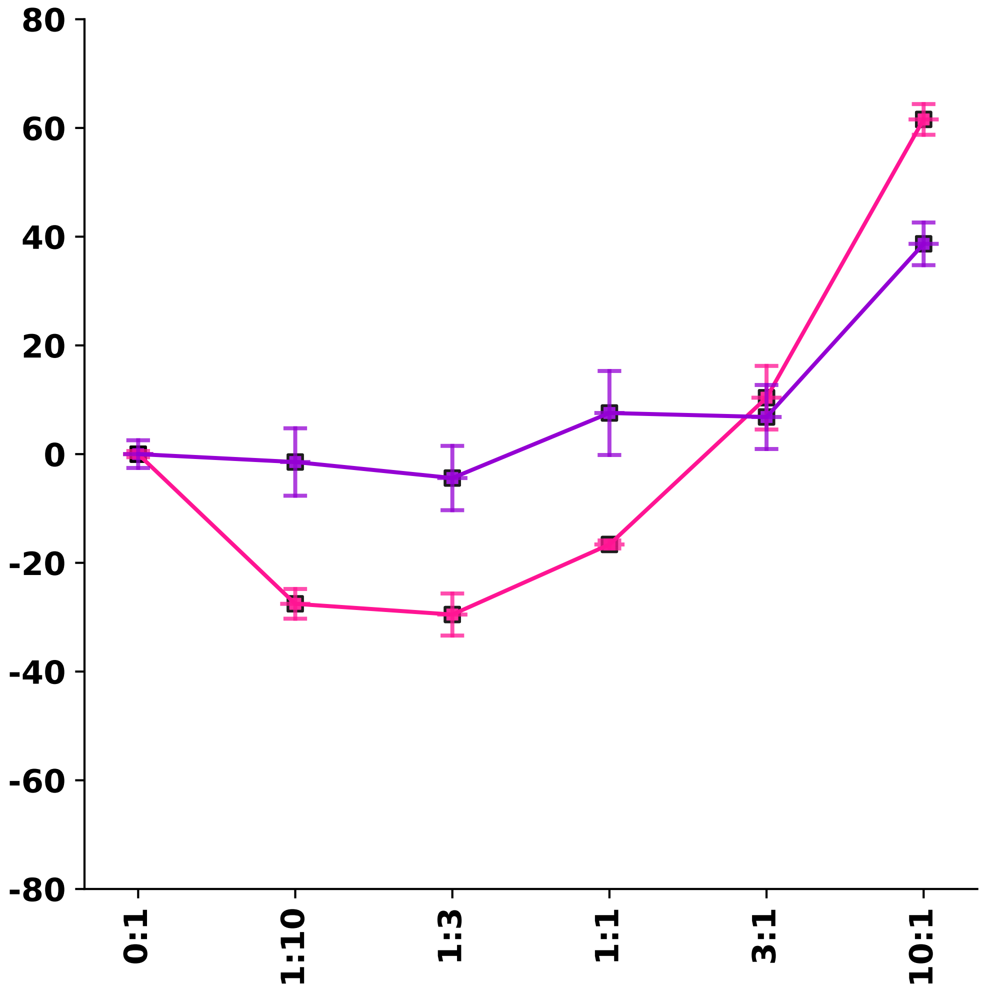

### carpj_thp1_growth.pdf

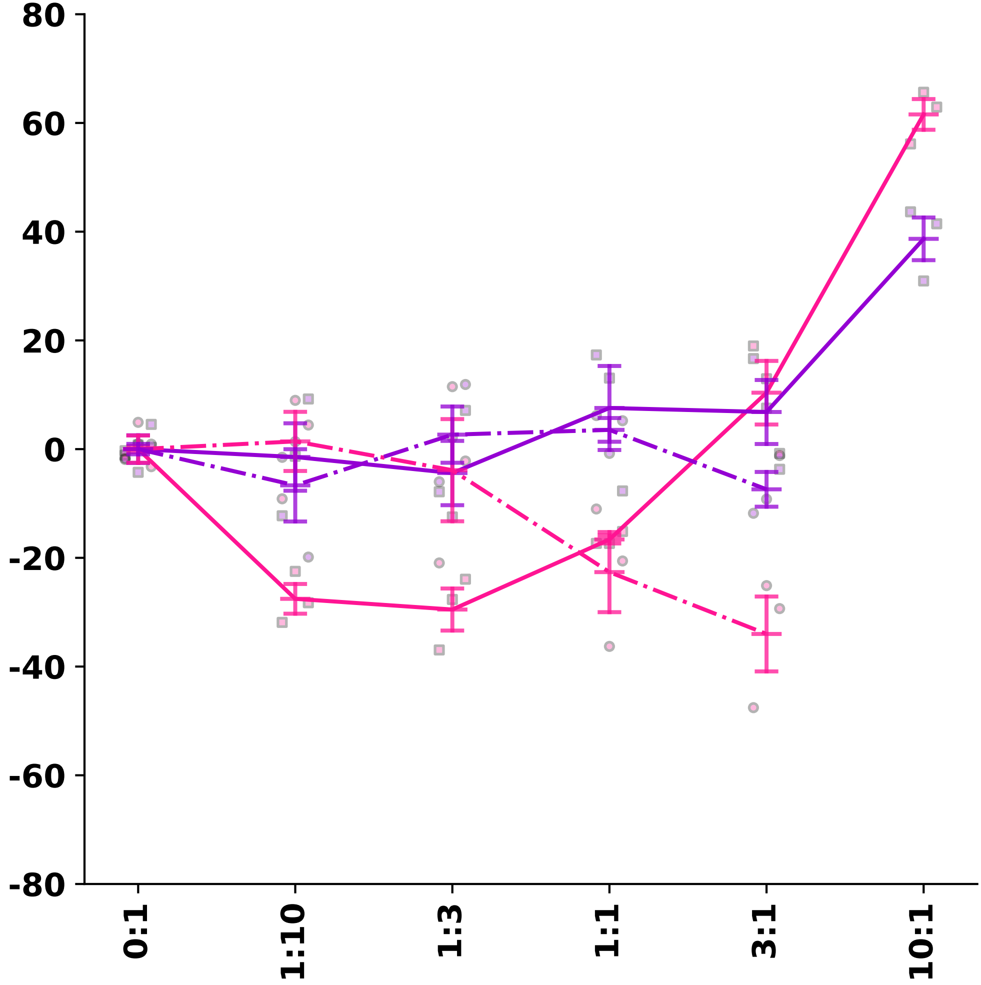

### cars_$^{y53}$_foldchange_m.pdf

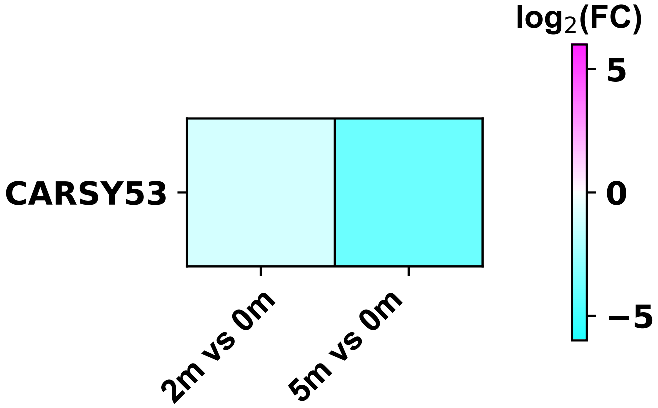

### caskin2_$^{y253}$_foldchange_m.pdf

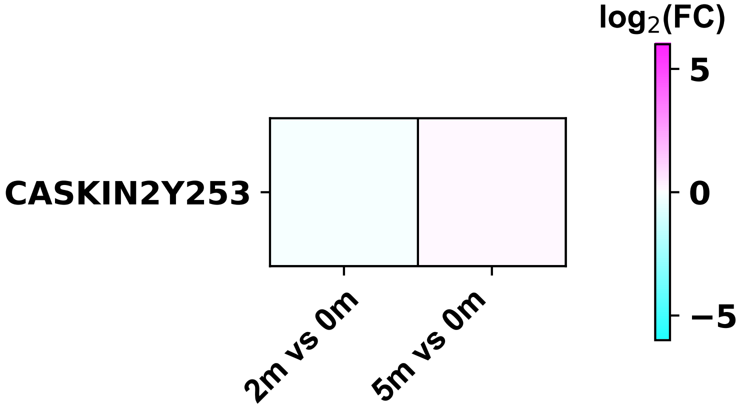

### cbl_$^{s667y674}$_foldchange_m.pdf

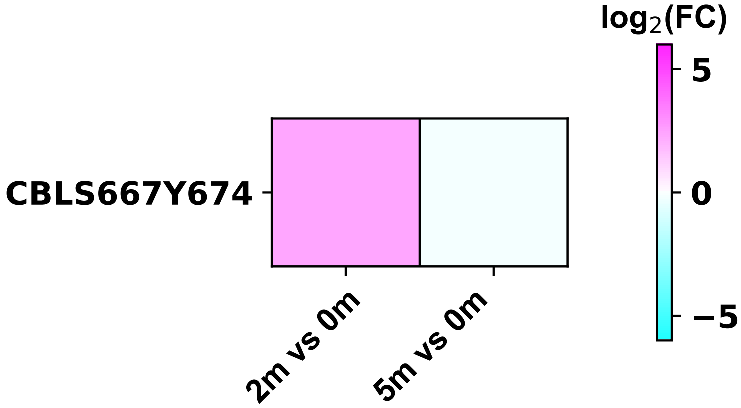

### cbl_$^{y674}$_foldchange_m.pdf

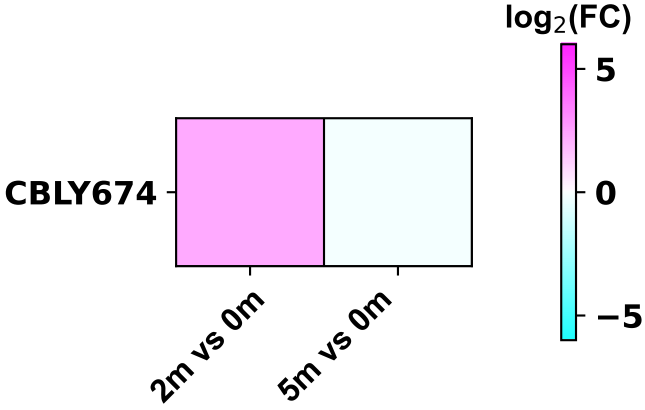

### ccdc14_$^{y564}$_foldchange_m.pdf

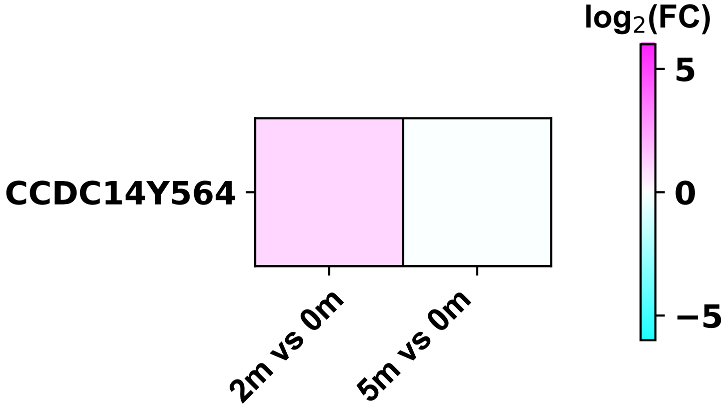

### cd2ap_$^{y361}$_foldchange_m.pdf

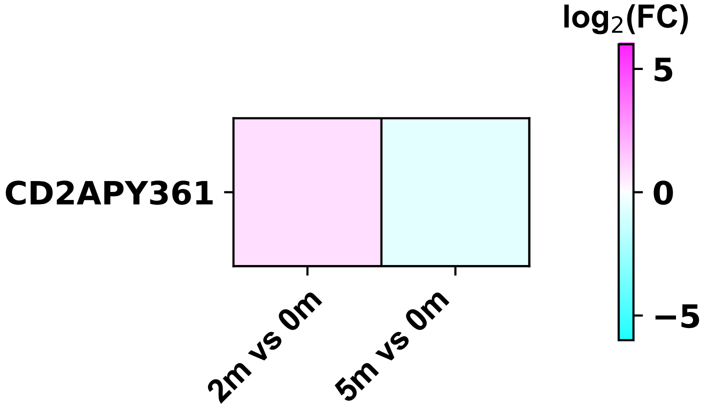

### cd3d_$^{y160}$_foldchange_m.pdf

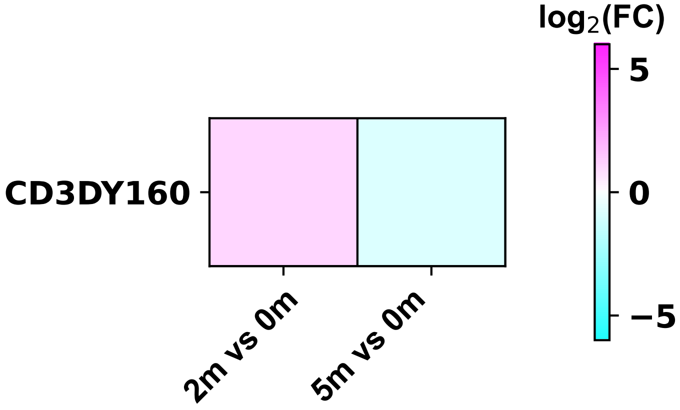

### cd3e_$^{y188}$_foldchange_m.pdf

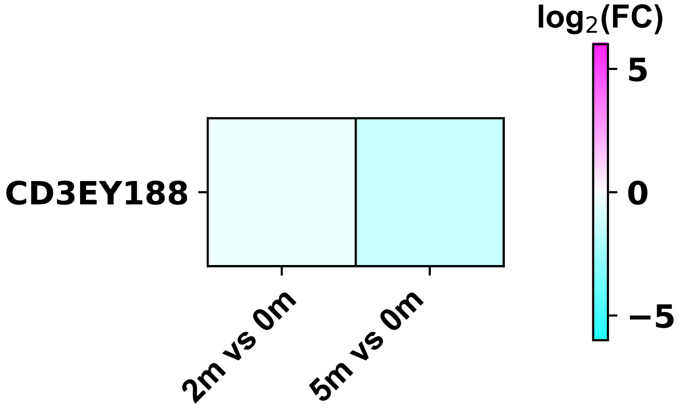

### cd3e_$^{y199}$_foldchange_m.pdf

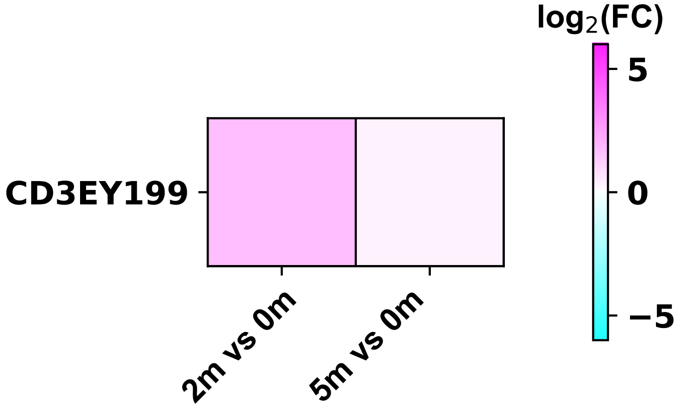

### cd3g_$^{y160}$_foldchange_m.pdf

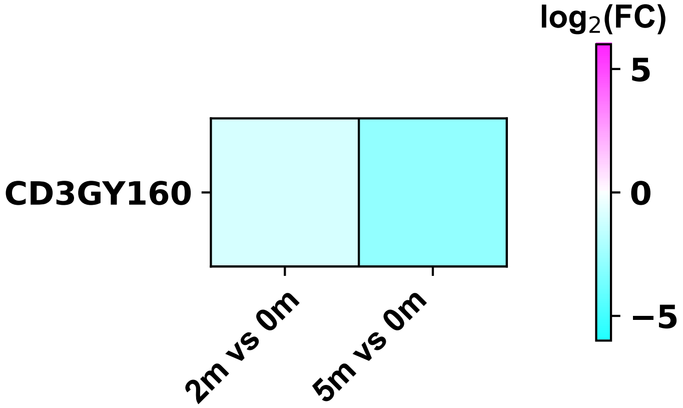

### cd3g_$^{y171}$_foldchange_m.pdf

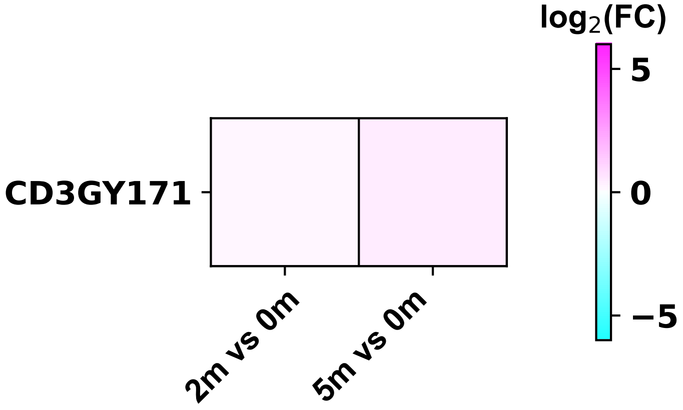

### cd5_$^{y453}$_foldchange_m.pdf

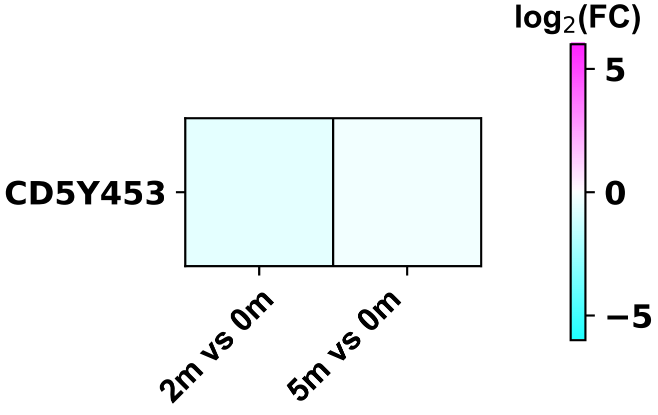

### cd7_$^{y222}$_foldchange_m.pdf

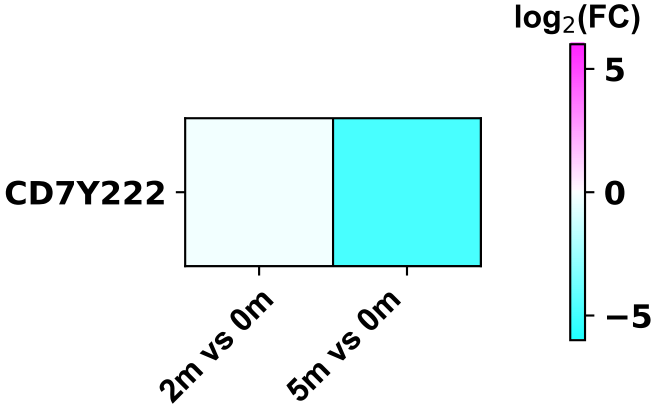

### cd19cargen3_$^{s189y191}$_foldchange_m.pdf

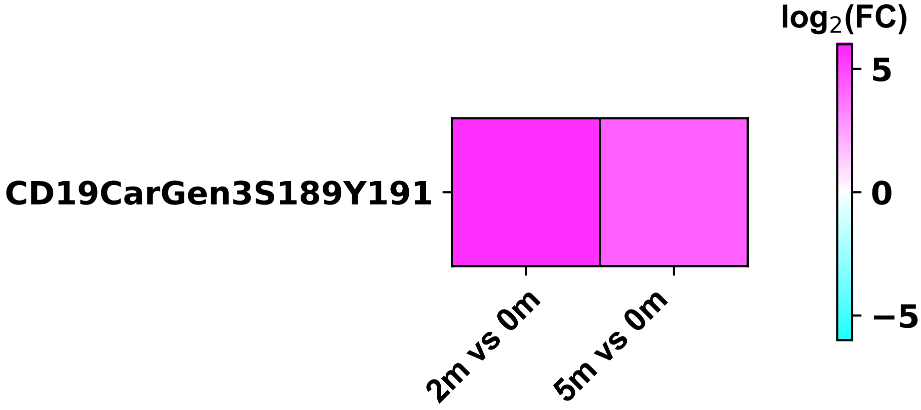

### cd19cargen3_$^{y83}$_foldchange_m.pdf

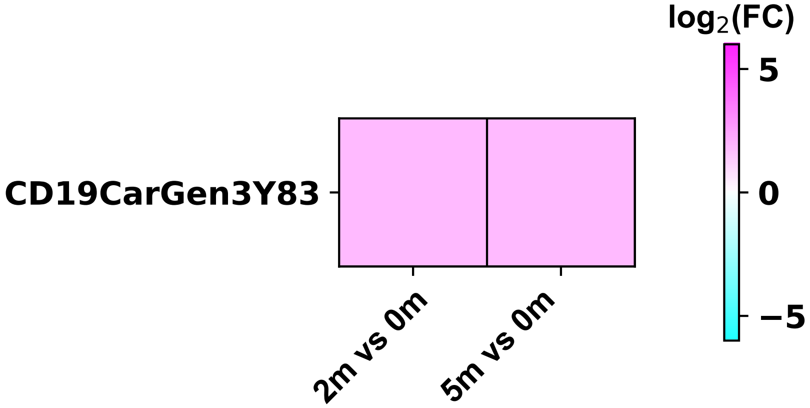

### cd19cargen3_$^{y111}$_foldchange_m.pdf

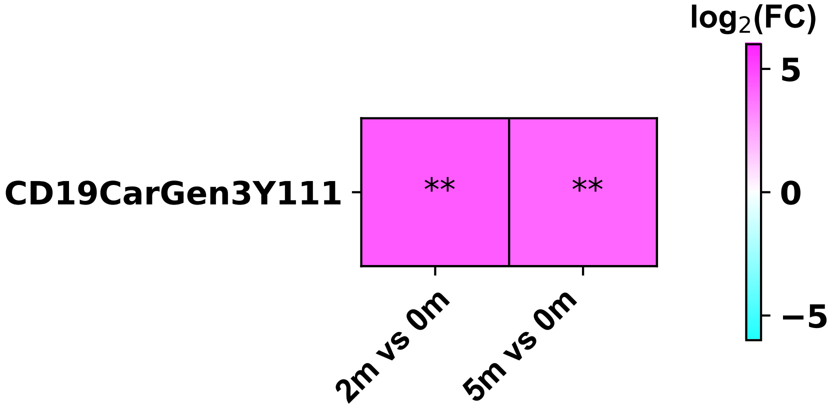

### cd19cargen3_$^{y123}$_foldchange_m.pdf

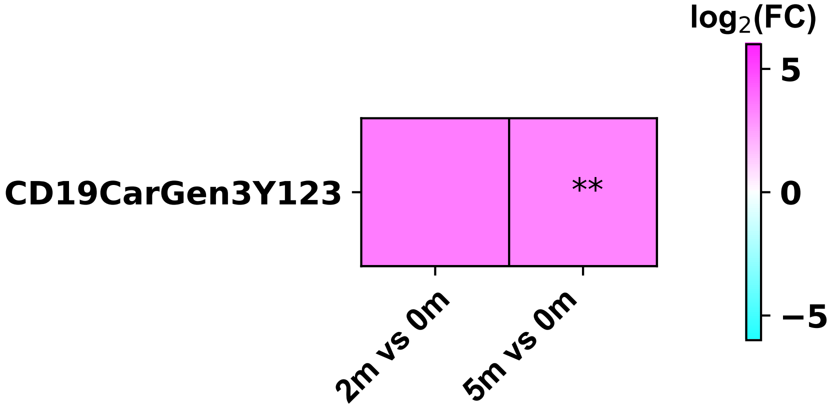

### cd19cargen3_$^{y142}$_foldchange_m.pdf

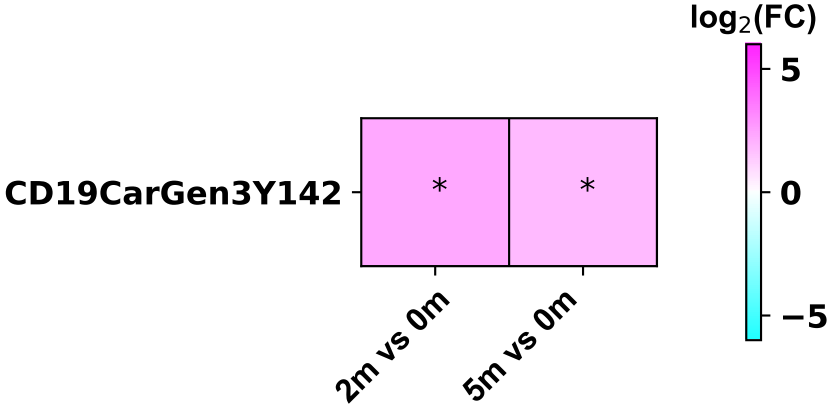

### cd19cargen3_$^{y153}$_foldchange_m.pdf

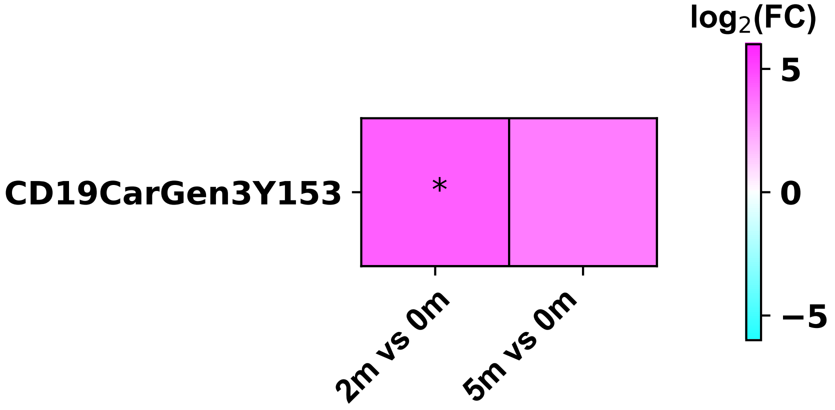

### cd19cargen3_$^{y191t195}$_foldchange_m.pdf

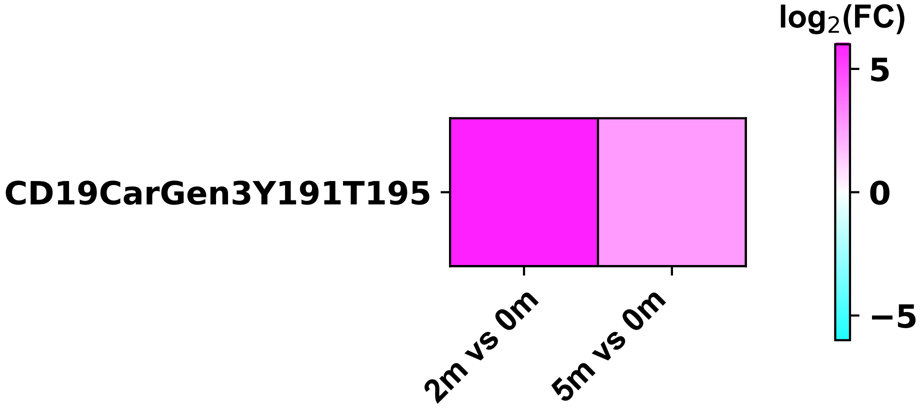

### cd19cargen3_$^{y191}$_foldchange_m.pdf

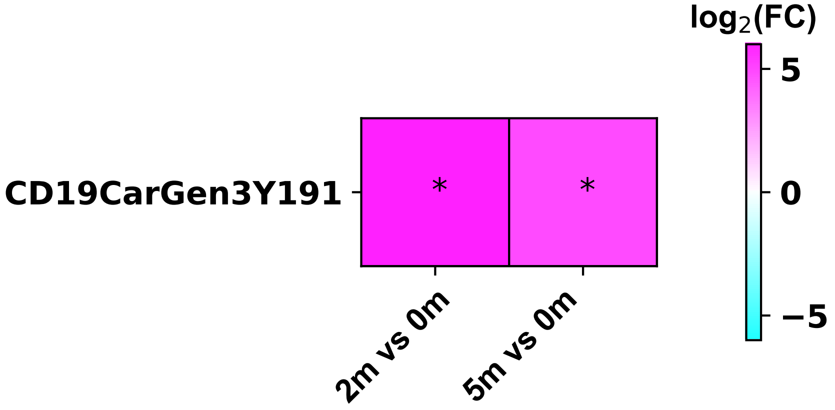

### cd84_$^{y262}$_foldchange_m.pdf

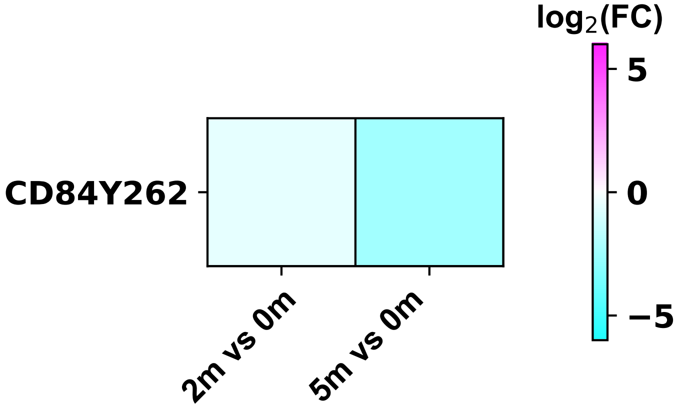

### cd84_$^{y279}$_foldchange_m.pdf

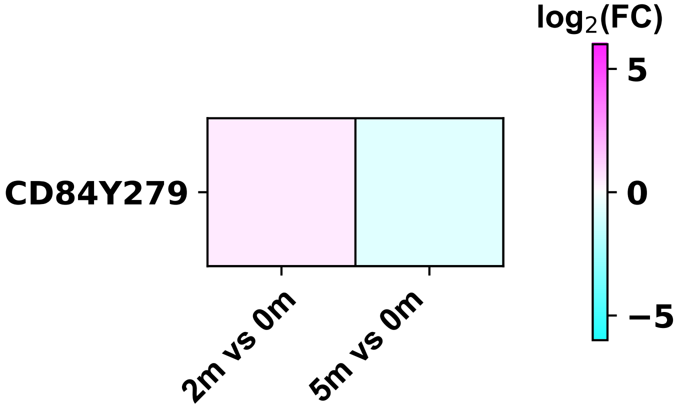

### cdk1_$^{t14y15}$_foldchange_m.pdf

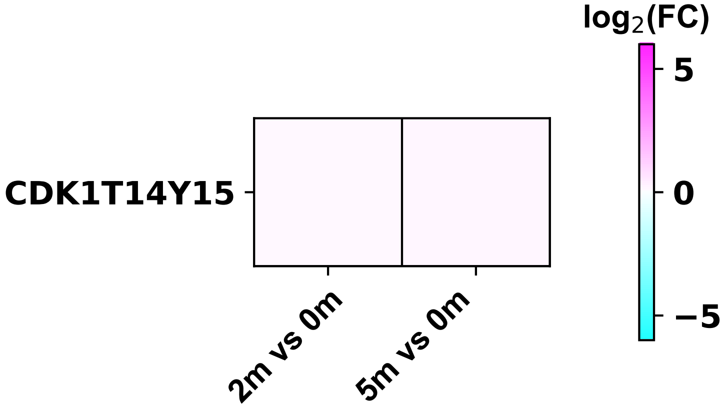

### cdk1_$^{y15}$_foldchange_m.pdf

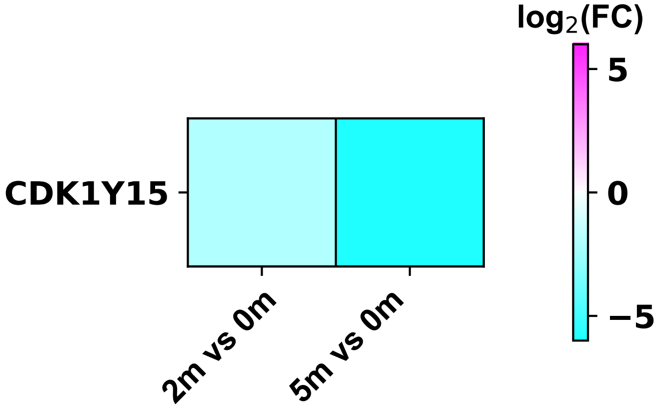

### cdk2_$^{y15}$_foldchange_m.pdf

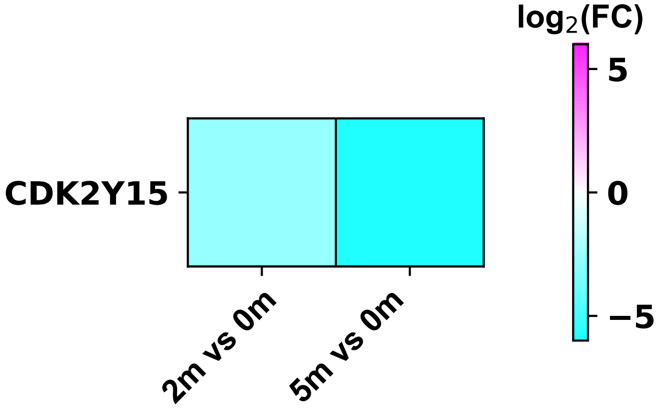

### cdk5_$^{y15}$_foldchange_m.pdf

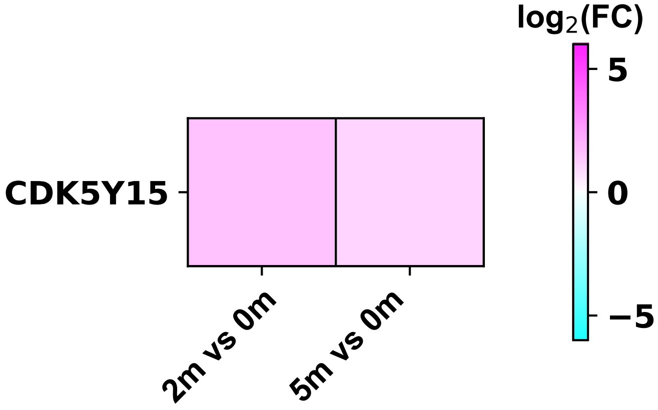

### cdk6_$^{y13}$_foldchange_m.pdf

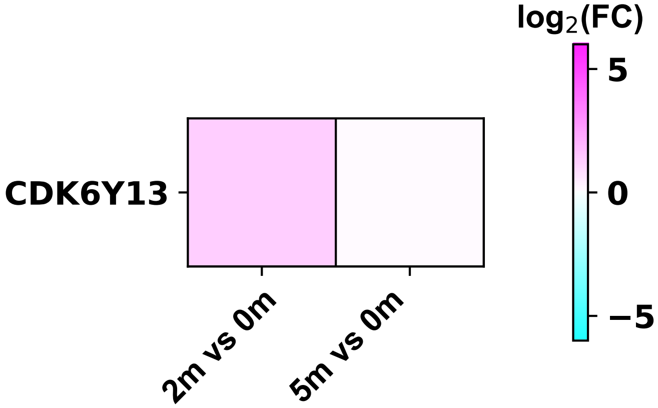

### cdk6_$^{y24}$_foldchange_m.pdf

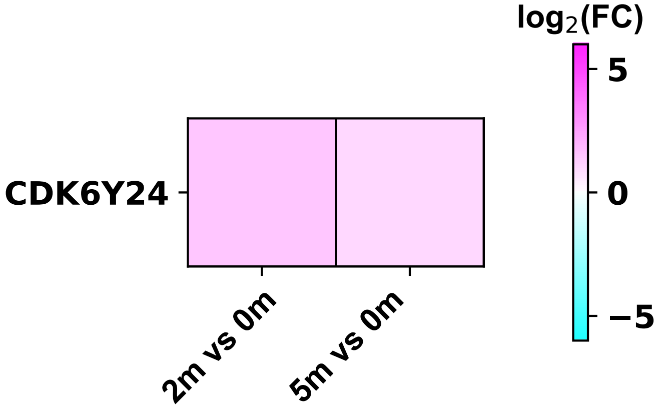

### cdk16_$^{y176}$_foldchange_m.pdf

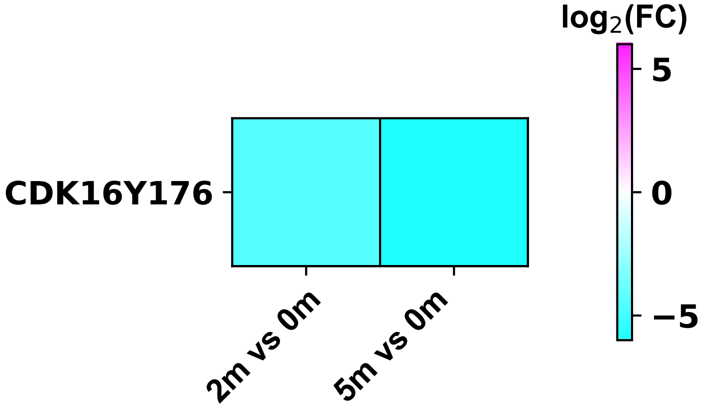

### cdkl5_$^{y171}$_foldchange_m.pdf

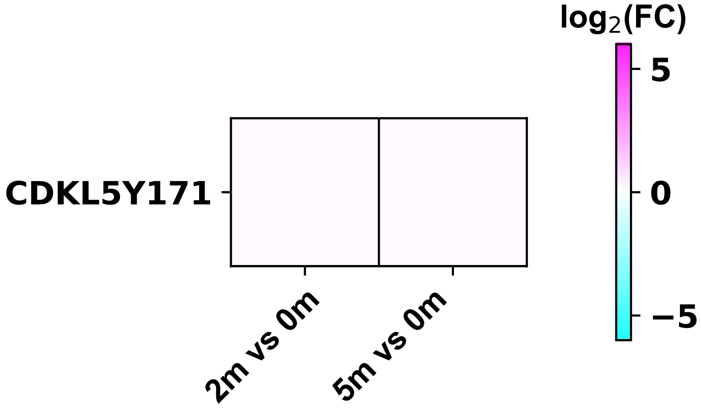

### cdv3_$^{y190}$_foldchange_m.pdf

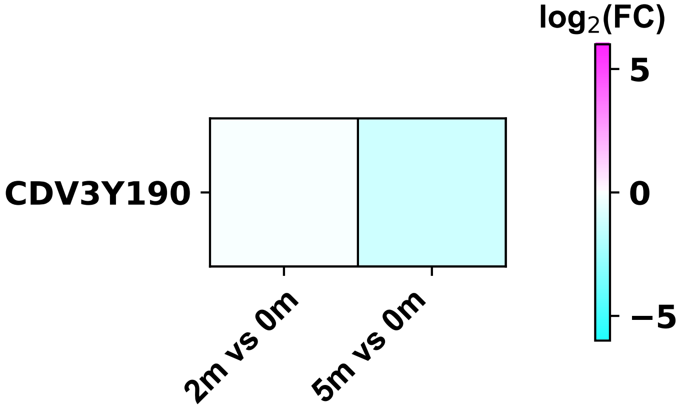

### cdv3_$^{y244}$_foldchange_m.pdf

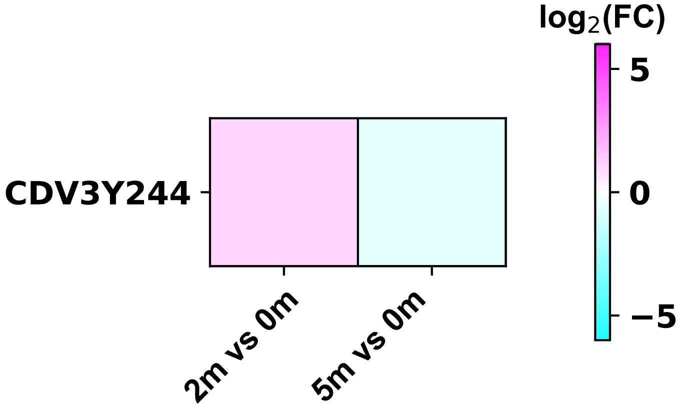

### cfl1_$^{y68}$_foldchange_m.pdf

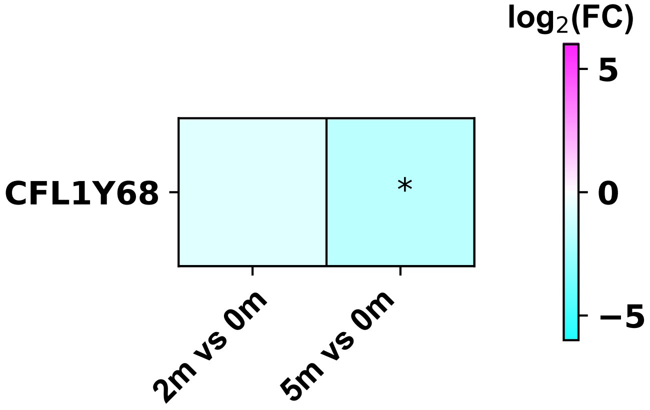

### cfl1_$^{y140}$_foldchange_m.pdf

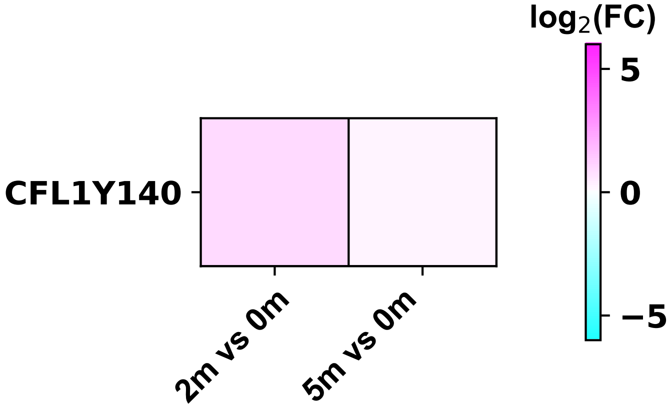

### DISEASE-PSP_0.pdf

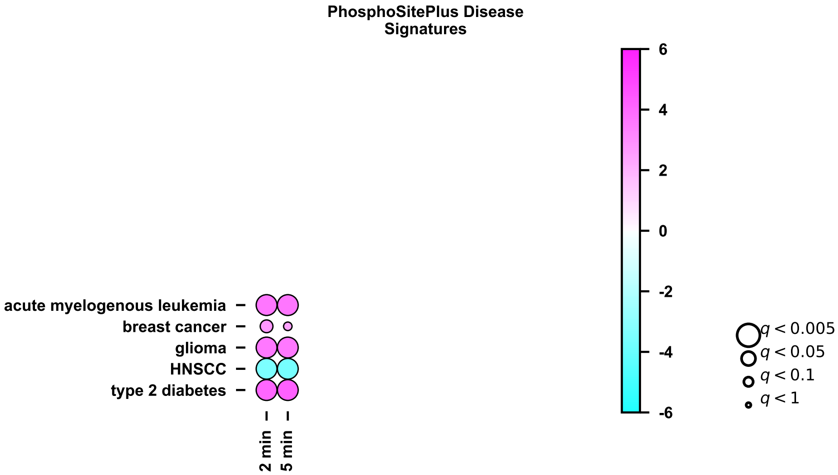

### KINASE-iKiP_0.pdf

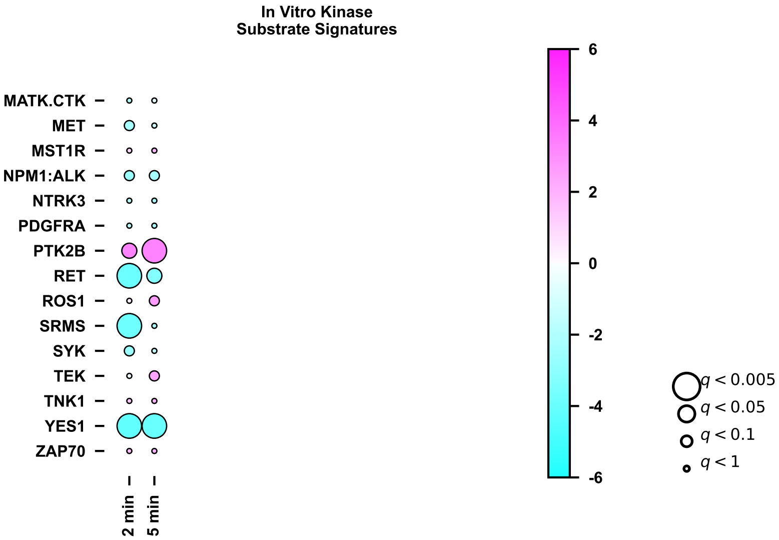

### KINASE-iKiP_1.pdf

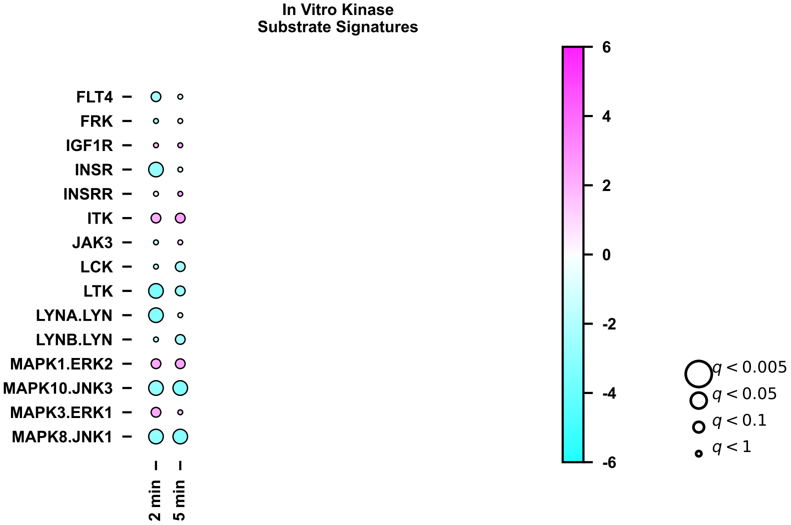

### KINASE-iKiP_2.pdf

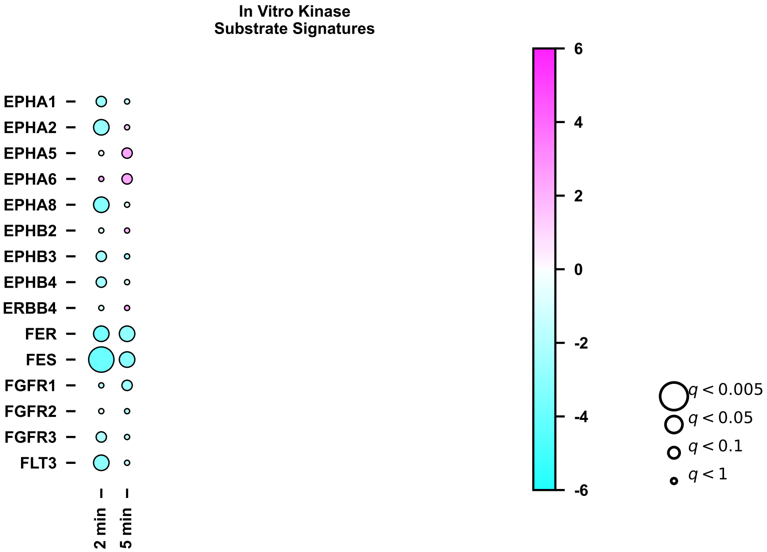

### KINASE-iKiP_3.pdf

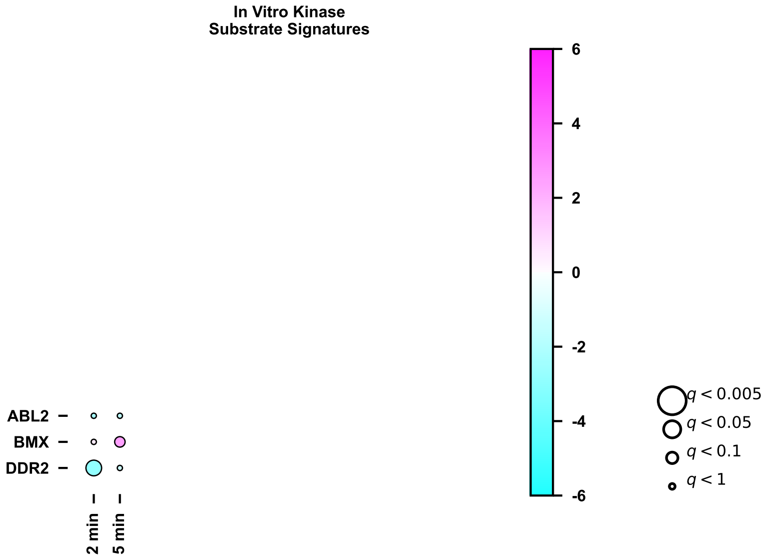

### KINASE-PSP_0.pdf

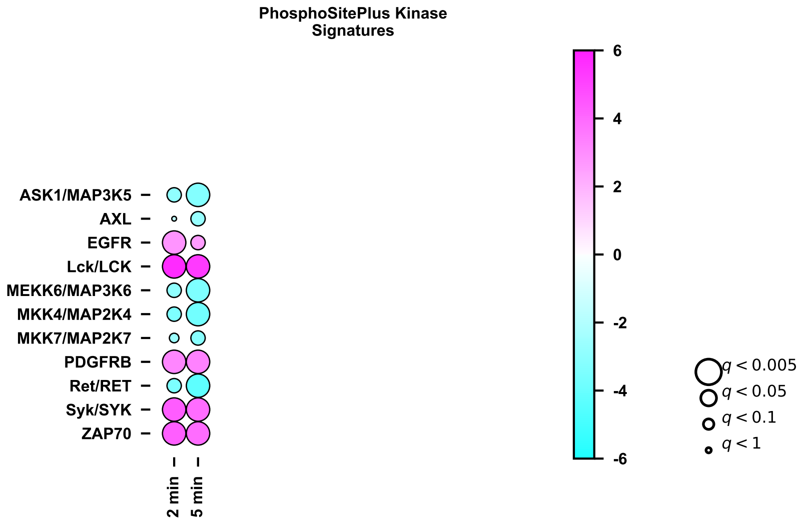

### PATH-BI_0.pdf

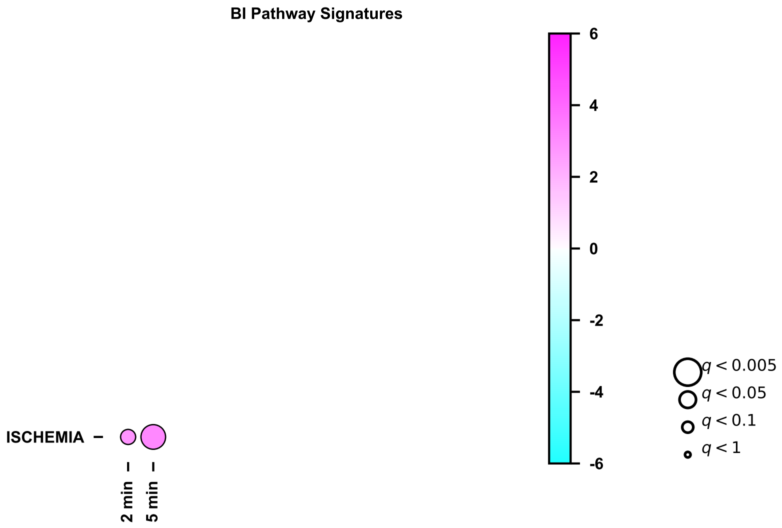

### PATH-NP_0.pdf

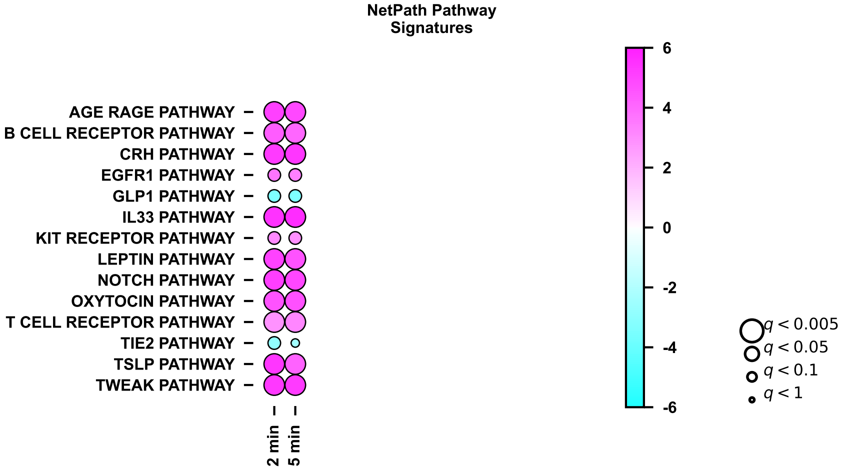

### PATH-WP_0.pdf

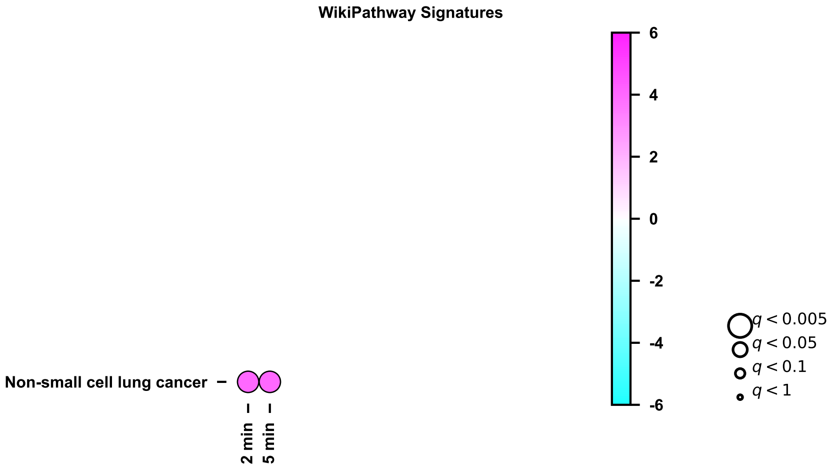

### PERT-P100-DIA2_0.pdf

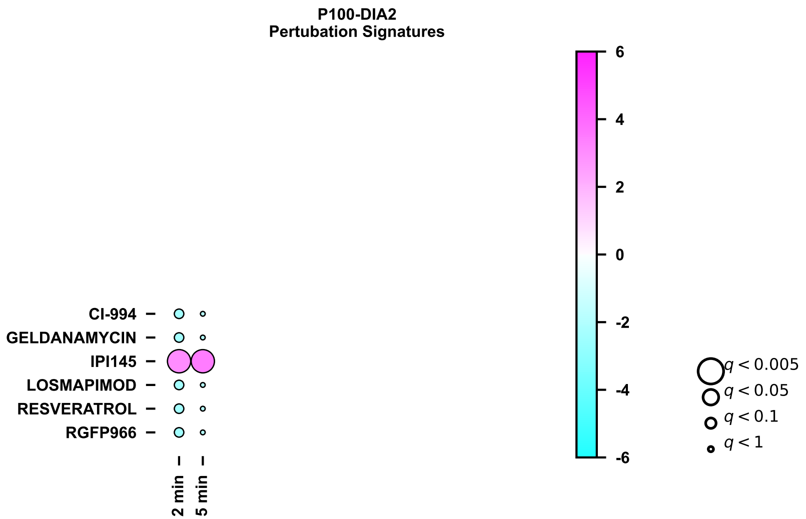

### PERT-PSP_0.pdf

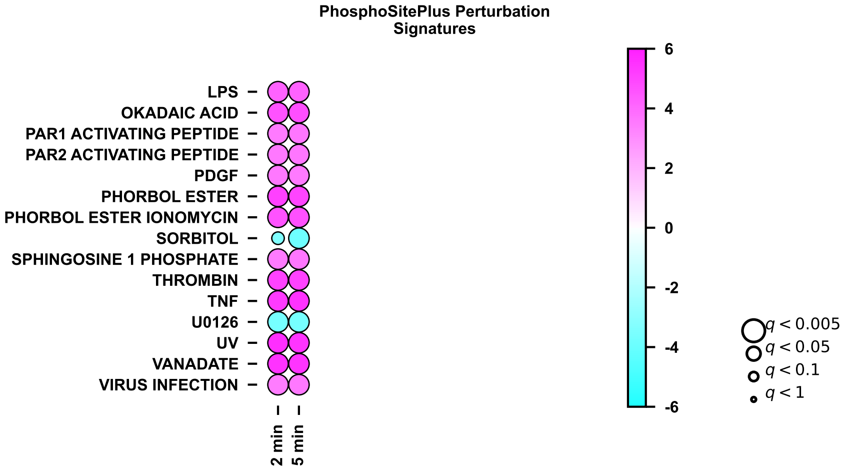

### PERT-PSP_1.pdf

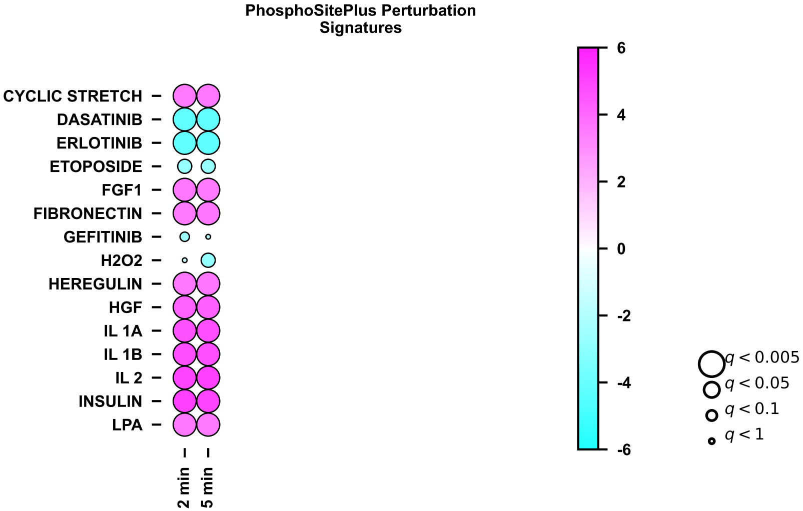

### PERT-PSP_2.pdf

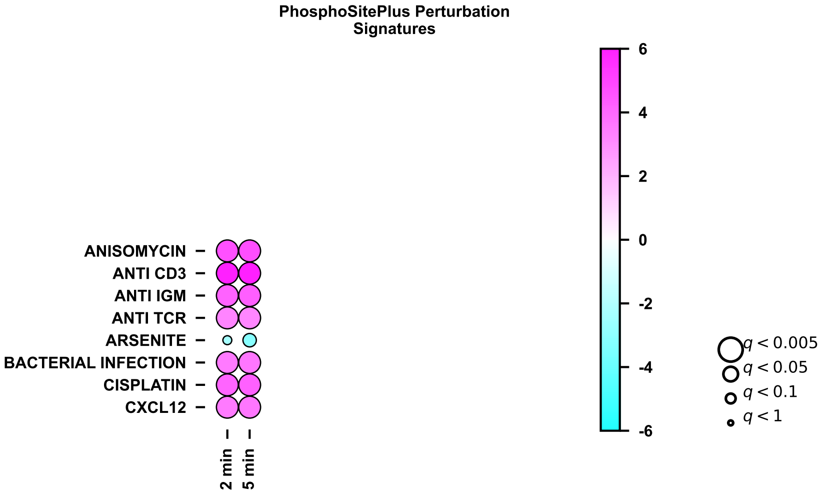
