## Supplemental Figures for "CSF1R-CAR T cells induce CSF1R signalling and promote cancer cell growth"

**Abstract**:

Chimeric antigen receptor (CAR) T cells have transformed the landscape of cancer therapy and demonstrate unprecedented success in treating relapsed/refractory blood cancers. The mechanism underlaying the interactions and responses of CAR T cells and their targets remain incompletely understood. Previous studies focus on the activation of CAR T cells and attempt to optimise CAR design to increase efficacy, meanwhile ignoring tumours and their responses to CAR ligation. Here, we evaluate the signalling capacity of a second generation CSF1-targeted CSF1R CAR compared with a scFv-targeted CD19-CAR using a SILAC co-culture approach coupled with phosphotyrosine (pY) enrichment and LC-MS/MS. We show that ligation of CSF1R-expressing THP1 cells with CSF1R-CAR T cells induces CSF1R-like signalling in THP1 cells, whereas no target cell signalling response is observed after CD19-CAR/Raji B cell ligation. Using small molecule inhibitors of Lck, actin polymerisation, and CSF1R, we find that CAR-induced CSF1R signalling in THP1 cells depends exclusively on CSF1R kinase activity with no participation from T cell activation. Consistently, CSF1R-CAR T cells promote THP1 growth at low effector-to-target (E:T) ratios but prevent THP1 growth at high E:T ratios. Our data provide evidence for an unintended consequence of CARs; CAR-induced signalling in cancer cells. These data may have broad implications for the choice of CAR antigen for optimal clinical efficacy.

**One Sentence Summary:** CSF1R-CAR activates intracellular signalling cascades in THP1 cells, which promote THP1 cell growth.

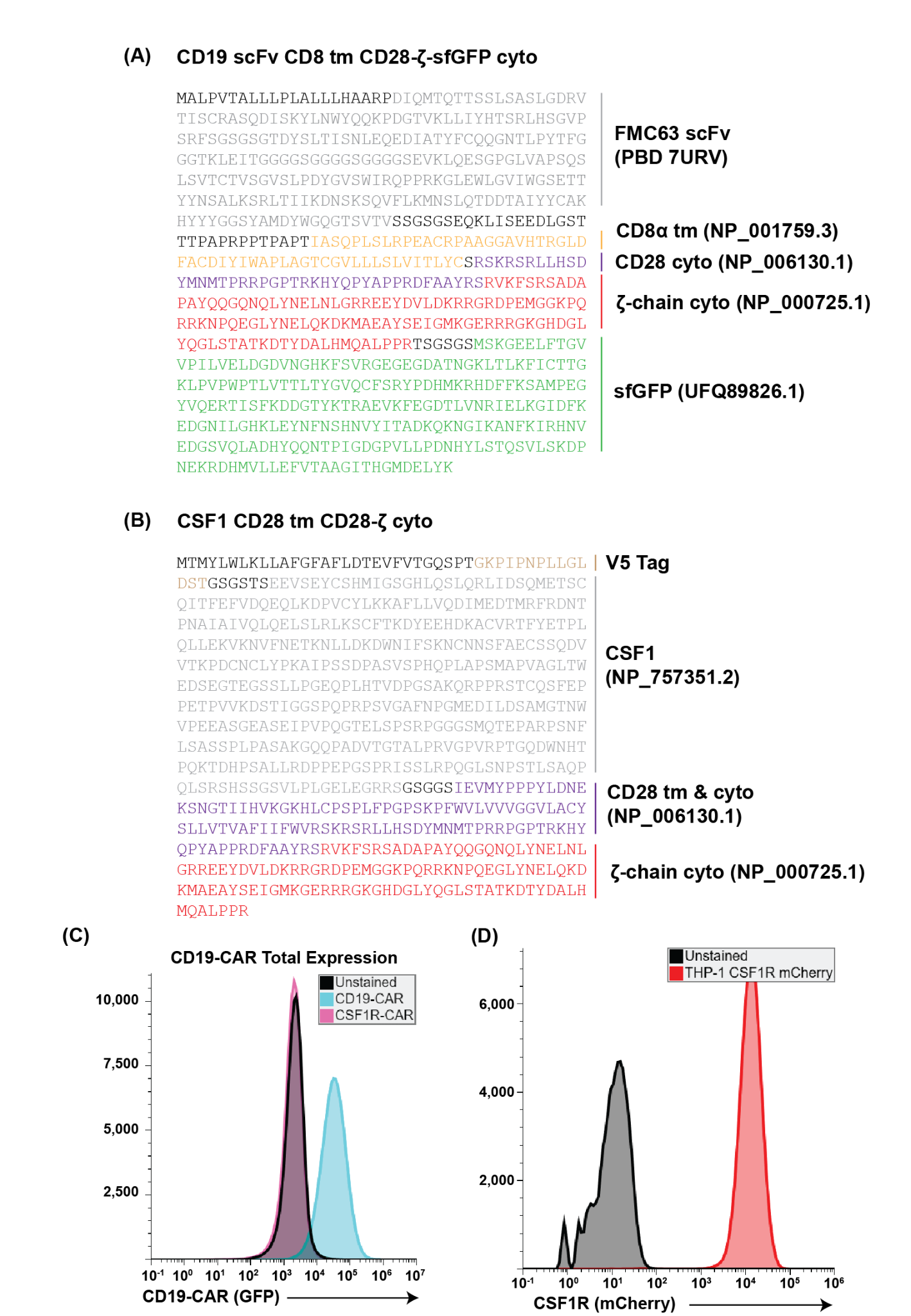

**Supporting Figure 1:** Protein sequences for (A) CD19-CAR and (B) CSF1R-CAR. Colours represent areas of perfect alignment with the proteins shown to the right. (C) GFP Flow cytometry showing total CD19-CAR expression. (D) Flow cytometry evaluating mCherry abundance in THP1 cells overexpressing CSF1R mCherry.

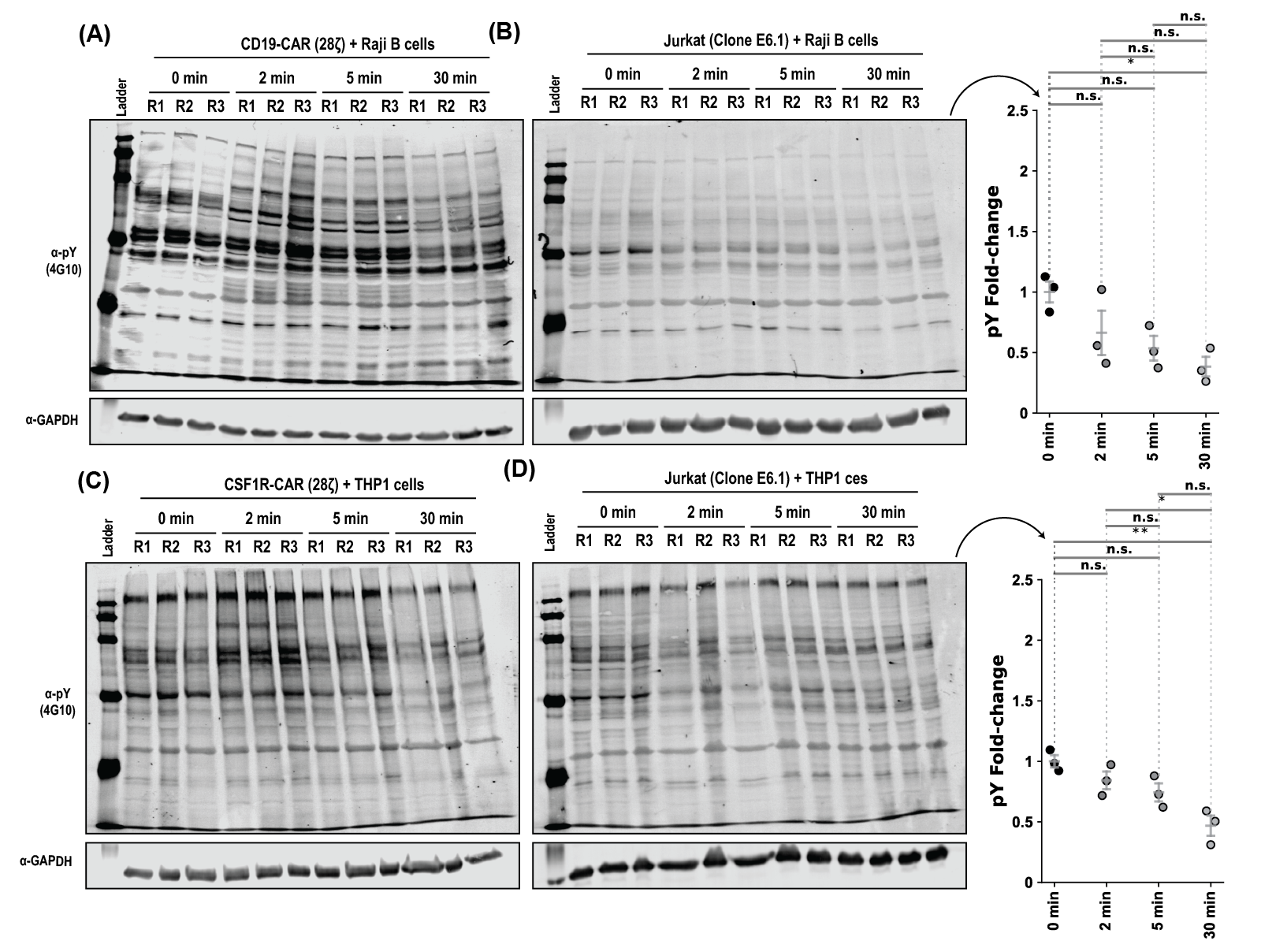

**Supporting Figure 2:** CSF1R-CAR/THP1 co-cultures exhibit altered pY induction compared with CD19-CAR/Raji co-cultures, while non-CAR Jurkat T cells elicit no pY responses. Total phosphotyrosine (pY) Western blots on co-cultures of (A) CD19-CAR and Raji B cells, (B) Jurkat (clone E6.1) and Raji B cells, (C) CSF1R-CAR and THP1 monocytes, and (D) Jurkat (clone E6.1) and THP1 monocytes. Total pY quantification, normalised to GAPDH, is displayed to the right of (B) and (D). * p < 0.05, ** p < 0.01, *** p < 0.001 (Fisher LSD with the Holm-Sidak FWER adjustment). Quantification for (A) and (C) is shown in Figure 1.

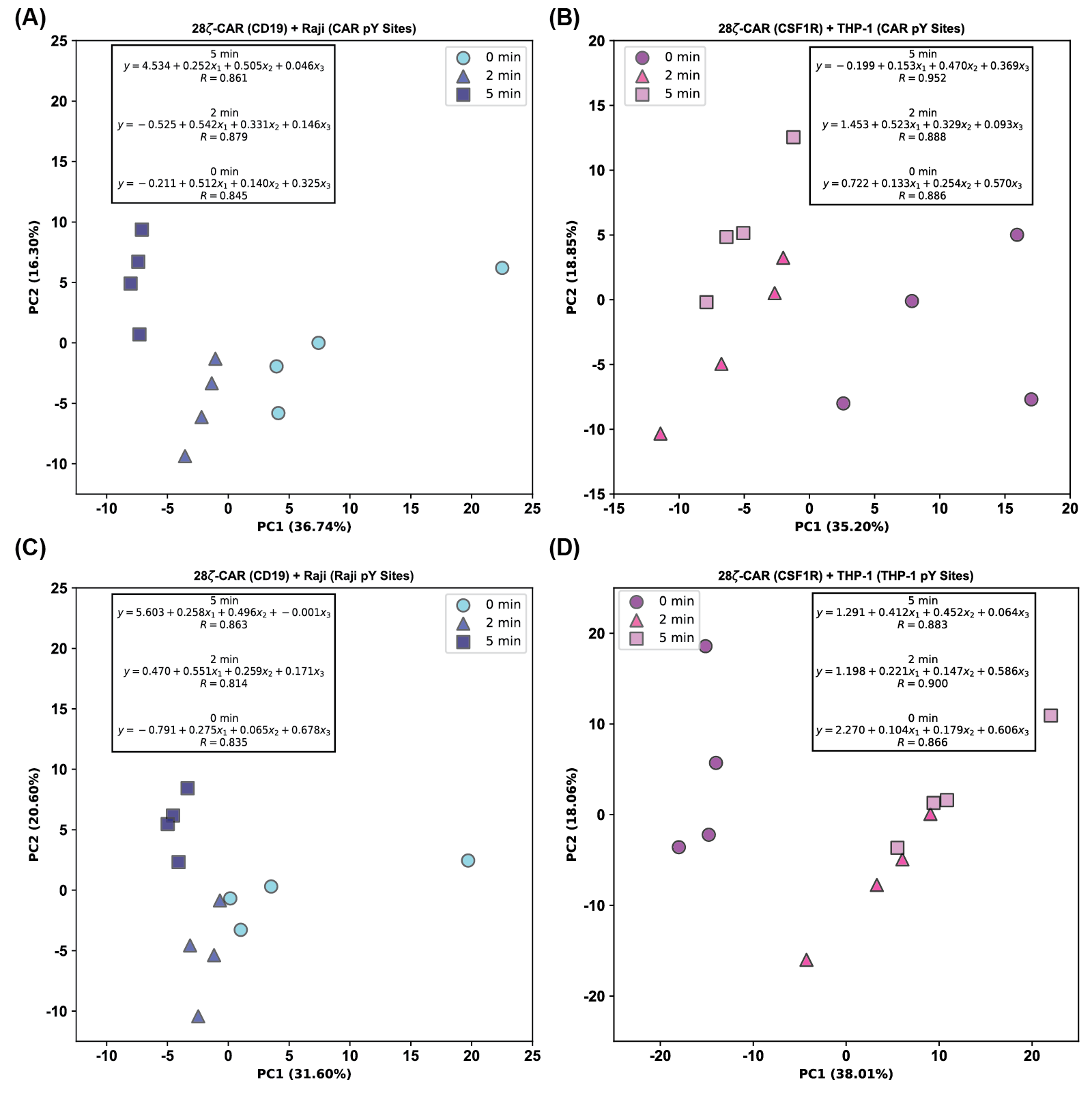

**Supporting Figure 3:** Reproducibility and clustering of pY proteomics data. (A) Principal component analysis of CD19-CAR T cell samples. Inset shows the results of multiple regression, including the line of best fit and the coefficient of determination. (B), (C), and (D) as in (A) except for CSF1R-CAR T cell, Raji cell, and THP1 monocytes, respectively.

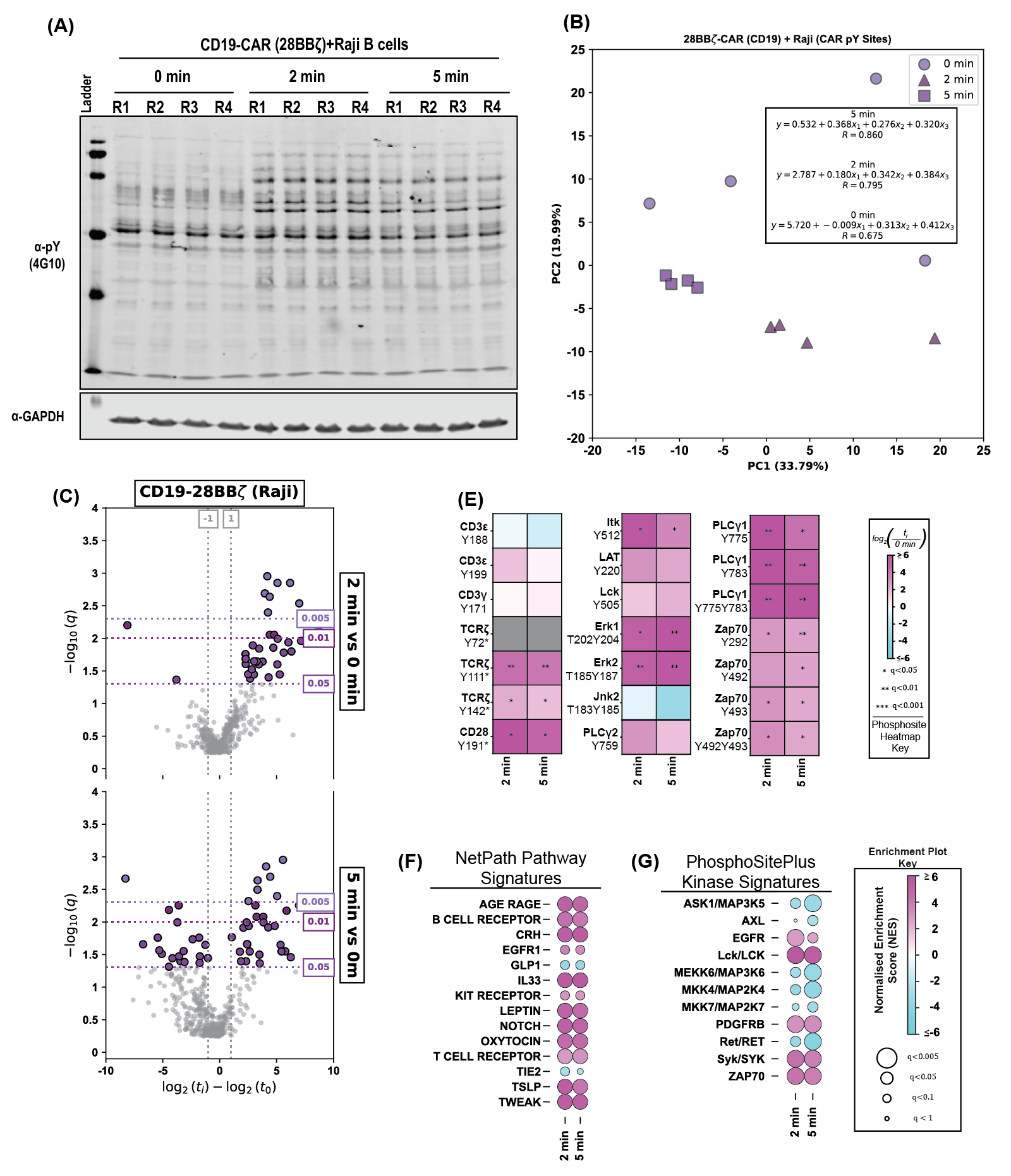

**Supporting Figure 4:** A third generation CD19-CAR with CD28 and 4-1BB costimulation shows T cell signalling engagement comparable to the second generation CD19-28ζ-CAR. (A) Principal component analysis of CD19-28bbζ-CAR T cell samples. Inset shows the results of multiple regression, including the line of best fit and the coefficient of determination. (B) Volcano plot analysis of pY sites observed in CD19-28bbζ-CAR T cells during co-culture with Raji cells after 2- and 5-minutes. (C) Heatmaps showing quantification of individual phosphorylation sites observed in CD19-28bbζ-CAR T cells during co-culture with Raji cells. (D) PTM-SEA showing enrichment of CD19-CAR T cell PTM sites in NetPath Pathways. (E) PTM-SEA showing enrichment of CD19-CAR T cell PTM sites in the PhosphoSitePlus Kinase-signatures database.

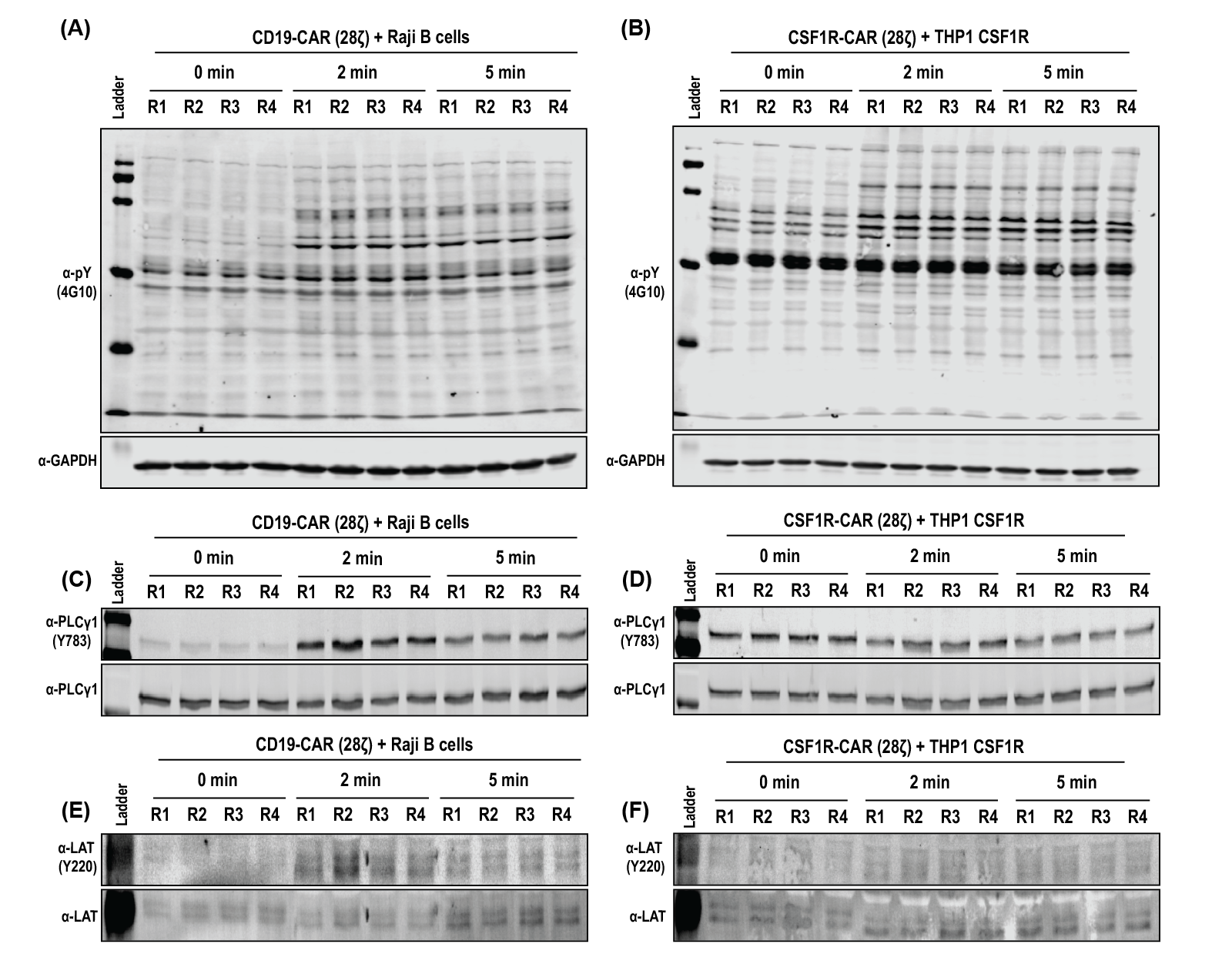

**Supporting Figure 5:** Western blot verification of T cell signalling proteins from our co-culture pY proteomics results. α-pY (clone 4G10) Western blots for (A) CD19-CAR/Raji and (B) CSF1R-CAR/THP1 co-culture proteomics samples. α-PLCγ1 Y783 Western blots for (C) CD19-CAR/Raji and (D) CSF1R-CAR/THP1 co-culture proteomics samples. α-LAT Y220 Western blots for (E) CD19-CAR/Raji and (F) CSF1R-CAR/THP1 co-culture proteomics samples. Quantification for these Western blots are presented in Figure 2.

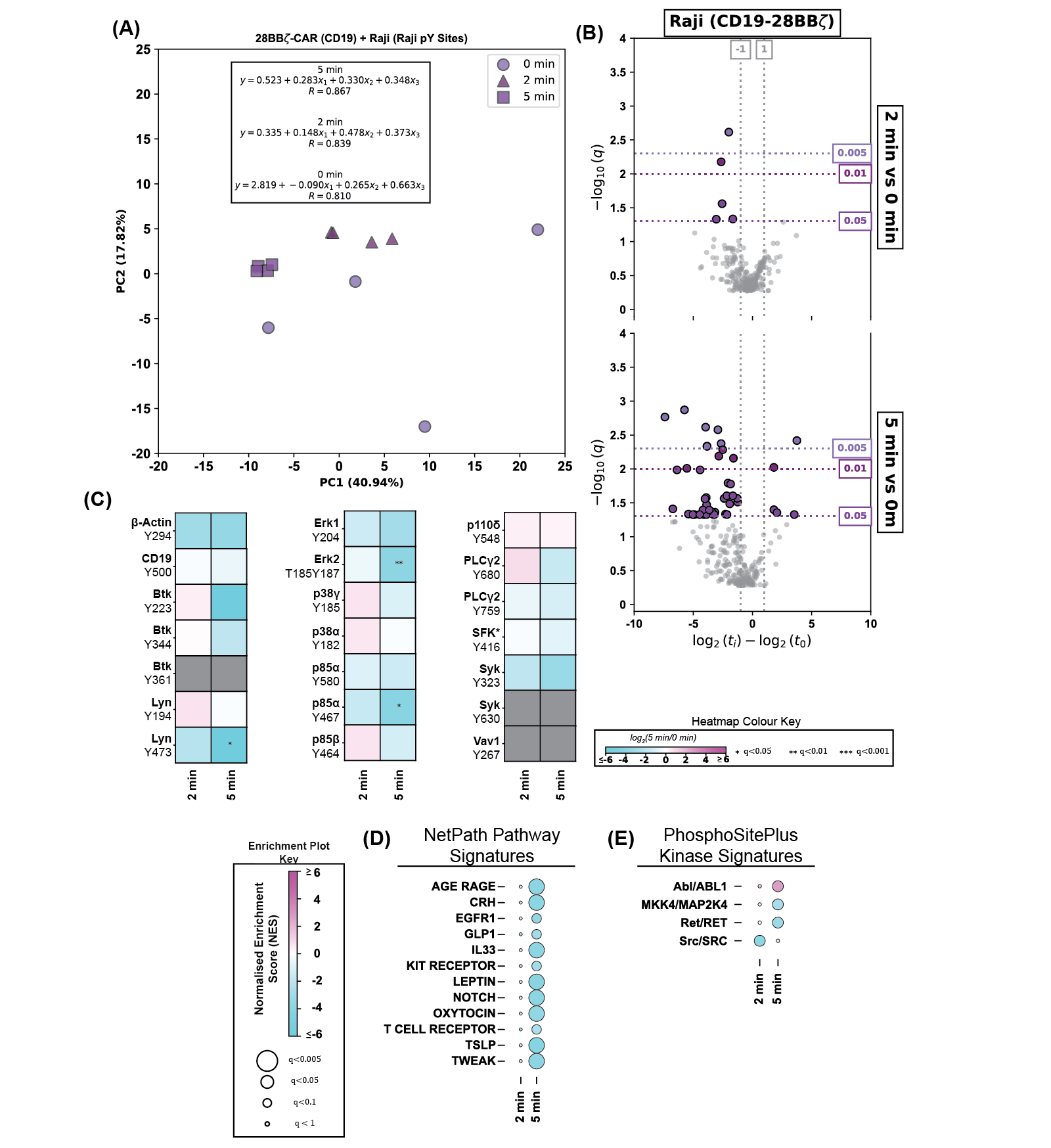

**Supporting Figure 6:** Raji cells show minimal signal induction after co-culture with CD19-28bbζ-CAR T cells (A) Principal component analysis of Raji cell (28bbζ) samples. Inset shows the results of multiple regression, including the line of best fit and the coefficient of determination. (B) Volcano plot analysis of pY sites observed in Raji cells during co-culture with CD19-28bbζ-CAR T cells after 2- and 5-minutes. (C) Heatmaps showing quantification of individual phosphorylation sites observed in Raji cells during co-culture with CD19-28bbζ-CAR T cells. (D) PTM-SEA showing enrichment of Raji cells PTM sites in NetPath Pathways. (E) PTM-SEA showing enrichment of Raji cells PTM sites in the PhosphoSitePlus Kinase-signatures database.

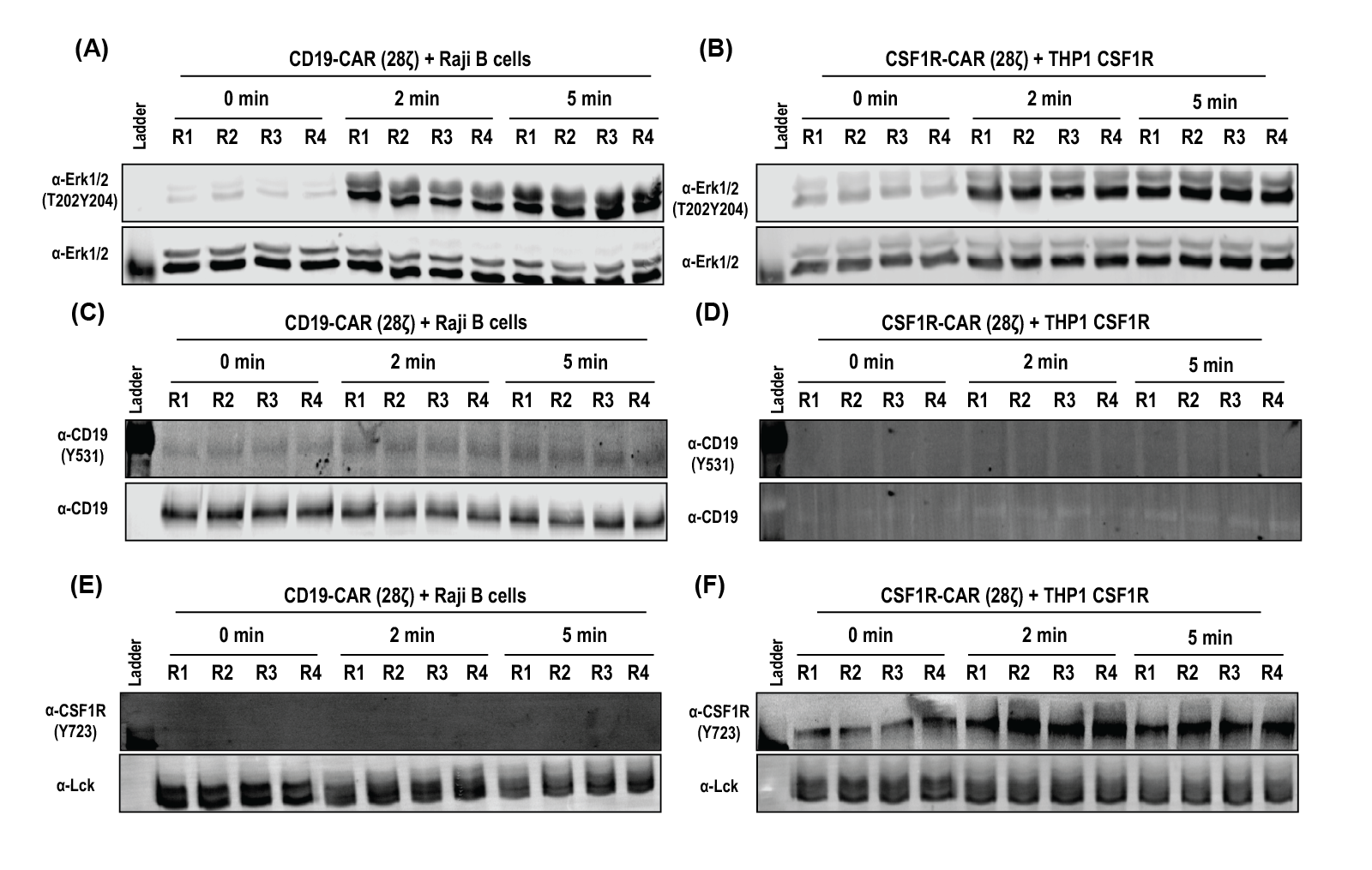

**Supporting Figure 7:** Western blot verification of target cell signalling proteins from our co-culture pY proteomics results. α-Erk T202Y204 Western blots for (A) CD19-CAR/Raji and (B) CSF1R-CAR/THP1 co-culture proteomics samples. α-CD19 Y531 Western blots for (C) CD19-CAR/Raji and (D) CSF1R-CAR/THP1 co-culture proteomics samples. α-CSF1R Y723 Western blots for (E) CD19-CAR/Raji and (F) CSF1R-CAR/THP1 co-culture proteomics samples. Quantification of these Western blots are presented in Figure 3.

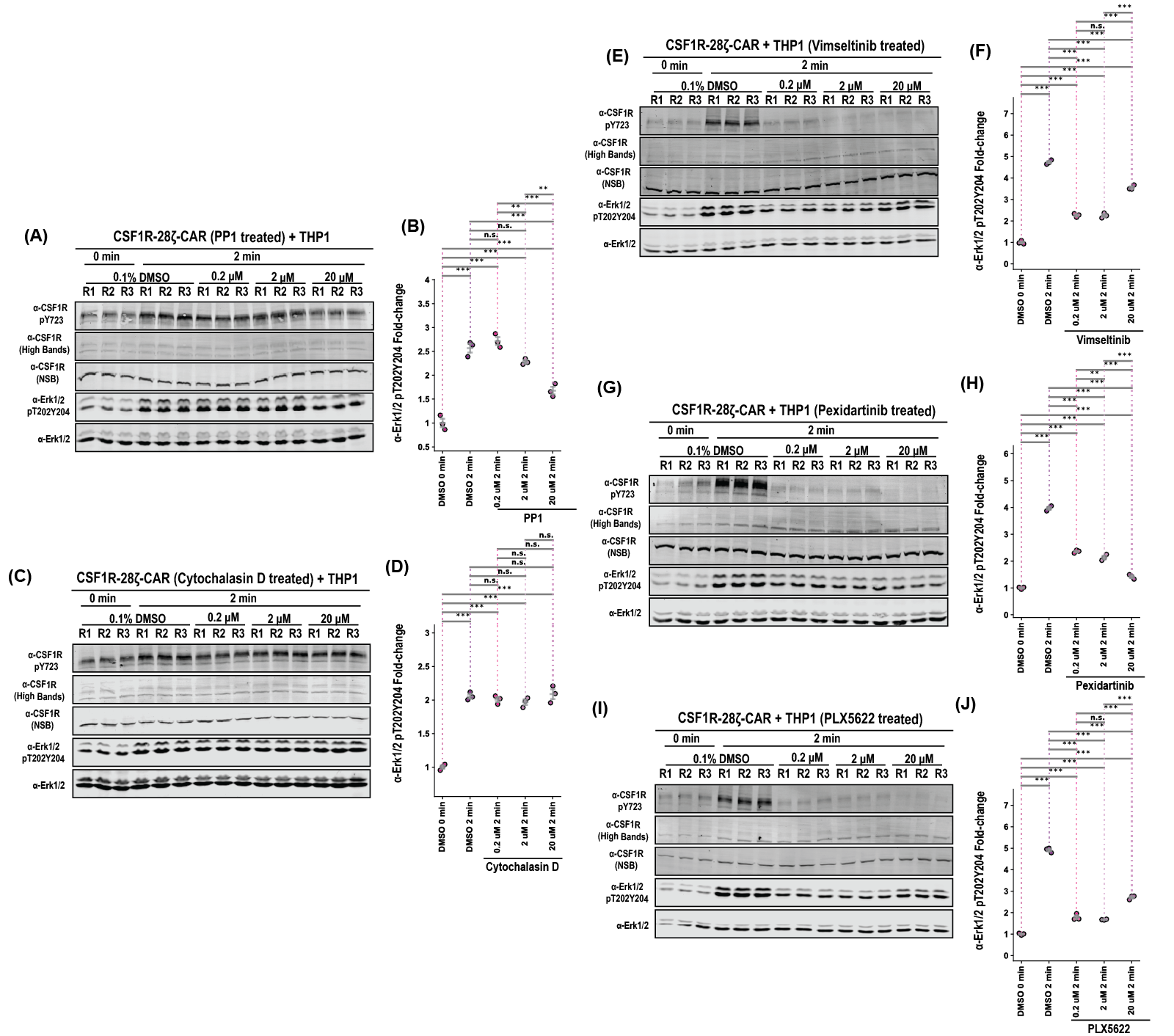

**Supporting Figure 8:** CSF1R kinase inhibitors ablate CSF1R^Y723^ induction and significantly reduce phosphorylation of Erk1^T202Y204^ and Erk2^T185Y187^. (A) Western blot analysis of α-CSF1R^Y723^ and α-Erk^T202Y204^ in PP1-treated CSF1R-CAR T cells after co-culture with THP1 monocytes, with (B) α-Erk^T202Y204^ quantification. (C-D) As in (A-B), except for Cytochalasin D-treated CSF1R-CAR T cells. (E) Western blot analysis of α-CSF1R^Y723^ and α-Erk^T202Y204^ in Vimseltinib treated THP1 monocytes after co-culture with CSF1R-CAR T cells, with (F) α-Erk^T202Y204^ quantification. (G-H) As in (E-F), except for Pexidartinib-treated THP1 monocytes. (I-J) As in (E-F), except for PLX 5622-treated THP1 monocytes. Statistical significance is determined by a two-way ANOVA with Fisher’s least significant difference and the Holm-Sidak correction for multiple comparisons. * : p < 0.05, ** : p<0.01, *** : p<0.001.

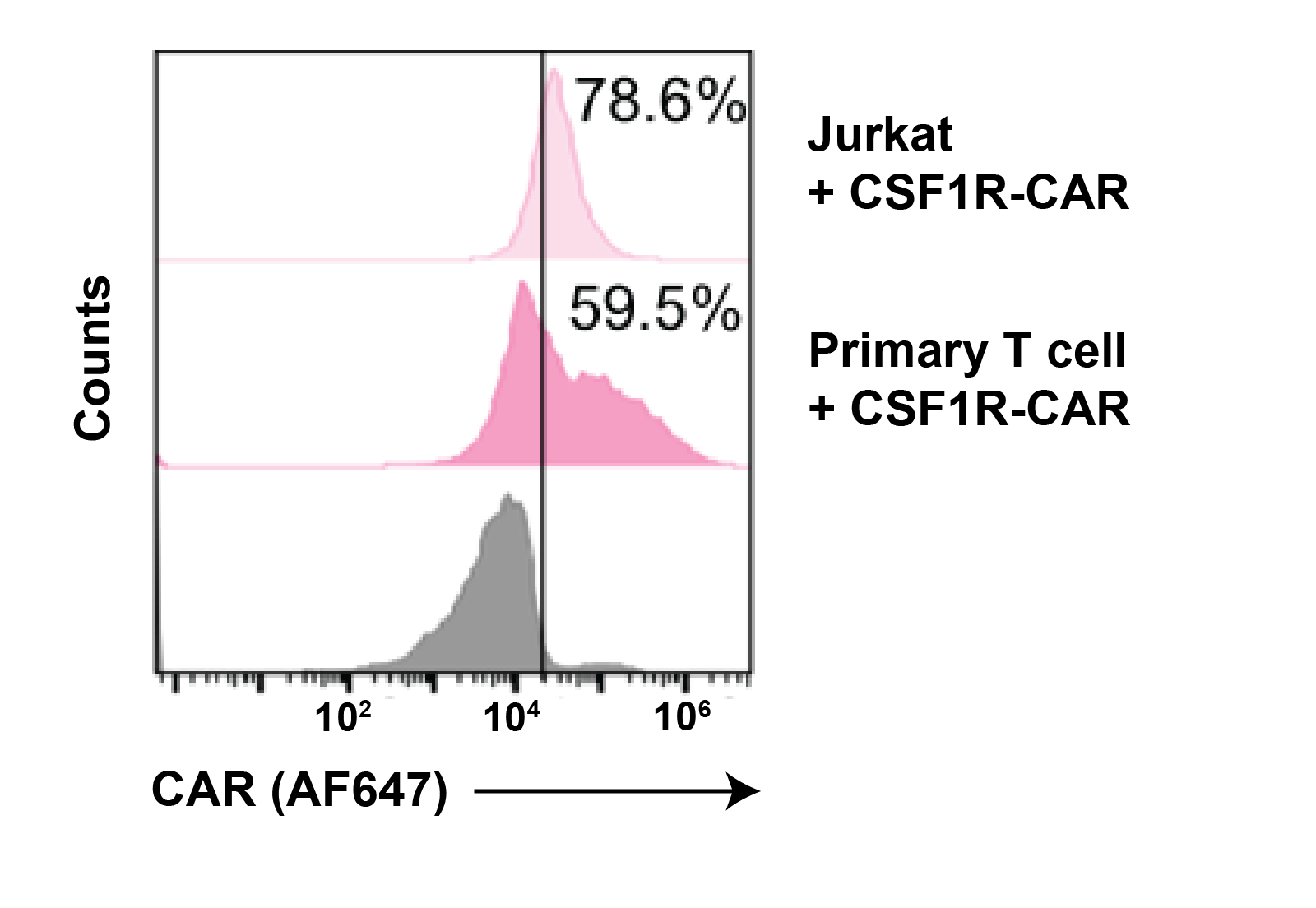

**Supporting Figure 9:** Expression of CAR-Jurkats and CAR-Primary T cells used for luciferase co-culture lysis assays.
